# Worth the effort? A practical examination of random effects in hidden Markov models for animal telemetry data

**DOI:** 10.1101/2020.07.10.196410

**Authors:** Brett T. McClintock

## Abstract

1. Hidden Markov models (HMMs) that include individual-level random effects have recently been promoted for inferring animal movement behaviour from biotelemetry data. These “mixed HMMs” come at significant cost in terms of implementation and computation, and discrete random effects have been advocated as a practical alternative to more computationally-intensive continuous random effects. However, the performance of mixed HMMs has not yet been sufficiently explored to justify their widespread adoption, and there is currently little guidance for practitioners weighing the costs and benefits of mixed HMMs for a particular research objective.
2. I performed an extensive simulation study comparing the performance of a suite of fixed and random effect models for individual heterogeneity in the hidden state process of a 2-state HMM. I focused on sampling scenarios more typical of telemetry studies, which often consist of relatively long time series (30 – 250 observations per animal) for relatively few individuals (5 – 100 animals).
3. I generally found mixed HMMs did not improve state assignment relative to standard HMMs. Reliable estimation of random effects required larger sample sizes than are often feasible in telemetry studies. Continuous random effect models performed reasonably well with data generated under discrete random effects, but not vice versa. Random effects accounting for unexplained individual variation can improve estimation of state transition probabilities and measurable covariate effects, but discrete random effects can be a relatively poor (and potentially misleading) approximation for continuous variation.
4. When weighing the costs and benefits of mixed HMMs, three important considerations are study objectives, sample size, and model complexity. HMM applications often focus on state assignment with little emphasis on heterogeneity in state transition probabilities, in which case random effects in the hidden state process simply may not be worth the additional effort. However, if explaining variation in state transition probabilities is a primary objective and sufficient explanatory covariates are not available, then random effects are worth pursuing as a more parsimonious alternative to individual fixed effects.
5. To help put my findings in context and illustrate some potential challenges that practitioners may encounter when applying mixed HMMs, I revisit a previous analysis of long-finned pilot whale biotelemetry data.

## 1 Introduction

Hidden Markov models (HMMs) are used extensively in ecology for inferences about unobservable state processes from sequential (e.g. time series) data (Zucchini *et al*. 2016;McClintock *et al*. 2020). Some of the most widely used HMMs in population ecology include capture-recapture (e.g. Pradel 2005), species occurrence (e.g. Gimenez *et al*. 2014), and animal movement (e.g. Franke *et al*. 2004) models. As recent advances in animal-borne biologging technology have permitted the collection of detailed location and biotelemetry data (e.g. Cooke *et al*. 2004), HMMs for inferring animal movement behaviour have become particularly popular (e.g. Morales *et al*. 2004; Jonsen *et al*. 2005; Patterson *et al*. 2009; Langrock *et al*. 2012; McClintock *et al*. 2012). This has been bolstered by user-friendly software specifically tailored to HMMs for these data (Michelot *et al*. 2016; McClintock & Michelot 2018).

While animal movement HMMs were originally formulated as relatively simple 2-state (e.g. “foraging” and “transit”) models describing steps and turns between successive locations (e.g. Franke *et al*. 2004; Morales *et al*. 2004), they have since become much more complicated by incorporating location measurement error (e.g. Jonsen *et al*. 2005), > 2 movement behaviour states (e.g. Michelot *et al*. 2017; Pirotta *et al*. 2018), additional biotelemetry data streams (e.g. DeRuiter *et al*. 2017; Isojunno *et al*. 2017), and “mixed HMMs” including individual-level random effects (e.g. Langrock *et al*. 2012;Schliehe-Diecks *et al*. 2012; McClintock *et al*. 2013; McKellar *et al*. 2015). While all of these advances bring the potential for new and exciting inferences about animal movement behaviour, they also pose various challenges (e.g. Patterson *et al*. 2017; Pohle*et al*. 2017). The inferential benefits of accounting for measurement error or including additional data streams to characterize > 2 behavioural states can justify this added complexity (e.g. Bradshaw *et al*. 2007; McClintock 2017), yet the general benefits of mixed HMMs that include individual random effects are less well understood.

There is evidence for the benefits of individual random effects on the (conditional) observation process of HMMs in other contexts (e.g. Altman 2007; Rueda *et al*. 2013), but there is surprisingly little evidence for benefits on the hidden state process. In a simulation study, Altman (2007) concluded for their case that there was “far more information about the parameters associated with the conditional model than those associated with the hidden model” and that mixed HMMs allowing for individual differences in the hidden state process “may explain very little additional variation in the observed data and, hence, may not be worthwhile from a statistical standpoint.” Yet understanding individual heterogeneity in behaviour or life history strategies is a fundamental component of ecology and evolution (e.g. Johnson *et al*. 1986; Cam *et al*. 2002; Réale *et al*. 2007; Revilla & Wiegand 2008; Gimenez *et al*. 2018), and accounting for individual variation in the hidden state process is clearly worthwhile for this purpose. Most animal movement mixed HMM applications have employed random effects on the hidden state process more as a statistical tool to “mop up” unexplained variation and improve goodness-of-fit, with little attempt to interpret the mechanisms or implications of this variation (e.g. McKellar *et al*. 2015; Towner *et al*. 2016; DeRuiter *et al*. 2017). Part of the reason for this may be that, unlike effect sizes for explanatory covariates (e.g. age, sex, weight), generic random effects are difficult to interpret (e.g. Altman 2007; Gimenez *et al*. 2018), particularly in biological terms across free-ranging telemetered individuals, each typically with different deployment lengths and being observed in different environmental and behavioural contexts (e.g. Towner *et al*. 2016; DeRuiter *et al*. 2017). Nevertheless, failing to properly account for individual variation could be detrimental to the estimation of effect sizes for any explanatory covariates on the hidden state process (e.g. DeRuiter *et al*. 2017).

In their discussion of the benefits of individual random effects on the hidden state process, DeRuiter *et al*. (2017) claim “it is easy to argue that random effects to account for individual variation are a key component of animal behaviour models.” I certainly do not disagree with this, but the empirical performance of these complex models has not yet been sufficiently explored to justify their widespread adoption (*sensu* Hodges 2019), and there is currently little guidance for practitioners to determine when and how they should pursue random effects. This would all be relatively moot if mixed HMMs were easy for practitioners to implement, but they typically are not. Mixed HMMs have historically required custom-coded model fitting algorithms at significant computational cost (e.g. Altman 2007; Maruotti & Rydén 2009; Langrock *et al*. 2012;Schliehe-Diecks *et al*. 2012; McKellar *et al*. 2015; Towner *et al*. 2016) and can be very challenging to reliably fit to time series of animal biotelemetry data (e.g. DeRuiter *et al*. 2017; Isojunno *et al*. 2017). Furthermore, unlike other ecological applications of mixed HMMs that typically include relatively many individuals (e.g. capture-recapture; Burnham & White 2002; Gimenez & Choquet 2010) or sites (e.g. occupancy; Gimenez *et al*. 2014) and relatively short time series, animal-borne biotelemetry studies typically include relatively few individuals (e.g. due to financial and logistical constraints) and relatively long time series of unequal length for each individual (e.g. due to variable battery life, tag loss, and mortality). Little is currently known about how well mixed HMMs perform under these sampling conditions.

These challenges and uncertainties prompted me to investigate the benefits of accounting for individual variation in the hidden state process of HMMs that are frequently used for inferring animal movement behaviour from biotelemetry data. Using extensive simulation and a case study, my goal is to provide some guidance to aid practitioners in weighing the costs and benefits of mixed HMMs for a particular research objective. I first describe several of the most common HMM formulations and inferential procedures that account for individual variation. I then present a large-scale simulation study evaluating the performance of these various approaches in terms of hidden state estimation, parameter estimation, and detection of individual variation by standard information-theoretic model selection criteria. To help put my findings in context and illustrate some potential challenges that practitioners may encounter when applying mixed HMMs, I then revisit an analysis of long-finned pilot whale biotelemetry data originally performed by Isojunno *et al*. (2017). Finally, I discuss the implications of my findings in establishing some considerations for practitioners contemplating the inclusion of individual random effects in their own analyses.

## 2 Individual-level effects in HMMs

### 2.1 Model formulations

Under “complete pooling”, standard HMMs for *M* individual time series of length *T_m_* (*m* = 1,…, *M*) assume no individual effects on parameters (i.e. a common set of parameters is shared among the *M* individuals). Assuming independence between individuals, the likelihood function for this “null” model with *N* hidden states can be succinctly expressed using the forward algorithm:

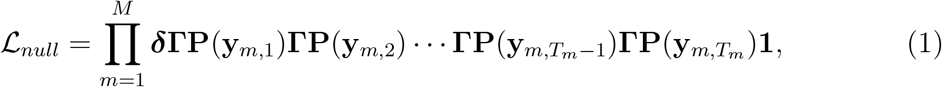

where ***δ*** = (*δ*_1_,…, *δ_N_*) is a row vector of initial state probabilities 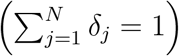, **Γ** = (*γ_i,j_*) is a *N* × *N* state transition probability matrix with entries *γ_i,j_* corresponding to the probability of switching from state *i* at time *t* – 1 to state *j* at time *t* 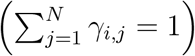, **P**(**y**_*m,t*_) = diag (*f*(**y**_*m,t*_ | *S_m,t_* = 1),…, *f*(**y**_*m,t*_ | *S_m,t_* = *N*)) is a *N* × *N* diagonal matrix with entries *f*(**y**_*m,t*_ | *S_m,t_* = *s*) corresponding to the (state-dependent) conditional probability density of observation **y**_*m,t*_ given the state *S_m,t_* ∈ {1,…, *N*} at time *t*, and 1 is a column vector of *N* ones (e.g. Zucchini *et al*. 2016). Here I assume all individuals share common state-dependent distribution parameters, a case where pooling collective movements across *M* individuals using a joint likelihood has been demonstrated to improve behavioural state assignment in animal movement HMMs (Jonsen 2016). When explanatory individual covariates (e.g. age, sex, weight) are available, the likelihood can be extended to accommodate individual variation attributable to these factors through link functions for the model parameters (e.g. McClintock & Michelot 2018).

Generic individual heterogeneity in HMMs is typically handled using individual-level fixed effects (termed “no pooling”; e.g. Patterson *et al*. 2009), discrete-valued random effects based on finite mixtures (e.g. Maruotti & Rydén 2009; McKellar *et al*. 2015;Towner *et al*. 2016; DeRuiter *et al*. 2017), or continuous-valued random effects (e.g. Altman 2007; Schliehe-Diecks *et al*. 2012). For individual fixed effects in the hidden state process, we have:

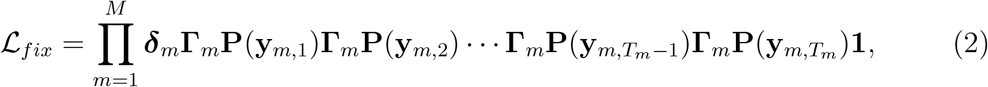

where ***δ***_*m*_ = (*δ*_*m*,1_,…, *δ_m,N_*) and **Γ**_*m*_ = (*γ_m,i,j_*) is a *N* × *N* state transition probability matrix with entries *γ_m,i,j_* corresponding to the probability of individual *m* switching from state *i* at time *t* – 1 to state *j* at time *t*. This model is highly parameterized with *M N*^2^ state transition probabilities, but it avoids any distributional assumptions about the individual effects.

For mixed HMMs with discrete-valued random effects, we have:

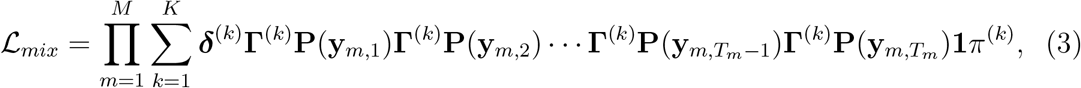

where *K* ∈ {1,…, *M* – 1} is the number of mixtures (typically chosen *a priori* or based on model selection criteria, e.g. DeRuiter *et al*. 2017), 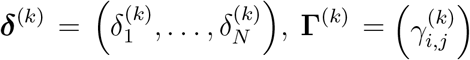 has entries 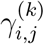 corresponding to the probability that an individual belonging to mixture *k* switches from state *i* at time *t* – 1 to state *j* at time *t*, and ***π*** = *π*^(1)^,…, *π*^(*K*)^ are the mixture probabilities 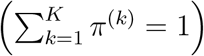. One would expect finite mixtures to be most appropriate for explaining individual variation attributable to unmeasured categorical covariates, but many such factors (e.g. age class, sex) can be measured in biotelemetry studies where individuals must be captured for tag deployment. It is less clear how appropriate finite mixtures are for “mopping up” unexplained continuous-valued variation, but they have recently been promoted for this purpose in animal movement HMMs (e.g. McKellar *et al*. 2015; Towner *et al*. 2016).

For mixed HMMs with continuous random effects, the simplest models typically assume independent and identically distributed Gaussian random effects 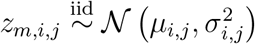 for *i* ≠ *j*, *z_m,i,j_* = 0 for *i* = *j*, and 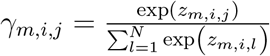, such that:

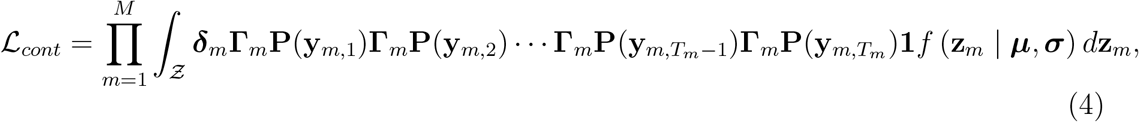

where 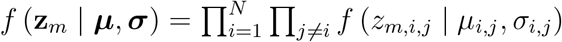 is the joint density of **z**_*m*_ = (*z_m,i,j_*)_*i*=*j*_ and 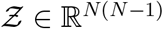 its support. One would expect continuous random effects to be more parsimonious than fixed effects (Eq. 2), but this comes at the cost of additional distributional assumptions and computational complexity.

### 2.2 Maximum likelihood inference

I focus on maximum likelihood (ML) inference because HMMs can be fitted relatively quickly using ML methods and thereby facilitate large-scale simulation experiments. Standard ML inference by direct numerical maximization of the likelihood is straight-forward in principle for the null (Eq. 1), fixed (Eq. 2), and finite mixture (Eq. 3) models (Patterson *et al*. 2009; McKellar *et al*. 2015; Zucchini *et al*. 2016), and there are R (R Core Team 2020) packages specifically designed for fitting these HMMs (e.g. Visser & Speekenbrink 2010; McClintock & Michelot 2018).

The integral in the continuous random effects model (Eq. 4) poses additional challenges that have historically made fitting by ML largely intractable (e.g. Altman 2007;Schliehe-Diecks *et al*. 2012; DeRuiter *et al*. 2017). The dimension of the integral is *N* (*N* – 1), which for *N* > 2 is generally not feasible for direct maximization of the likelihood using standard numerical integration techniques such as Gaussian quadrature (e.g. Abramowitz & Stegun 1964). This largely explains why custom Monte Carlo expectation-maximization (e.g. Altman 2007) and Bayesian Markov chain Monte Carlo (e.g. McClintock *et al*. 2013) algorithms have often been employed for fitting HMMs with continuous random effects.

Framed as a compromise between null and fixed effect models, Burnham & White (2002) proposed an approximate but computationally simple method for continuous random effect estimation based solely on maximum likelihood estimates from fixed effect models. Little used outside of capture-recapture applications, their approach was originally developed for temporal random effects in the Cormack-Jolly-Seber model of survival and, to my knowledge, has not been applied or investigated in contexts other than capture-recapture. However, standard open population capture-recapture models are simply special cases of HMMs (e.g. Pradel 2005; McClintock *et al*. 2020). Full technical details can be found in Burnham & White (2002), but I describe their approach in the context of individual random effects for the state transition probabilities (Eq. 4) in Section S1 of the Supplementary Material. Burnham & White (2002) demonstrated that their approximate random effects estimator worked well for the Cormack-Jolly-Seber model with a single temporal random effect on survival probability, but, as an approximate method, it is not immediately clear whether or not its simple extension to *N* (*N* – 1) individual random effects for the state transition probabilities in a *N*-state HMM would also perform well. One known issue with this approach is that it becomes unreliable when any *γ_m,i,j_* for the fixed effects model (Eq. 2) is estimated near 0 or 1, and one could suspect that such boundary issues will become more likely as *M* increases, *T_m_* decreases, and *N* increases. Bayesian analogues to the two-stage approach of Burnham & White (2002) could be less susceptible to these boundary issues (Hooten *et al*. 2016, 2019).

More recently, the R package Template Model Builder (TMB; Kristensen *et al*. 2016), which relies on reverse-mode automatic differentiation and the Laplace approximation for high dimension integrals, has made ML inference for continuous random effects much more tractable. `TMB` is less “plug-and-play” than the approach of Burnham & White (2002) because it currently requires advanced programming skills to custom code the HMM likelihood (Eq. 4) based on a `C++` template. In addition, little is currently known about how well the Laplace approximation performs for mixed HMMs or for sample sizes typical of animal telemetry studies (but for relevant applications see Albertsen *et al*. 2015; Auger-Méthé *et al*. 2017; Whoriskey *et al*. 2017; Benhaiem *et al*. 2018). Nevertheless, continuous random effect HMMs implemented using `TMB` can be fitted exceptionally fast, are amenable to more complex correlation structures, and likely do not suffer from the same boundary issues as the approximate approach of Burnham & White (2002). These promising capabilities of `TMB` facilitate large-scale simulation experiments for comparing the performance of these different approaches for modelling individual variation in the hidden state process.

## 3 Simulation study

### 3.1 Simulation methods

Based on 2-state HMMs commonly used in analyses of telemetry data (e.g. Franke *et al*. 2004; Morales *et al*. 2004), I performed a simulation experiment to evaluate the performance of models that include no effects, fixed effects, discrete random effects, and continuous random effects to account for individual heterogeneity in state transition probabilities. Two sets of simulations were performed (see Table 1). In the first (hereafter “without covariates”), the simulated data included no measurable individual-level covariates and the fitted models included no explanatory covariate terms for the state transition probabilities. In the second (hereafter “with covariates”), the simulated data included a measurable individual-level covariate and the models were fitted both with and without terms for the covariate effect on the state transition probabilities. R (R Core Team 2020) code for simulating data and fitting all models can be found in the Supplementary Material.

**Table 1.**
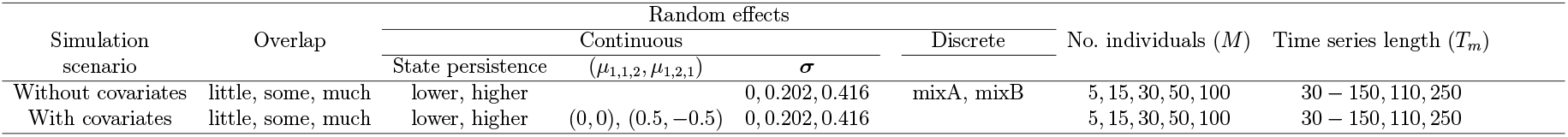
Design points for simulation scenarios with and without measurable covariate effects based on the degree of overlap in the state-dependent distributions, the support of the random effects (continuous or discrete), the number of individuals, and the length of the individual time series. Scenarios without covariates and continuous random effects included logit^−1^(μ_1,2_) = logit^−1^(μ_2,1_) ∈ {0.5, 0.25} (“lower” and “higher” state persistence, respectively). Scenarios with covariates and continuous random effects included logit^−1^(μ_0,1,2_) = logit^−1^(μ_0,2,1_) ∈ {0.5, 0.25} and (μ_1,1,2_, μ_1,2,1_) ∈ {(0, 0), (0.5, −0.5)}. There were therefore 3 × (2 × 3 + 2) × 5 × 3 = 360 scenarios without covariates and 3 × (2 × 2 × 3) × 5 × 3 = 540 scenarios with covariates.

#### 3.1.1 Without covariates

Data were simulated under five levels of individual heterogeneity: 1) no individual heterogeneity (*σ*_1,2_ = *σ*_2,1_ = 0); 2) continuous random effects with “moderate” heterogeneity (*σ*_1,2_ = *σ*_2,1_ = 0.202); 3) continuous random effects with “high” heterogeneity (*σ*_1,2_ = *σ*_2,1_ = 0.416); 4) *K* = 2 discrete random effects with *π*^(1)^ = 0.6, 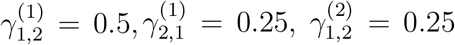, and 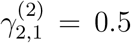 (hereafter “mixA”); and 5) *K* = 2 discrete random effects with *π*^(1)^ = 0.6, 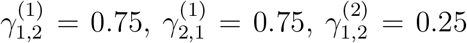, and 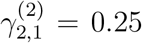 (hereafter “mixB”). For the 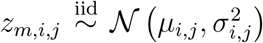 continuous random effect scenarios with *σ*_1,2_ = *σ*_2,1_ ∈ {0, 0.202, 0.416}, I included two levels for state persistence: 1) logit^−1^(*μ*_1,2_) = logit^−1^(*μ*_2,1_) = 0.5 (hereafter “lower” state persistence), corresponding to *γ*_*m*,1,1_ = *γ*_*m*,2,2_ = 0.5 when *σ*_1,2_ = *σ*_2,1_ = 0; and 2) logit^−1^(*μ*_1,2_) = logit^−1^(*μ*_2,1_) = 0.25 (hereafter “higher” state persistence), corresponding to *γ*_*m*,1,1_ = *γ*_*m*,2,2_ = 0.75 when *σ*_1,2_ = *σ*_2,1_ = 0. Scenarios with lower state persistence and *σ*_1,2_ = *σ*_2,1_ = 0.202 correspond to a mean of 0.5 and standard deviation of 0.05 on the state transition probability scale, whereas those with lower state persistence and *σ*_1,2_ = *σ*_2,1_ = 0.416 correspond to a mean of 0.5 and standard deviation of 0.10. For the discrete random effect scenarios, “mixA” can be considered less heterogeneous than “mixB” (because the “mixA” mixtures are more similar). All parameter values were chosen to keep state transition probabilities roughly between 0.1 and 0.9 (see Fig. S1 in Supplementary Material), thereby helping to reduce potential parameter boundary issues during model fitting and prevent either state from being relatively rare (Beyer *et al*. 2013). For simplicity, data were generated with ***δ***_*m*_ = (0.5, 0.5) for *m* = 1,…, *M* in all scenarios.

I limited simulated observations to a single data stream generated from a (state-dependent) gamma distribution for “step length”, 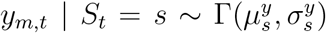 for *S_t_* ∈ {1, 2}, with varying degrees of overlap between the states based on the mean 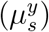 and standard deviation 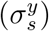. These correspond to “little” overlap 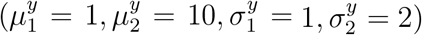, “some” overlap 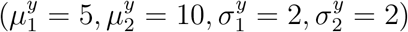, and “much” overlap 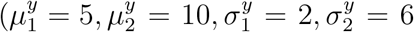; Fig. 1). The Kolmogorov-Smirnov test statistics for these distributions are respectively 0.99, 0.80, and 0.49, where 0 indicates the distributions are identical and 1 indicates no overlap. Standard HMMs are known to perform poorly when the state-dependent distributions overlap (e.g. Beyer *et al*. 2013; Jonsen 2016), but I included these scenarios to assess whether or not the inclusion of individual effects somehow alters this behaviour. While movement HMMs typically include two data streams (step length and turn angle), the number of data streams is arbitrary for my purposes. I therefore chose a single data stream in order to minimize the number of observations and parameters to be estimated, thereby reducing run times and facilitating interpretation across a large number of simulated scenarios.

**Figure 1.**
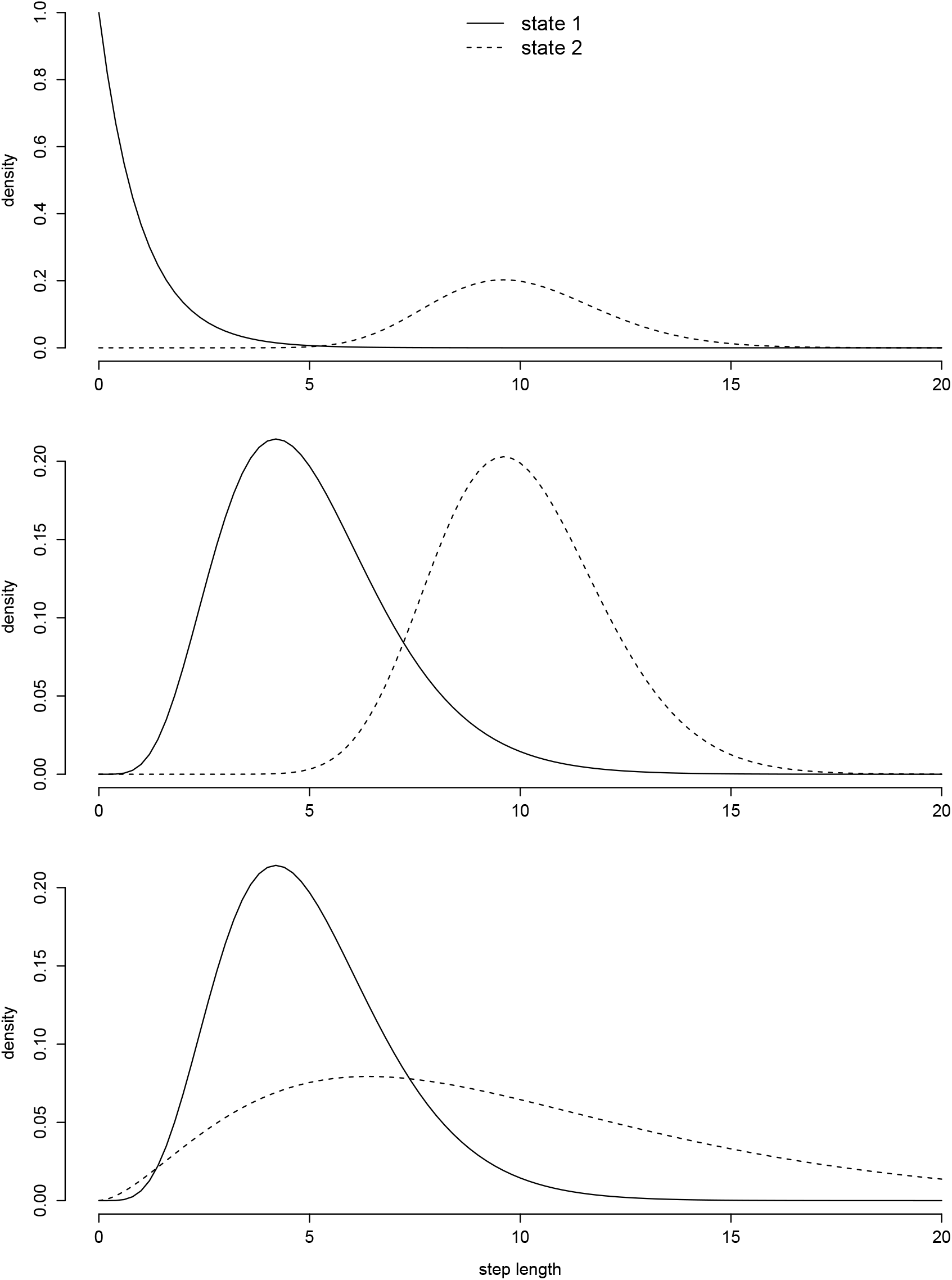
State-dependent observation distributions for “step length” in simulated scenarios with “little” (top panel), “some” (middle), and “much” (bottom) overlap.

Simulated sample sizes were chosen based on animal-borne telemetry studies that typically tend to include relatively few individuals, but relatively long time series for each individual (e.g. Morales *et al*. 2004; Langrock *et al*. 2012; McClintock *et al*. 2013;Towner *et al*. 2016; DeRuiter *et al*. 2017; Isojunno *et al*. 2017). I included five levels for the number of individuals *M* ∈ {5, 15, 30, 50, 100} and three levels for the number of observations per individual (30 – 250, 110, 250). For scenarios with 30 – 250 observations per individual, individuals were assigned to one of *T_m_* ∈ {30, 50, 70, 150, 250} in equal proportions such that 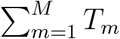 is identical for scenarios with 30 – 250 or 110 observations per individual. I limited *T_m_* ≤ 250 both to reduce computation time and to reflect the lengths of time series in many prominent applications of animal movement HMMs (e.g. Morales *et al*. 2004; Jonsen *et al*. 2005) and mixed HMMs (e.g. Towner *et al*. 2016; DeRuiter *et al*. 2017; Isojunno *et al*. 2017).

For each of the 360 scenarios examined, up to nine models were fitted to 400 simulated data sets using maximum likelihood methods. These models included 2-state HMMs with no individual effects (“null”; Eq. 1), individual fixed effects (“fixed”; Eq. 2), discrete random effects with *K* ∈ {2, 3, 4, 5, 6} mixtures (“mix2”, “mix3”, “mix4”, “mix5”, and “mix6”, respectively; Eq. 3), approximate continuous random effects estimated from the “fixed” model based on Burnham & White (2002; “BW”), and continuous random effects using numerical integration (“TMB”; Eq. 4) for the state transition probabilities. To reduce simulation run times, discrete random effect models with *K* = 5 or *K* = 6 were only fitted if the *K* – 1 mixture model resulted in a lower bias-corrected Akaike’s Information Criterion (AIC_*c*_; Burnham & Anderson 2002) value relative to the *K* – 2 mixture model. Models with *K* = 5 or *K* = 6 are therefore only included in model selection and multimodel inference results (see Section 3.1.4). Simulated data were generated using the `simData` function in R package `momentuHMM` (version 1.5.2; McClintock & Michelot 2018). The R package `TMB` (Kristensen *et al*. 2016) was used for fitting model TMB, and the `momentuHMM` function `fitHMM` was used for fitting all other models. The `momentuHMM` function `randomEffects` was used for implementing the BW approach based on the maximum likelihood estimates of the fixed effects model returned by `fitHMM`.

#### 3.1.2 With covariates

To investigate estimator performance in the presence of a (measurable) continuous-valued individual covariate, I performed an additional set of simulations with logit^−1^(*μ*_0,1,2_) = logit^−1^(*μ*_0,2,1_) ∈ {0.5, 0.25}, *σ*_1,2_ = *σ*_2,1_ ∈ {0, 0.202, 0.416}, *M* ∈ {5, 15, 30, 50, 100}, and the same three levels for the degree of overlap (“little”, “some”, “much”) and the number of observations per individual (30 – 250, 110, 250). Two covariate scenarios were included: 1) *μ*_1,1,2_ = *μ*_1,2,1_ = 0; and 2) *μ*_1,1,2_ = 0.5 and *μ*_1,2,1_ = −0.5, where 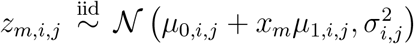 and *x_m_* is a measurable individual covariate drawn from a standard normal distribution. The covariate scenarios with *μ*_1,1,2_ = *μ*_1,2,1_ = 0 were included to investigate the potential for inferring spurious covariate effects from the different models, whereas the scenarios with *μ*_1,1,2_ = 0.5 and *μ*_1,2,1_ = −0.5 have *γ*_*m*,1,2_ and *γ*_*m*,2,1_ increasing and decreasing with *x_m_*, respectively (Fig. S2). For these 540 scenarios each consisting of 400 simulated data sets, I fitted the fixed effect model (with no covariate effects) and the null, finite mixture, and continuous random effect models both with and without terms for the covariate effects (up to 17 models total). For the finite mixture models, each of the *K* mixtures included context-specific 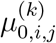 and 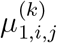 parameters (e.g. DeRuiter *et al*. 2017), from which population-level covariate effects were derived for comparisons with other models (see Section S2.1.3 in Supplementary Material).

#### 3.1.3 Estimator performance

For both sets of simulations, estimator performance for ***σ*** = (*σ*_1,2_, *σ*_2,1_), **Γ**_*m*_ = (*γ_m,i,j_*) for *m* = 1,…, *M*, 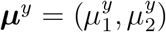, and 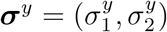 was evaluated based on mean bias, 95% confidence interval coverage, and standard error (or confidence interval length). For the set of simulations with covariates, summaries of estimator bias and precision for *μ*_1,1,2_ and *μ*_1,2,1_ were based on medians because with smaller sample sizes the means for these (unconstrained) parameters could be heavily influenced by a small number of outliers when estimated state transition probabilities fell on the boundary of the parameter space. I examined the performance of the Viterbi algorithm for global state decoding (e.g. Zucchini *et al*. 2016) while accounting for state classification agreement entirely due to chance using the Kappa statistic (Congalton 1991; Beyer *et al*. 2013), which ranges from 0 (entirely chance agreement) to 1 (perfect agreement not attributable to chance). I also evaluated the proportion of estimated local state probabilities (based on the forward-backward algorithm; e.g. Zucchini *et al*. 2016) with at least 0.50 and 0.20 probability assigned to the true state: 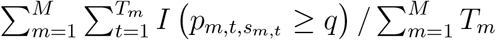, where *I*() is the indicator function, *p_m,t,j_* is the estimated probability of state *j* for individual *m* at time *t, s_m,t_* is the true state for individual *m* at time *t*, and *q* ∈ {0.50, 0.20}. To facilitate comparisons across models, individual-level estimates for state transition probabilities were derived from the maximum likelihood estimates for the finite mixture models. For global state decoding and local state probabilities, I used modified Viterbi and forward-backward algorithms accommodating finite mixtures (see Sections S2.1.3-S2.1.5 in Supplementary Material). For the continuous random effect models, the standard Viterbi and forward-backward algorithms were used based on the shrinkage estimates for each individual 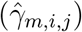.

#### 3.1.4 Model selection and multi-model inference

Standard model selection criteria are often used to choose among competing HMMs (e.g. DeRuiter *et al*. 2017; Isojunno *et al*. 2017; Pohle *et al*. 2017). For both sets of simulations, I evaluated the performance of AIC_*c*_ in selecting among competing models for individual heterogeneity in state transition probabilities by calculating the standard AIC_*c*_ for the null, fixed, and finite mixture models, the conditional AIC_*c*_ for the BW random effects model (Burnham & White 2002), and the marginal AIC_*c*_ for the TMB random effects model using 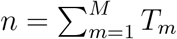 as the sample size. While the BW conditional AIC_*c*_ is comparable with the AIC_*c*_ for the null, fixed, and finite mixture models, it’s unclear how well the marginal AIC_*c*_ for TMB will perform for selecting among fixed and random effect models (e.g. Bolker *et al*. 2009; Gimenez & Choquet2010). I therefore also performed a likelihood ratio test (LRT) comparing the null and TMB models as suggested by Gimenez & Choquet (2010). To my knowledge, there is currently no straightforward way to calculate a conditional AIC even for simpler (non-Markov) random effect models fitted in `TMB`; for example, the `TMB`-based generalized linear mixed modelling R package `glmmTMB` (Brooks *et al*. 2017) only provides marginal AIC. Multi-model inference can be used to account for model selection uncertainty, and I calculated model-averaged estimates, standard errors, and 95% confidence intervals for **Γ**_*m*_, ***μ***^*y*^, and ***σ***^*y*^ based on AIC_*c*_ weights (Burnham & Anderson 2002) for four sets of candidate models: 1) null and finite mixture models (hereafter “modMix”); 2) null, fixed, and finite mixture models (“modFix”); 3) null, fixed, finite mixture, and BW models (“modBW”); and 4) null, finite mixture, and TMB models (“modTMB”). For simulations with covariates, model-averaged estimates for *μ*_1,*i,j*_ were calculated from AIC_*c*_ weights for the models that included the covariate effect.

#### 3.1.5 Nested loop plots

Simulation scenario results are presented using the nested loop plot of Rücker & Schwarzer (2014). Similar to a time series plot, nested loop plots serve to present a large number of simulation results by putting all scenarios into a lexicographical order and arranging them consecutively along the horizontal axis. The quantity of interest (e.g. bias, coverage) is then plotted on the vertical axis. The decision on how to nest the results is subjective, with the top-level of nesting receiving the most emphasis. For BW and modBW, results are only reported for those scenarios where at least 50 of the simulated data sets yielded admissible **Γ**_*m*_ estimates from the fixed effects model, where I considered any 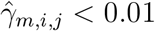 or 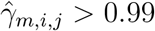 as inadmissible.

### 3.2 Simulation results

#### 3.2.1 Summary

Given the large amount of information afforded by so many simulation scenarios, I first provide a brief summary of the main findings before delving into more detail on the simulation results without (Section 3.2.2) and with (Section 3.2.3) covariates. As expected, across all simulated scenarios the data-generating model tended to perform better in terms of state assignment and parameter estimation as sample sizes (*M* and *T_m_*) increased, the degree of overlap in state-dependent distributions decreased, and state persistence increased. There was generally little difference in state assignment among the null, fixed, and random effect models, indicating mixed HMMs did not improve classification of the hidden state process. Owing to negative bias and poor precision, the continuous random effect variance parameters (***σ***) generally proved difficult to reliably estimate except with larger sample sizes (e.g. *M* > 50). While little difference was found between the various models in terms of state-dependent distribution parameter (***μ***^*y*^, ***σ***^*y*^) estimation, discrete random effect models exhibited poor coverage of state transition probabilities (*γ_m,i,j_*) under continuous variation, and continuous random effect models exhibited inflated standard errors for *γ_m,i,j_* under discrete variation. Model selection and model-averaging based on AIC_*c*_ generally worked well when the data-generating model was included in the candidate model set, but it did little to mitigate model misspecification in the “modMix” and “modFix” model sets for the continuous variation scenarios. Finally, in the “with covariates” scenarios, discrete random effect models again proved to be a poor (and potentially misleading) approximation for continuous variation, often resulting in a substantial reduction in confidence interval coverage for the covariate effects (*μ*_1,*i,j*_) and, in some cases, less accurate state assignment relative to the other models.

#### 3.2.2 Without covariates

##### State assignment

The degree of overlap between the (state-dependent) observation distributions was by far the most important factor for state estimation. The performance of each model declined as overlap increased, but, within each level of overlap (“little”, “some”, “much”), there was little difference among the null, fixed, and random effect models (Fig. 2 and Table S1 in Supplementary Material). When there was “some” or “much” overlap, performance tended to slightly improve as state persistence increased (Fig. S3) and as individual heterogeneity, the number of individuals, and the lengths of the time series increased. After accounting for chance agreement, the Viterbi algorithm for global state decoding produced an average correct state assignment of 99% (range: 99–99%) for “little” overlap, 82% (range: 76–84%) for “some” overlap, and 51% (range: 40–57%) for “much” overlap across all scenarios and models. The proportion of correct state assignments using the Viterbi algorithm therefore closely corresponded to the degree of overlap in the state-dependent distributions as calculated by the Kolmogorov-Smirnov test statistic (respectively 0.99, 0.80, and 0.49). Estimated state probabilities tended to assign at least 0.5 probability to the true state > 75% of the time (and at least 0.20 probability to the true state > 90% of the time) across all scenarios and models, indicating local state probabilities are better able to account for uncertainty in state assignment that is attributable to overlap in state-dependent observation distributions. While the discrete random effect models tended to assign less probability to the true state with “some” or “much” overlap, the inclusion of individual fixed, discrete random, or continuous random effects generally made little difference in local state probabilities relative to the null model.

**Figure 2.**
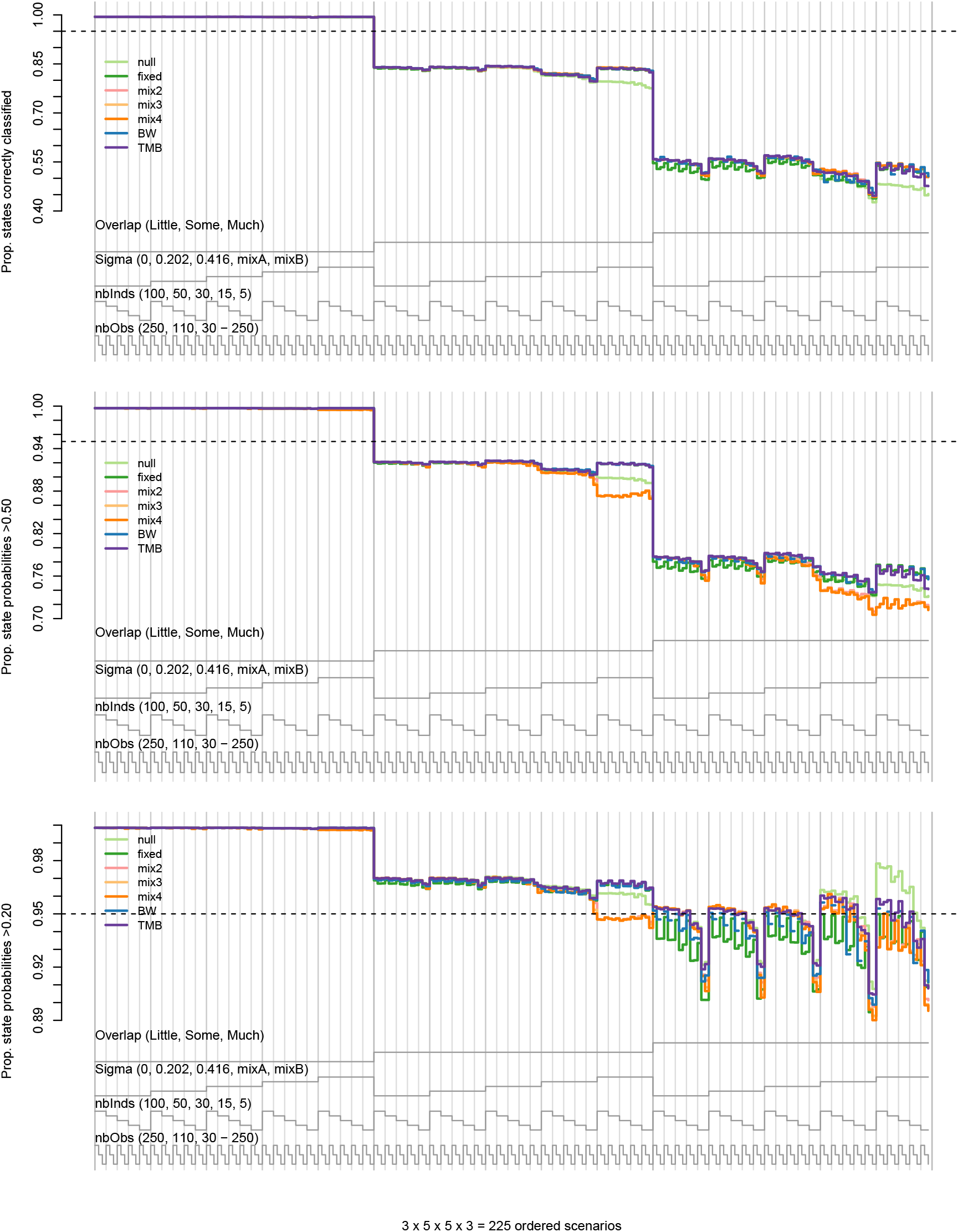
Nested loop plots for the proportion of Viterbi-decoded states that were correctly classified after accounting for chance agreement (top) and the proportion of estimated state probabilities in which the true state received at least 0.50 (middle) or 0.20 (bottom) probability from 225 simulated scenarios without covariate effects. Scenarios are ordered from outer to inner loops by the degree of state-dependent distribution overlap (“Overlap”), individual heterogeneity (“Sigma”), number of individuals (“nbInds”), and length of time series (“nbObs”). Comparisons are for the null (light green), fixed (dark green), mix2 (pink), mix3 (light orange), mix4 (dark orange), BW (blue), and TMB (purple) models. Continuous random effect scenarios are limited to those with “higher” state persistence.

##### Random effect variance

Models that included continuous random effects (BW and TMB) generally performed best in terms of bias, coverage, and confidence interval length with “little” overlap, “higher” state persistence (logit^−1^(*μ*_1,2_) = logit^−1^(*μ*_2,1_) = 0.25), “high” heterogeneity (*σ*_1,2_ = *σ*_2,1_ = 0.416), and larger sample sizes (Table 2, Fig. 3). With percent confidence interval lengths (calculated as 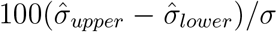 for *σ* > 0) for individual scenarios ranging from 81% (*M* = 100, *T_m_* = 250, and “little” overlap) to 5139% (*M* = 5, *T_m_* = 30 – 250, and “much” overlap), both estimators tended to produce very wide confidence intervals for ***σ*** as the number of individuals decreased, the length of the time series decreased, and the degree of overlap increased. With “little” overlap and few individuals, BW tended to perform better than TMB, and, because its point estimates and confidence intervals can include zero, BW also tended to perform better than TMB for scenarios with no heterogeneity (i.e. *σ*_1,2_ = *σ*_2,1_ = 0; Fig. 3). However, with individual heterogeneity and “some” or “much” overlap, BW tended to underperform relative to TMB as the number of individuals increased and the length of the time series decreased, particularly when there was “lower” state persistence. Under these conditions, the fixed model increasingly tended to estimate at least one state transition probability near a boundary, thereby making it unsuitable for the BW approach. This likely explains the increased negative bias for BW with “some” and “much” overlap, as the subset of simulated data sets that produced admissible **Γ**_*m*_ estimates from the fixed model will exhibit truncated tails for the random effect distributions (and hence smaller ***σ***). However, the fact that the bias tended to be greater for *σ*_2,1_ than for *σ*_1,2_ also suggests that the sampling variance-covariance matrix approximation of Burnham & White (2002) might be inadequate in accounting for the additional state uncertainty under scenarios with non-negligible overlap. TMB therefore appears to be more robust to overlap in state-dependent observation distributions and shorter time series with “moderate” to “high” heterogeneity, but, owing to negative bias and poor precision, neither TMB nor BW performed particularly well except in scenarios with “little’ overlap, “high” heterogeneity, and at least *M* = 50 individuals.

**Figure 3.**
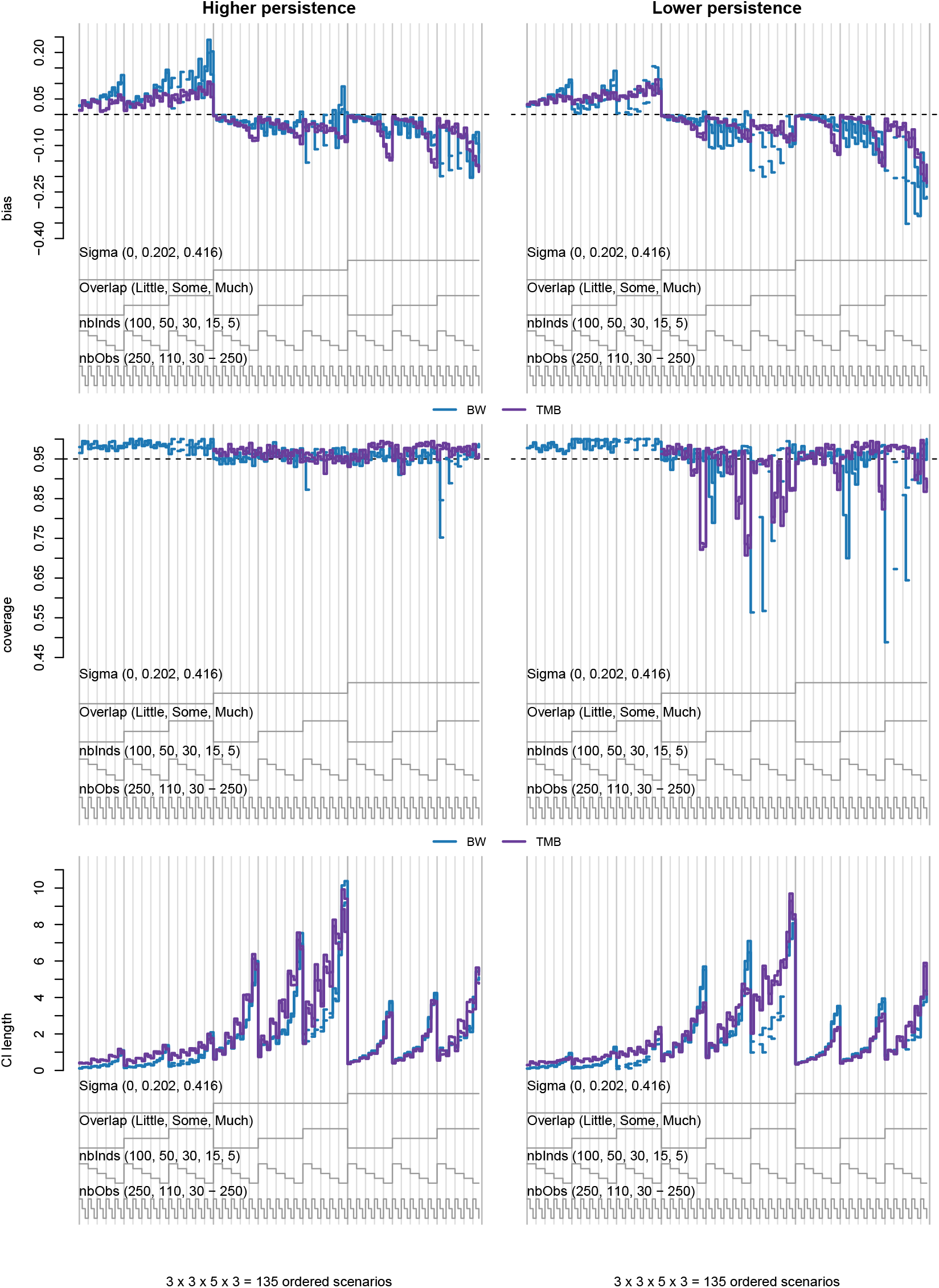
Nested loop plots for BW (blue) and TMB (purple) mean bias (top), 95% confidence interval coverage (middle), and confidence interval length (bottom) for σ_1,2_ and σ_2,1_ from 135 simulated scenarios without covariates that included “higher” (left column) or “lower” (right column) state persistence. Missing values for BW indicate scenarios with < 50 data sets producing admissible estimates for the state transition probabilities from the fixed effects model. Coverage for TMB was 0% for all scenarios with σ_1,2_ = σ_2,1_ = 0.

**Table 2.** BW and TMB overall mean percent relative bias, 95% confidence interval coverage, and percent confidence interval length for *σ*_1,2_ and *σ*_2,1_ by design points for state-dependent distribution overlap, state persistence, individual heterogeneity (***σ***), number of individuals, and time series length in simulations without covariates.

| Model | Parm. | Overlap | | | State persistence | | $\sigma$ | | No. individuals ($M$) | | | | | Time series length ($T_m$) | | | Overall |
| --- | --- | --- | --- | --- | --- | --- | --- | --- | --- | --- | --- | --- | --- | --- | --- | --- | --- |
|  |  | Little | Some | Much | Higher | Lower | 0.202 | 0.416 | 100 | 50 | 30 | 15 | 5 | 250 | 110 | 30 - 250 |  |
| Percent relative bias |  |  |  |  |  |  |  |  |  |  |  |  |  |  |  |  |  |
| BW | $\sigma_{1,2}$ | -6.6 | -13.4 | -46.3 | -16.8 | -22.9 | -21.9 | -17.6 | -16.4 | -18.9 | -21.1 | -22.4 | -19.6 | -15.9 | -23.7 | -19.9 | -19.8 |
| BW | $\sigma_{2,1}$ | -6.6 | -20.4 | -22.6 | -11.6 | -20.5 | -17.1 | -14.8 | -13.4 | -15.0 | -16.3 | -18.5 | -16.1 | -9.8 | -18.0 | -21.3 | -15.9 |
| TMB | $\sigma_{1,2}$ | -12.4 | -14.6 | -22.8 | -16.9 | -16.3 | -19.6 | -13.6 | -6.7 | -9.6 | -12.8 | -18.5 | -35.4 | -12.8 | -17.7 | -19.3 | -16.6 |
| TMB | $\sigma_{2,1}$ | -12.1 | -15.3 | -21.5 | -16.8 | -15.8 | -19.5 | -13.2 | -6.1 | -8.9 | -12.3 | -18.6 | -35.7 | -12.9 | -17.1 | -19.0 | -16.3 |
| Coverage |  |  |  |  |  |  |  |  |  |  |  |  |  |  |  |  |  |
| BW | $\sigma_{1,2}$ | 0.96 | 0.96 | 0.89 | 0.95 | 0.93 | 0.94 | 0.94 | 0.91 | 0.93 | 0.94 | 0.95 | 0.97 | 0.93 | 0.94 | 0.96 | 0.94 |
| BW | $\sigma_{2,1}$ | 0.96 | 0.94 | 0.96 | 0.95 | 0.95 | 0.95 | 0.95 | 0.93 | 0.95 | 0.95 | 0.96 | 0.96 | 0.95 | 0.95 | 0.95 | 0.95 |
| TMB | $\sigma_{1,2}$ | 0.96 | 0.95 | 0.95 | 0.97 | 0.94 | 0.94 | 0.96 | 0.96 | 0.96 | 0.96 | 0.95 | 0.92 | 0.97 | 0.94 | 0.95 | 0.95 |
| TMB | $\sigma_{2,1}$ | 0.96 | 0.95 | 0.94 | 0.97 | 0.94 | 0.94 | 0.96 | 0.96 | 0.96 | 0.96 | 0.95 | 0.92 | 0.97 | 0.94 | 0.95 | 0.95 |
| Percent confidence interval length |  |  |  |  |  |  |  |  |  |  |  |  |  |  |  |  |  |
| BW | $\sigma_{1,2}$ | 408 | 495 | 710 | 545 | 495 | 1349 | 386 | 203 | 276 | 347 | 507 | 1164 | 409 | 544 | 635 | 521 |
| BW | $\sigma_{2,1}$ | 408 | 477 | 815 | 564 | 519 | 1409 | 400 | 213 | 294 | 368 | 530 | 1197 | 428 | 584 | 638 | 542 |
| TMB | $\sigma_{1,2}$ | 400 | 534 | 897 | 646 | 574 | 1740 | 376 | 277 | 417 | 526 | 733 | 1098 | 447 | 710 | 673 | 610 |
| TMB | $\sigma_{2,1}$ | 402 | 523 | 983 | 679 | 592 | 1799 | 398 | 289 | 438 | 542 | 761 | 1149 | 470 | 730 | 707 | 636 |

##### State transition probabilities

All models generally exhibited little bias in state transition probabilities with “little” or “some” overlap, but performance was more variable in terms of confidence interval coverage and precision (Table 3, Fig. 4). Although the overall mean bias across all **Γ**_*m*_ was close to zero with “much” overlap (Table 3), this is somewhat misleading because the overlap tended to induce increasing negative bias for *γ*_*m*,1,2_ and positive bias for *γ*_*m*,2,1_ as the number of individuals decreased, particularly for the finite mixture scenarios (“mixA” and “mixB”) and the continuous random effect scenarios with lower state persistence (Fig. S4, Tables S2–S3). Despite the null and finite mixture models generally exhibiting little bias under continuous variation, it is worth noting that by construction these models are not able to correctly capture the individual-level state transition probabilities (as indicated by poor coverage and underestimation of uncertainty), and other measures of estimator performance (such as absolute bias or mean squared error) would perhaps better reflect this. With “little” overlap, the null model tended to perform best with no heterogeneity and BW tended to perform best with “moderate” to “high” heterogeneity because TMB tended to underestimate uncertainty (except when *M* = 100). Consistent with its performance for ***σ*** estimation, BW did not perform as well as TMB under “some” or “much” overlap, often exhibiting larger variances and asymmetric biases for *γ*_*m*,1,2_ and *γ*_*m*,2,1_ (albeit with somewhat better coverage).

**Figure 4.**
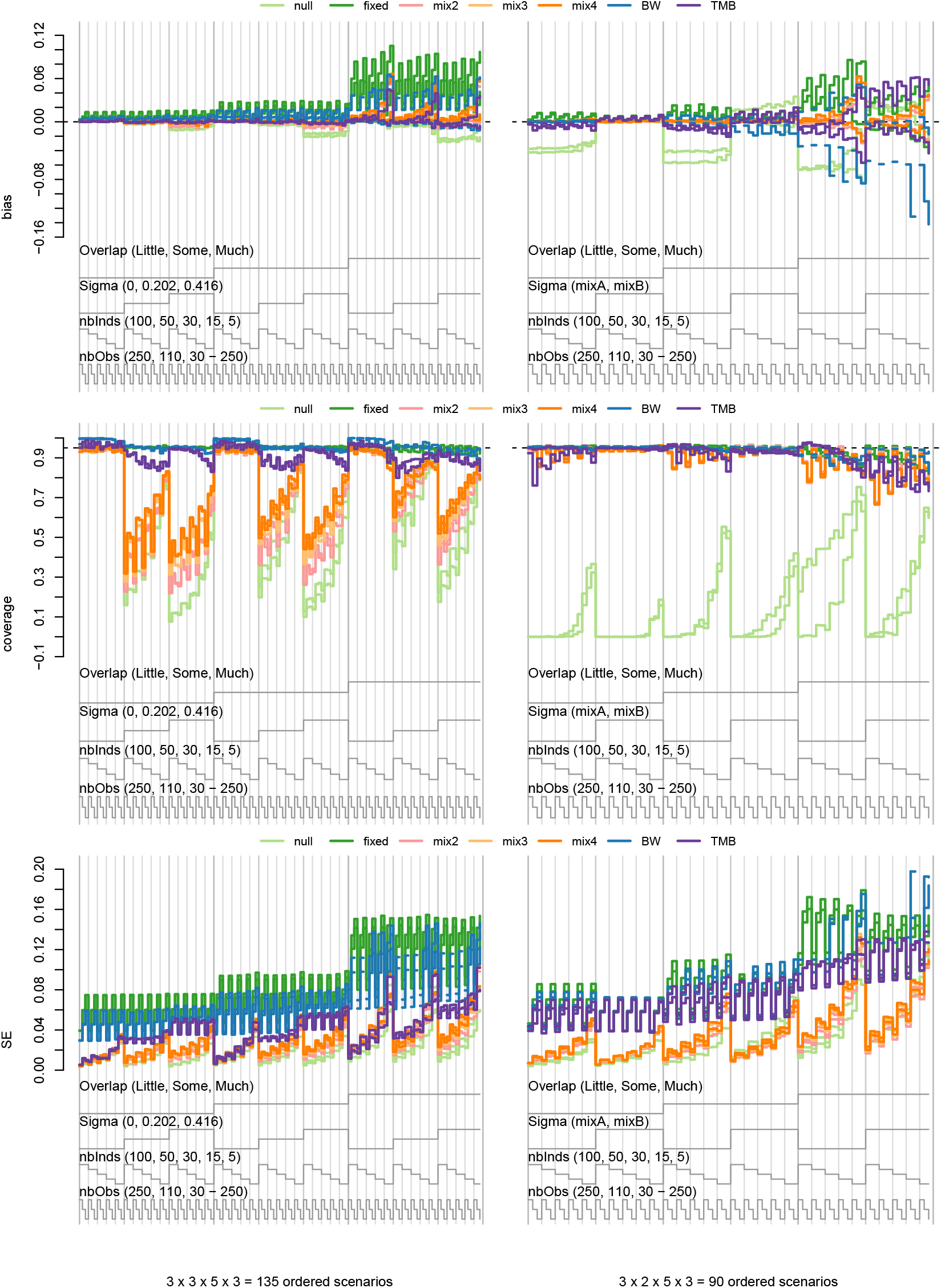
Nested loop plots for mean bias (top row), 95% confidence interval coverage (middle row), and standard error (SE; bottom row) for γ_m,1,2_ and γ_m,2,1_ from simulated scenarios without covariates, including 135 scenarios with “higher” state persistence (left column) and 90 scenarios with “mixA” or “mixB” finite mixtures (right column). Missing values for BW indicate scenarios with < 50 data sets producing admissible estimates.

**Table 3.**
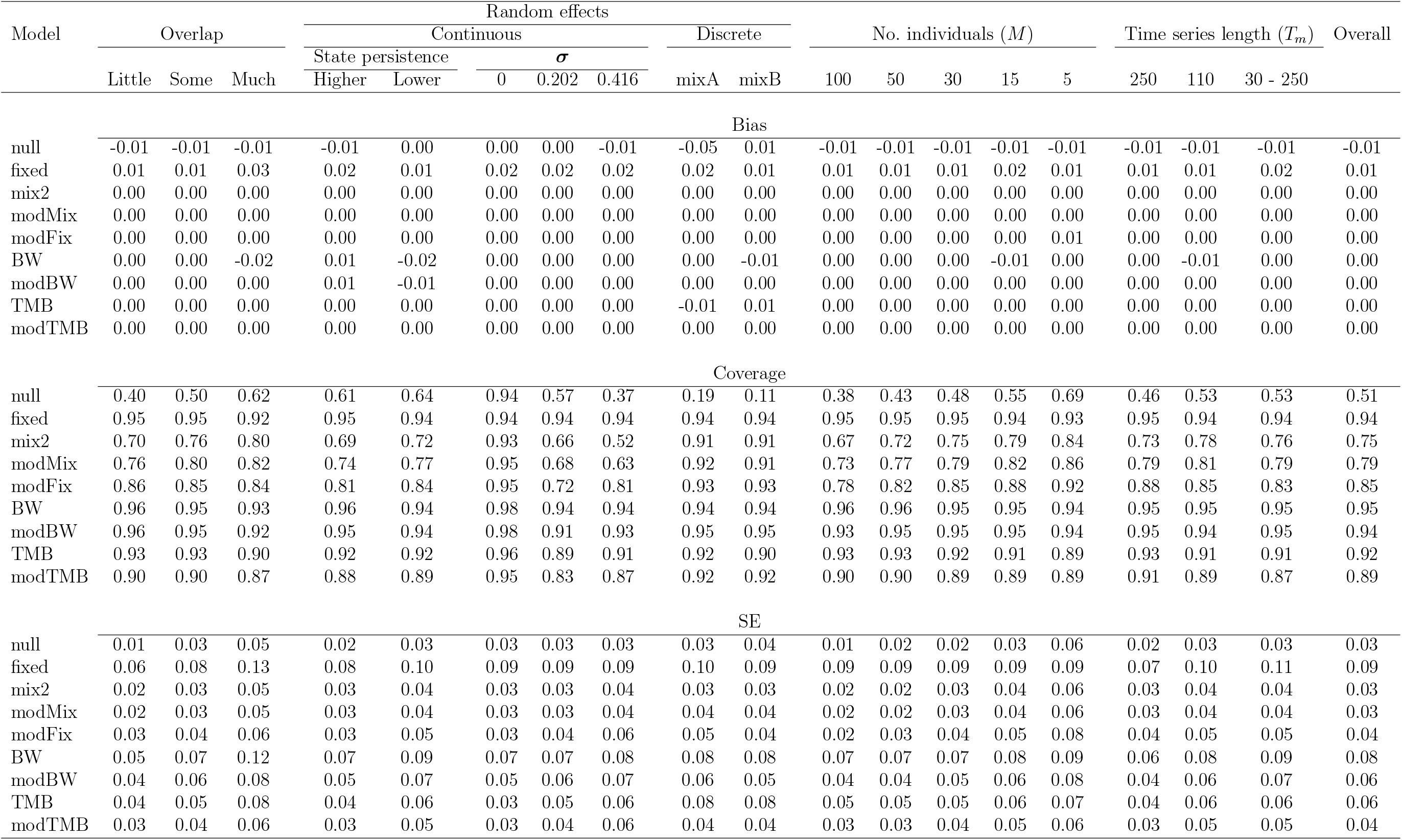
Overall mean bias, 95% confidence interval coverage, and standard error (SE) for both γ_m,1,2_ and γ_m,2,1_ by design points for state-dependent distribution overlap, continuous or discrete random effects, number of individuals, and time series length in simulations without covariates.

As expected, the mix2 model generally performed best in the “mixA” and “mixB” finite mixture scenarios, but coverage was increasingly below the nominal 95% as the number of individuals increased, the length of time series decreased, and the degree of overlap increased. Underestimation of uncertainty by the mix2 model was unexpected under these scenarios, but the finite mixture models did not appear to always be able to adequately distinguish individual-level variability from sampling variability when estimating mixture probabilities for individuals with shorter time series. However, the mix2 model was less prone to reduced coverage for shorter time series under the more distinct “mixB” scenario with “little” or “some” overlap, so this behaviour also appears to depend on the specific characteristics of the mixture distributions. All models except the null generally performed well with data generated under *K* = 2 finite mixtures (albeit with mean standard errors 3.2, 3.5, and 3.7 times larger than mix2 for the TMB, BW, and fixed models, respectively), but the finite mixture models did not generally perform well with data generated under continuous random effects. Under these scenarios, the discrete random effect models tended to perform slightly better in terms of coverage as the number of mixtures increased, but coverage was still well below nominal (as low as 32% with *K* = 4) and tended to decrease as sample sizes increased (Table 3, Fig. 4).

##### State-dependent distributions

There was not much variability in the performance of the models when estimating statedependent probability distribution parameters 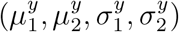. With “little” overlap, all models performed well (see Figs S7–S12 and Tables S4–S7 in Supplementary Material). With smaller sample sizes under “some” or “much” overlap, bias tended to increase and coverage tended to decrease. Under these scenarios, increasing overlap generally resulted in positive bias for 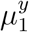, large positive bias for 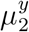, positive bias for 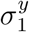, and negative bias for 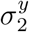, particularly for the continuous random effect scenarios with lower state persistence (Figs S8–S9 and S11–S12). Although coverage remained near nominal under these scenarios, none of the models were able to recover unbiased point estimates for all of the state-dependent distribution parameters. Thus model performance was primarily driven by the degree of overlap, and the inclusion of individual fixed, discrete random, or continuous random effects generally made little difference in state-dependent distribution parameter estimation.

##### Model selection and multi-model inference

When the candidate model set was limited to null and finite mixture models (“modMix”), AIC_*c*_ model selection performed well for data generated under no heterogeneity (*σ*_1,2_ = *σ*_2,1_ = 0) and the finite mixture scenarios, with the generating model (null and mix2, respectively) generally receiving the most AIC_*c*_ support (Figs S13–S14) and the modMix model-averaged parameter estimates performing well (Figs S5–S10, Tables 3 and S2–S7). However, with moderate to high individual heterogeneity (*σ*_1,2_ = *σ*_2,1_ > 0), AIC_*c*_ tended to favor models with increasingly more finite mixtures as individual heterogeneity and sample sizes increased, with poor coverage of model-averaged **Γ**_*m*_ estimates. When the candidate set of models was expanded to include the fixed model (“modFix”), performance was similar for the no heterogeneity and finite mixture scenarios, but the fixed model tended to receive greater support as individual heterogeneity increased, sample sizes increased, and the degree of overlap decreased (Figs S13–S14), resulting in improved (but still less than nominal) performance of the modFix model-averaged **Γ**_*m*_ estimates (Figs S5–S6, Tables 3 and S2–S3).

For the full candidate model set including the null, fixed, finite mixture, and BW models (“modBW”), AIC_*c*_ model selection generally performed well across most simulation scenarios, but support for the generating model tended to decline with smaller sample sizes and higher degrees of overlap (Figs S13–S14). In terms of **Γ**_*m*_ estimation, modBW generally exhibited less bias and greater precision than BW, but performance relative to TMB declined with moderate or high heterogeneity as sample sizes increased, the degree of overlap increased, and state persistence decreased (Figs 4 and S5–S6, Tables 3 and S2–S3). However, under these scenarios modBW tended to perform better than BW, indicating that AIC_*c*_ model averaging can help mitigate poorer performance of BW in scenarios with “some” or “much” overlap that tend to produce inadmissible **Γ**_*m*_ estimates from the fixed model.

For the “modTMB” candidate set, the marginal AIC_*c*_performed better than I expected, but proved to be conservative when selecting among fixed and random effect models for scenarios with moderate or high individual heterogeneity (Figs S15–S16). Under these scenarios, AIC_*c*_ tended to favor the null model as individual heterogeneity decreased, the degree of overlap increased, and sample sizes decreased. Unlike modBW, modTMB generally reduced coverage of **Γ**_*m*_ estimates relative to TMB (Figs 4 and S5–S6, Tables 3 and S2–S3). Likelihood ratio tests between the null and TMB models were less conservative and not as sensitive to the degree of overlap and sample sizes under these scenarios (Figs S15–S16). However, LRTs do not provide a means for selecting between discrete and continuous random effect models, and LRTs tended to strongly favor TMB over the null model when data were generated from finite mixtures (scenarios “mixA” and “mixB”).

#### 3.2.3 With Covariates

For the set of simulations examining measurable individual covariate effects, performance patterns were similar to the set of simulations without covariate effects (see Section 3.2.2) for the null, fixed, finite mixture, and continuous random effect models in terms of state assignment, parameter estimation, and AIC_*c*_ model selection (see Table S8, Figs S17-S25). Both BW and TMB performed well in estimating the covariate effects (Fig. 5), although for (*μ*_1,1,2_, *μ*_1,2,1_) = (0.5, −0.5) with smaller sample sizes and moderate to high heterogeneity, BW exhibited a small negative bias for *μ*_1,1,2_ and a small positive for *μ*_1,2,1_ that was mitigated by AIC_*c*_ model averaging, but became worse as state-dependent distribution overlap increased (Figs. S26 and S27, Tables S9–S10).

**Figure 5.**
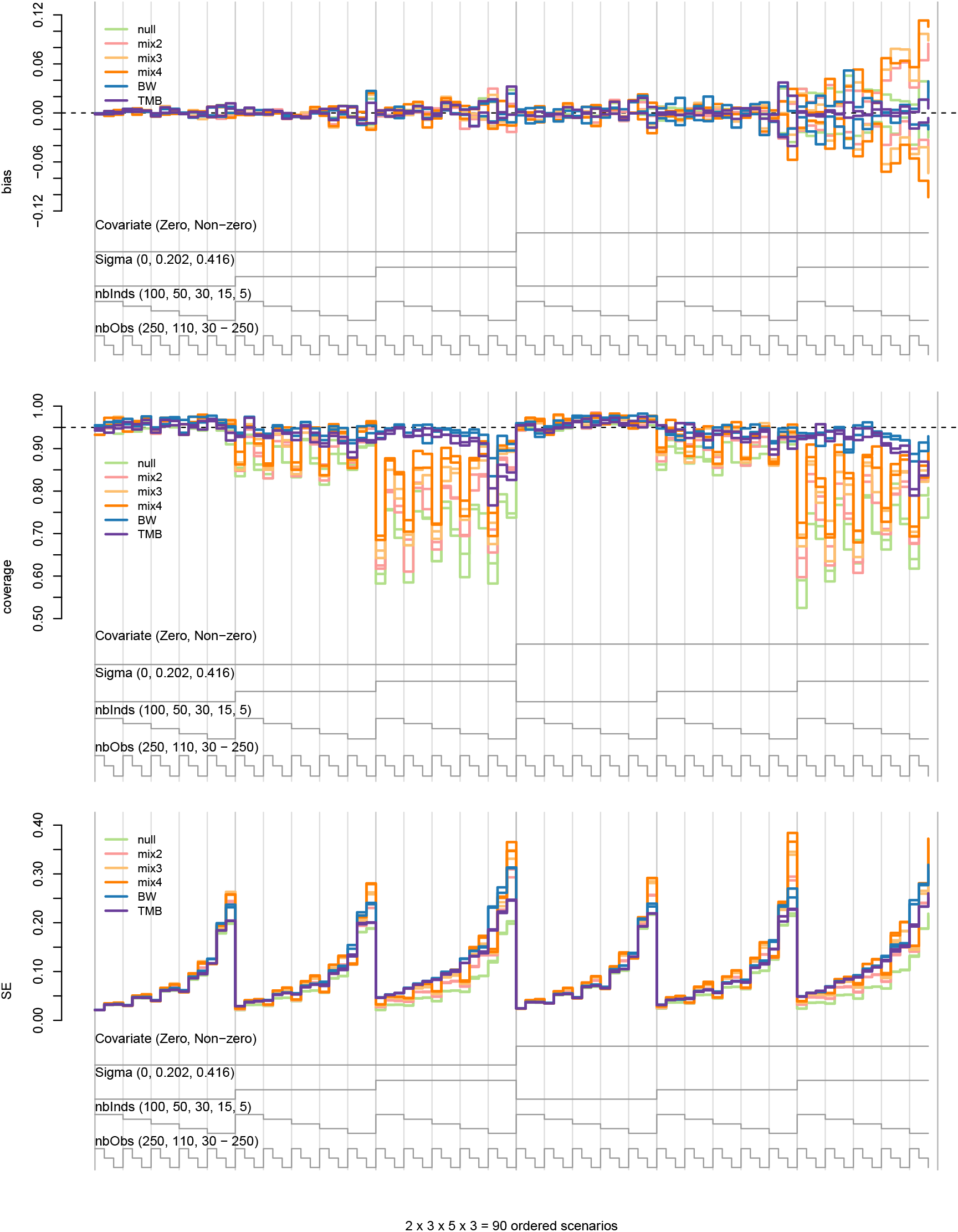
Nested loop plots for median bias (top row), mean 95% confidence interval coverage (middle row), and median standard error (SE; bottom row) for covariate effects μ_1,1,2_ and μ_1,2,1_ from 90 simulated scenarios with “little” overlap, “higher” state persistence, and (μ_1,1,2_, μ_1,2,1_) ∈ {(0, 0), (0.5, −0.5)}. Scenarios are ordered from outer to inner loops by the data-generating values for the covariate effect (“Covariate”, where “Zero” = (0, 0), “Nonzero” = (0.5, −0.5)), individual heterogeneity (“Sigma”), number of individuals (“nbInds”), and length of time series (“nbObs”).

While the finite mixture models performed better than the null model in terms of coverage of *μ*_1,*i,j*_, they generally exhibited greater bias and performance became increasingly poor as individual heterogeneity increased. Coverage of *μ*_1,*i,j*_ for the discrete random effect models increased with the number of mixtures, but it was still well below nominal with *K* = 4 mixtures under higher levels of heterogeneity and “little” overlap (Fig. 5). Coverage for the discrete random effect models actually tended to become worse as sample sizes *increased* and overlap *decreased* (Fig. S26, Tables S9–S10). Under the “best” data-generating scenarios with “little” overlap, *M* = 100, *T_m_* = 250, finite mixture models in some cases reduced coverage of *μ*_1,*i,j*_ by > 37% relative to the TMB model. In a handful of small sample scenarios with *σ*_*i,j*_ > 0 where finite mixture models had near-nominal coverage, this was attributable to large standard errors (up to 101% larger than the TMB model). Because coverage of the finite mixture models was often poor when *μ*_1,*i,j*_ = 0, there was clearly a greater risk of Type I error (i.e. inferring a covariate effect when there is none) when discrete random effects were used to account for continuous individual variation. With AIC_*c*_ for the modMix candidate set tending to support models with both covariate effects and a large number of mixtures (Fig. S25), model averaging did little to mitigate this risk under these scenarios (Fig. S27, Tables S9–S10).

## 4 Example: long-finned pilot whales

To help put my findings in context and illustrate some potential challenges that practitioners may encounter when applying mixed HMMs to animal telemetry data, I revisit a *N* = 4 state multivariate mixed HMM analysis of long-finned pilot whale biotelemetry data originally performed by Isojunno *et al*. (2017). Full details can be found therein, but the data consist of 11 data streams believed to characterize “exploratory” (state 1), “foraging” (state 2), “crowded” (state 3), and “directed” (state 4) diving behaviours for *M* = 15 individuals, with *T_m_* ranging from 50 – 254 (median = 148). In order to limit the number of models in the candidate set, Isojunno *et al*. (2017) first used model selection criteria to determine *N* = 4 was the optimal number of states under the null (*K* = 1) model, then used model selection criteria to choose among finite mixture random effect models (up to *K* = 3), and finally used this model to investigate individual and time-dependent explanatory covariates (e.g. size class, sonar exposure) for the state transition probabilities.

I focus on the second stage of this analysis, where they used AIC and the Bayesian Information Criterion (BIC; e.g. Burnham & Anderson 2002) to select the best-supported random effect model for subsequent covariate modelling and model selection. They found conflicting results based on AIC and BIC, with AIC favoring *K* = 3 mixtures (5.2 unit decrease in AIC relative to the null model) and BIC strongly favoring the null model with *K* = 1. Faced with this apparent conundrum, Isojunno *et al*. (2017) sided with BIC to “avoid selection of overly complex models” given the relatively “weak support for any random effects” afforded by AIC, and proceeded with covariate model fitting and selection under the null model (*K* = 1) with no individual random effects on state transition probabilities.

To explore this further, I re-analyzed the pilot whale data by fitting the null, fixed, finite mixture (up to *K* = 4), and TMB models using the same methods described in Section 3, but with ***δ***_*m*_ assumed to be the stationary distribution (as in Isojunno *et al*. 2017) instead of freely estimated (see Supplementary Material for data and R code). In addition to AIC_*c*_, I calculated standard BIC for the null, fixed, and finite mixture models, as well as the marginal BIC for TMB, using 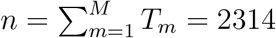 for the sample size (as in Isojunno *et al*. 2017). Unfortunately, the fixed model yielded **Γ**_*m*_ estimates on the boundary, thereby making this data set inadmissible for the BW model. With boundary issues becoming more likely as the number of individuals increases, the lengths of time series decrease, and the number of states increases, this highlights a key limitation of the BW approach in practice.

Despite due diligence by Isojunno *et al*. (2017) in exploring the likelihood surface using 50 sets of randomly drawn starting values for optimization, I found that their finite mixture models failed to converge to global maxima. Relative to those reported by Isojunno *et al*. (2017), my fits increased the log likelihood for the mix2 and mix3 models by 14.6 and 36.7 units, respectively (Table 4). This highlights a common pitfall when attempting to fit complex mixed HMMs to relatively small data sets, where flat likelihood surfaces, local minima, and numerical instability can become increasingly problematic as the number of states and/or parameters increase. Indeed, it’s certainly possible that my fits also failed to converge to the global maxima, although I was unable to improve them any further using hundreds of random normal perturbations of the maximum likelihood estimates as starting values for the optimization.

**Table 4:**
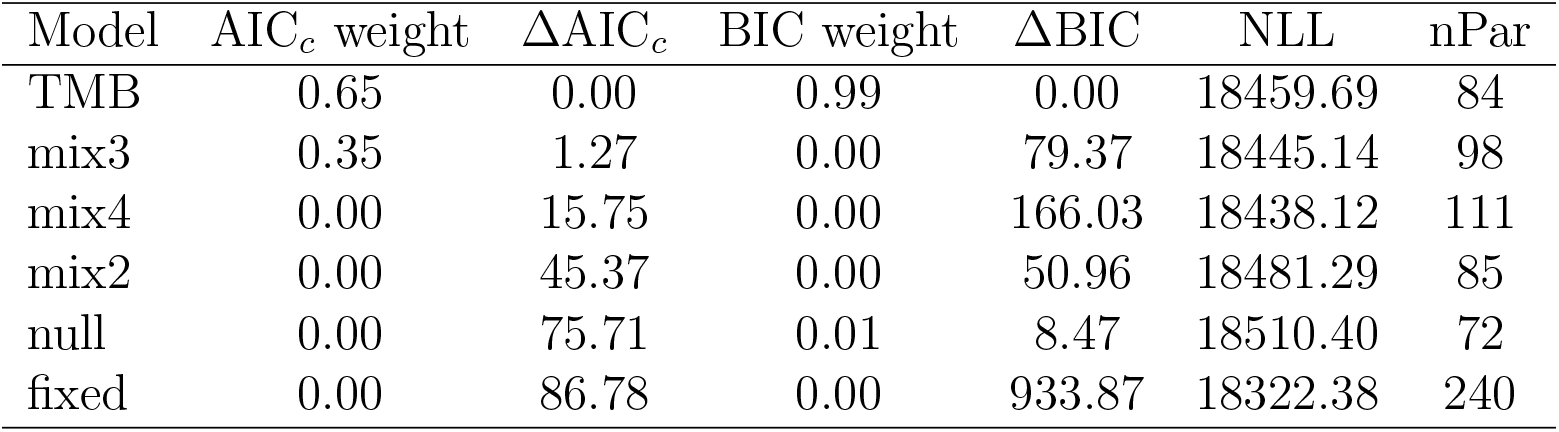
Model selection results for the long-finned pilot whale example. Results include AIC_c_ weights, ΔAIC_c_, BIC weights, ΔBIC, negative log-likelihood value (NLL), and number of parameters (nPar).

Both AIC_*c*_ and BIC now favor TMB (Table 4), and the LRT between the null and TMB model with *N*(*N* – 1) = 12 random effects also favors TMB 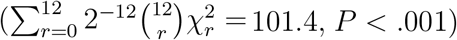. Thus whether one prefers AIC_*c*_, BIC, or LRT, there is clearly evidence of individual variation in the state transition probabilities that is not well explained by null or finite mixture models. As could be expected for relatively short time series with *N* = 4 states, some of the random effect variance estimates were imprecise (Table 5), particularly for state transitions that were relatively rare (Fig. 6). Evidence of individual variation was greatest for transitions from the “exploratory” and “directed” states, and, based on the results of Isojunno *et al*. (2017), this variation was not well explained by any of the individual covariates included in their analysis. However, it is possible that these explanatory covariates could now better account for any additional variation that is not already well explained by the individual random effects.

**Figure 6.**
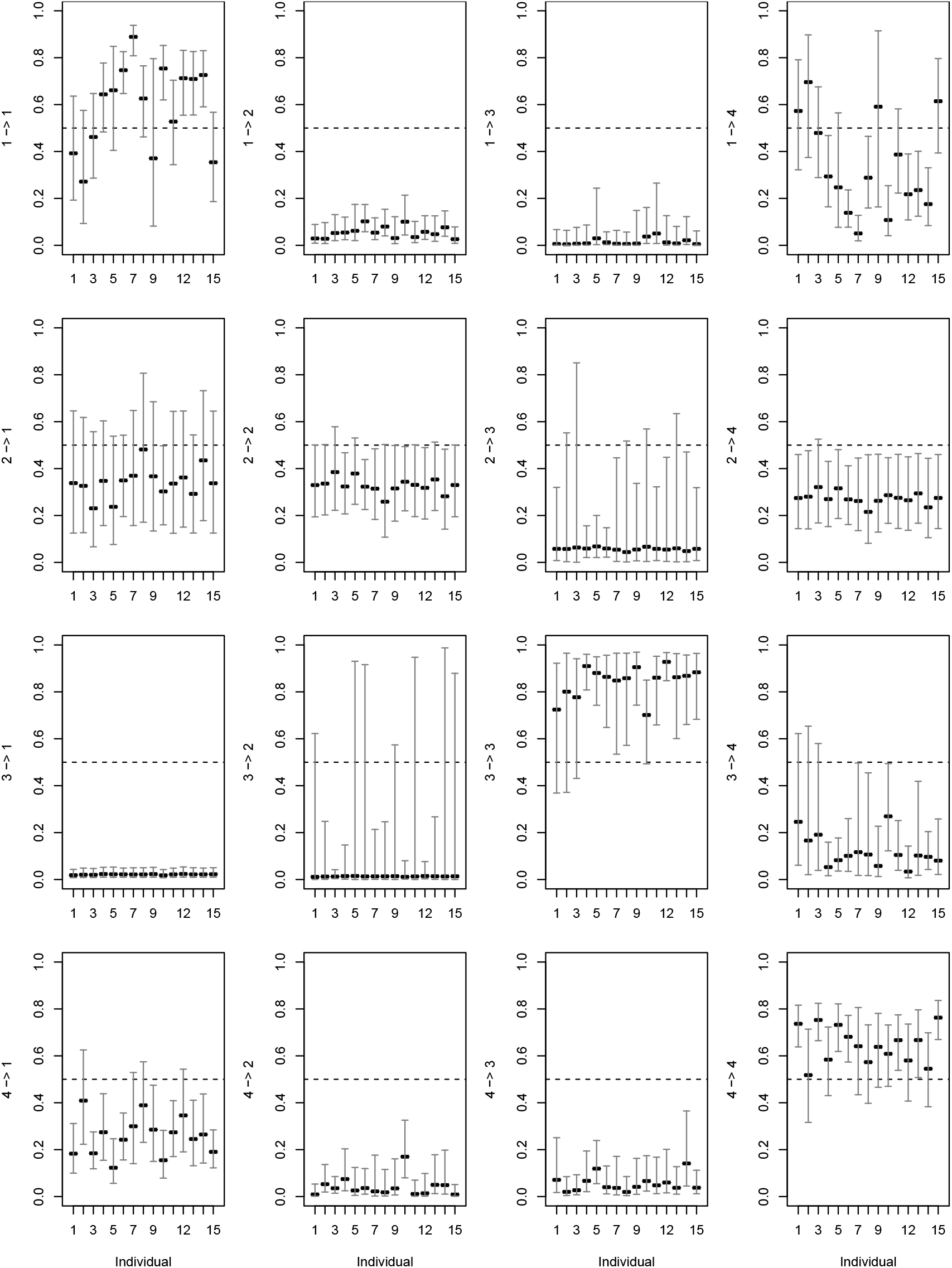
Estimated state transition probabilities (and 95% confidence intervals) among N = 4 states (1 = “exploratory”, 2 = “foraging”, 3 = “crowded”, 4 = “directed”) for M = 15 long-finned pilot whales from the TMB model including continuous individaul-level random effects. Each of the 4 × 4 = 16 state transition probabilities is labeled on the y-axis as “i −> j “, indicating the probability of switching from state i to state j.

**Table 5.** Estimates, standard errors (SE), 95% confidence intervals (Lower, Upper), percent coefficient of variation (%CV), and percent confidence interval length (%CIL) for individual random effect variance parameters (**σ**) from the TMB model fitted to the long-finned pilot whale data.

| Parameter | Estimate | SE | Lower | Upper | %CV | %CIL |
| --- | --- | --- | --- | --- | --- | --- |
| $\sigma_{1,2}$ | 0.47 | 0.27 | 0.17 | 1.35 | 58 | 251 |
| $\sigma_{1,3}$ | 1.17 | 0.69 | 0.40 | 3.39 | 59 | 256 |
| $\sigma_{1,4}$ | 1.16 | 0.30 | 0.71 | 1.92 | 26 | 104 |
| $\sigma_{2,1}$ | 0.56 | 0.41 | 0.15 | 2.03 | 74 | 336 |
| $\sigma_{2,3}$ | 0.25 | 2.96 | 0.00 | 19.63 | 1173 | 7766 |
| $\sigma_{2,4}$ | 0.03 | 0.34 | 0.00 | 2.26 | 1195 | 7895 |
| $\sigma_{3,1}$ | 0.02 | 0.38 | 0.00 | 2.35 | 1926 | 11773 |
| $\sigma_{3,2}$ | 0.22 | 7.62 | 0.00 | 40.52 | 3498 | 18611 |
| $\sigma_{3,4}$ | 0.91 | 0.39 | 0.41 | 2.04 | 43 | 179 |
| $\sigma_{4,1}$ | 0.55 | 0.18 | 0.29 | 1.04 | 34 | 138 |
| $\sigma_{4,2}$ | 1.14 | 0.40 | 0.58 | 2.24 | 36 | 146 |
| $\sigma_{4,3}$ | 0.79 | 0.32 | 0.37 | 1.68 | 40 | 166 |

Although I have found new evidence of individual variation in the state transition probabilities, accounting for this heterogeneity made little impact on state assignment and estimated activity budgets. Consistent with my simulation results (see Fig. 2), Viterbi-decoded states for the null and TMB models were in agreement 94% of the time. However, while “foraging” and “crowded” state assignments were largely unchanged, estimated overall activity budgets changed slightly for the “exploratory” state (36% for null, 33% for TMB) and the “directed” state (36% for null, 40% for TMB).

I do not intend to be critical of Isojunno *et al*. (2017) for limiting their candidate model set to finite mixtures or failing to achieve convergence for these complex models. They focused on discrete random effect models presumably because maximum likelihood inference for HMMs with continuous random effects has historically been very difficult (e.g. Altman 2007; Langrock *et al*. 2012; Schliehe-Diecks *et al*. 2012). Discrete random effects have been promoted as a practical alternative for movement HMMs (e.g. McKellar *et al*. 2015; Towner *et al*. 2016; DeRuiter *et al*. 2017), and the potential for TMB (Kristensen *et al*. 2016) as a tool to overcome such problems has only recently begun to be recognized by movement ecologists (e.g. Auger-Méthé *et al*. 2017). False convergence to a local maximum is notoriously difficult to assess, and it was entirely due to luck that my random draws of starting values for the finite mixture models happened to converge to parameter estimates with higher likelihood. Finally, as state assignment and calculating activity budgets was the primary purpose of the mixed HMM analysis performed by Isojunno *et al*. (2017), their main results and conclusions would likely be largely unaltered if they were instead based on a model that accounted for this individual variation. However, this would not be the case had their primary objective been to quantify individual heterogeneity in state transition probabilities.

If the primary objective of this analysis had been to gain an understanding of heterogeneity in individual movement behaviour, it is not clear how one would proceed with such a complex random effects model. Interpreting generic individual variation is already difficult in simpler finite mixture models (e.g. Towner *et al*. 2016; DeRuiter *et al*. 2017), and it would be challenging to concoct a biological story explaining these various pieces of evidence for individual heterogeneity across *N* (*N* – 1) = 12 continuous random effects (Table 5) and *MN*^2^ = 240 state transition probabilities (Fig. 6).

The random effects could reflect unexplained population-level behavioural heterogeneity attributable to different animal personalities (e.g. Réale *et al*. 2007; Hertel *et al*. 2020), but they could also simply be an artifact of deployments of differing lengths being observed in different environmental and behavioural contexts (e.g. Towner *et al*. 2016; DeRuiter *et al*. 2017). This is very difficult to determine, and I will not attempt to do so here. However, this highlights the challenge of interpreting random effects in biological terms.

## 5 Discussion

I have investigated the benefits of accounting for individual variation in the hidden state process of HMMs from a practical perspective, with emphasis on data sets common to animal movement behaviour biotelemetry studies. While my simulations covered a wide range of scenarios, they were by no means exhaustive. For example, I did not investigate pathological or degenerate cases with “bathtub-shaped” distributions for the state transition probabilities and instead focused on less extreme forms of individual heterogeneity. While limiting *T_m_* ≤ 250 proved sufficient for demonstrating general patterns in model performance as a function of time series lengths, telemetry devices can of course produce much longer time series. For much larger data sets (e.g. *M* ≫ 100 and *T_m_* ≫ 250), parameter estimates from the data-generating model can be expected to exhibit reduced bias and increased precision relative to the scenarios examined here. I did not examine sampling scenarios more typical of capture-recapture (Pradel 2005) or species occurrence (Gimenez *et al*. 2014) HMMs, which often involve a larger number of individuals and shorter time series than the scenarios examined here. I also limited my study to maximum likelihood inference via direct numerical maximization of the likelihood, although I believe similar patterns would emerge using expectation-maximization (e.g. Altman 2007) or Bayesian analysis methods (e.g. Turek *et al*. 2016).

I focused on individual-level random effects in the hidden state process because these have received the most attention in the movement modelling literature so far (e.g. McKellar *et al*. 2015; Towner *et al*. 2016; DeRuiter *et al*. 2017; Isojunno *et al*. 2017). Importantly, I have not examined individual effects on the (state-dependent) observation distribution parameters and would not necessarily expect the same patterns to emerge from a similar investigation of individual variation in the observation process. While this also warrants further investigation, there is already some evidence for the importance of accounting for individual variation in the state-dependent distributions (e.g. Altman 2007; Langrock *et al*. 2012; Schliehe-Diecks *et al*. 2012; McClintock *et al*. 2013; Rueda *et al*. 2013; Carter *et al*. 2020). In particular, I would expect unexplained individual variation in the observation process to be more consequential for state assignment.

HMMs tend to perform better as serial dependence in the data increases, particularly when state-dependent observation distributions overlap. Thus there may be specific conditions (e.g. “much” overlap with much stronger serial dependence) where discrete random effects may perform better as an approximation for continuous variation. However, the consequences of this model misspecification will be dependent on the form of heterogeneity, the degree of overlap, the amount of serial dependence, and other qualities of the data. I showed that this approximation is not generally robust, and, in practice, it may be very difficult to determine if it is reliable when truth is unknown.

The simulations and case study have highlighted some important considerations for practitioners contemplating the inclusion of individual random effects in their own analyses. As the results have provided much to digest, I break down these considerations under the following themes: 1) When to account for individual variation?; 2) How to account for individual variation?; 3) Is there evidence of individual variation?; and 4) How to interpret individual variation?

### 5.1 When to account for individual variation?

Accounting for generic individual variation comes at significant cost in terms of implementation and computation, particularly for random effects. When weighing the costs and benefits of random effects, three primary considerations are the study objectives, sample size, and model complexity. If the objective is strictly state assignment, then the inclusion of individual effects on the hidden state process makes little difference in terms of inference. Many movement HMM applications primarily focus on state assignment for inferences about behaviour, activity budgets, and/or resource selection (e.g. Beyer *et al*. 2013; Roever *et al*. 2014; Pirotta *et al*. 2018), with little emphasis on quantifying or understanding heterogeneity in state transition probabilities. Under these circumstances, random effects in the hidden state process simply may not be worth the additional effort.

If inference about individual variation in state transition probabilities is a primary objective, then random effect models should certainly be explored. When properly specified and fitted, individual random effects never hurt and tend to decrease bias and increase coverage (relative to the null model) and increase precision (relative to the fixed effects model). Using random effects to account for unexplained individual variation can also improve our ability to reliably estimate the effects of measurable covariates on the hidden state process. However, it’s important to consider that the feasibility and performance of a given mixed HMM will depend on model complexity and the amount of information contained in the data, as well as other factors that are typically out of the control of researchers, such as the degree of state-dependent distribution overlap (where less is generally better) and state persistence (where more is generally better). My results suggest that mixed HMMs generally don’t perform that well (and can be challenging to fit) with relatively few individuals and short time series, and these issues will only be exacerbated when *N* > 2. As *N* increases, the likelihood of observing all state transitions for individuals with very short time series decreases, thereby potentially making the estimation less reliable. With smaller sample sizes typical of animal biotelemetry studies (e.g. *M* < 50 and *T_m_* < 250), inferences about individual heterogeneity based on continuous random effects will tend to be weak. This is due to poor precision for the random effects variance parameters, which also tend to be underestimated with smaller sample sizes. When designing studies, researchers interested in applying continuous random effect models should consider allocating additional resources to maximise *M* (and, to a lesser degree, *T_m_*).

### 5.2 How to account for individual variation?

Continuous random effect models (BW and TMB) proved more robust to the underlying form of individual heterogeneity than null or discrete random effect models. The approximate approach of BW generally performed as well as (or better than) TMB when there was little overlap in the observation distributions, but the TMB model proved more robust to higher degrees of overlap. Thus if custom coding a continuous random effects model using `TMB` is beyond the skill set of a practitioner, the BW approach can be a reliable alternative in limited cases with very distinct observation distributions. This often applies to animal movement HMMs describing very different modes of movement (e.g. “foraging” and “transit”), but the degree of overlap should be investigated (e.g. Beyer *et al*. 2013) before proceeding with the BW approach. As demonstrated in the long-finned pilot whale example, BW is also of limited utility due to boundary issues that are more likely to occur as the number of states increases, the number of individuals increases, the lengths of time series decrease, and the degree of overlap increases. The TMB model does not suffer from these limitations, and I generally found the Laplace approximation as implemented in TMB to perform reasonably well across all HMM scenarios examined.

Discrete random effects are generally the best option only when individual heterogeneity is attributable to unmeasured categorical factors. While many categorical factors (e.g. sex, age class) can often be measured when deploying telemetry tags, others such as disease or breeding status often cannot. I do not advise using discrete random effects to account for continuous individual variation as this can underestimate uncertainty in state transition probabilities and lead to spurious inferences about covariate effects. If inference about individual variation in state transition probabilities is the primary objective, then investigating both discrete and continuous random effect models appears to be worth the effort for potential gains in terms of parameter estimation and state assignment. Yet care must be taken when fitting these complicated models. As illustrated by the long-finned pilot whale example, false convergence of random effect models can be particularly problematic as the number of states increases and sample sizes decrease.

Individual covariates are arguably the best way to account for (and learn about) potential factors driving individual heterogeneity in the hidden state process. Relative to random effect models, covariate models are also much easier to implement using existing software (Table 6). When designing telemetry studies, careful thought should be put towards identifying and collecting any measurable individual covariates that may be informative. When available covariates do not sufficiently explain individual heterogeneity, then random effects are certainly worth pursuing as a more parsimonious alternative to individual fixed effects. Discrete random effects and the approximate continuous random effect approach of Burnham & White (2002) can be implemented using the R package `momentuHMM` (McClintock & Michelot 2018). If the user is familiar enough with the `C++` template to custom code HMMs from scratch, then `TMB` (Kristensen *et al*. 2016) can be used to implement any of the models. `TMB` will also often be faster than optimization routines that rely on numerical differentiation.

**Table 6.** R packages for fitting HMMs with individual variation in the hidden state process using maximum likelihood methods.

| Package | Individual covariates | Random effects |  | Reference |
| --- | --- | --- | --- | --- |
|  |  | Discrete | Continuous |  |
| moveHMM | ✓ |  |  | Michelot *et al.* (2016) |
| msm | ✓ |  |  | Jackson (2011) |
| depmixS4 | ✓ |  |  | Visser & Speekenbrink (2010) |
| momentuHMM | ✓ | ✓ | ✓<sup>†</sup> | McClintock & Michelot (2018) |
| TMB | ✓ | ✓ | ✓ | Kristensen *et al.* (2016) |

<sup>†</sup> Continuous random effects are limited to the approximate approach of Burnham & White (2002).

### 5.3 Is there evidence of individual variation?

There are several ways to evaluate the strength of evidence for individual variation. These can include null hypothesis tests for the estimated coefficients of individual effects (for covariate, fixed, or discrete random effect models) or random effect variance estimates (for continuous random effect models). Estimated coefficients or random effect variances that are significantly different from zero typically indicate evidence of individual variation. Model selection criteria such as AIC or BIC can also be used for selecting or averaging models from a candidate set. I found the conditional AIC to work well for selecting among null, fixed, finite mixture, and BW models when sample sizes were larger and the observation distributions more distinct. The marginal AIC also worked reasonably well for selecting among the null, finite mixture, and TMB models under these scenarios, but it was conservative relative to the null likelihood ratio test. If inference about individual variation in the hidden state process is the primary objective and the exact form of any unexplained heterogeneity is unknown, my results suggest that candidate model sets should include models with both discrete and continuous random effects. This was evident in the long-finned pilot whale example, where I found stronger evidence for continuous random effects than for finite mixture models.

### 5.4 How to interpret individual variation?

Interpretation of measured individual covariate effects is straightforward, but it can be difficult to interpret generic individual-level effects. Discrete random effect models are sensitive to small sample sizes and can tend to identify spurious “behavioural contexts” that are an artifact of shorter time series (e.g. Towner *et al*. 2016; DeRuiter *et al*. 2017). In my simulation study, I also found individual fixed effect models to be susceptible to these small sample issues. Care should therefore be taken to avoid overinterpretation of individual fixed effects or finite mixture distributions. Continuous random effect models will tend to be less susceptible to small sample issues because they will shrink effect sizes for individuals with shorter time series toward the population mean. However, as demonstrated in the long-finned pilot whale example, this makes continuous random effects no less difficult to interpret in biological terms. Generic individual random effects only indicate evidence of individual variation in the hidden state process, but from evolutionary theory we already understand that biological parameters must in reality vary across individuals. Perhaps the most useful inference from evidence of generic individual heterogeneity is that there remains a need to identify and collect more informative covariates that can help explain the drivers of the underlying variation in the hidden state process.

## Acknowledgments

P. Conn, S. DeRuiter, S. Isojunno, D. Johnson, R. Langrock, and T. Michelot for helpful discussions. The findings and conclusions in the manuscript are those of the author(s) and do not necessarily represent the views of the National Marine Fisheries Service, NOAA. Any use of trade, product, or firm names does not imply an endorsement by the US Government.

