## Supplementary Material for "Worth the effort? A practical examination of random effects in hidden Markov models for animal telemetry data"

Brett T. McClintock

Marine Mammal Laboratory
Alaska Fisheries Science Center
NOAA National Marine Fisheries Service
Seattle, U.S.A.

January 5, 2021

### List of Figures

S1 State transition probability distributions for $\gamma_{m,1,2}$ and $\gamma_{m,2,1}$ with "moderate" ($\sigma_{1,2} = \sigma_{2,1} = 0.202$; solid lines) and "high" ($\sigma_{1,2} = \sigma_{2,1} = 0.416$; dashed lines) individual heterogeneity, where $\text{logit}^{-1}(\mu_{1,2}) = \text{logit}^{-1}(\mu_{2,1}) = 0.5$ ("lower" state persistence; black lines) or $\text{logit}^{-1}(\mu_{1,2}) = \text{logit}^{-1}(\mu_{2,1}) = 0.25$ ("higher" state persistence; red lines). . . . . . . 12

S2 State transition probabilities as a function of the covariate ($x_m$) for $\text{logit}^{-1}(\mu_{0,1,2}) = \text{logit}^{-1}(\mu_{0,2,1}) = 0.5$ ("lower" state persistence; black lines) and $\text{logit}^{-1}(\mu_{0,1,2}) = \text{logit}^{-1}(\mu_{0,2,1}) = 0.25$ ("higher" state persistence; red lines), where $\mu_{1,1,2} = 0.5$ and $\mu_{1,2,1} = -0.5$. From left to right, vertical blue lines respectively indicate standard normal 0.01, 0.025, 0.5, 0.975, and 0.99 quantiles. . . . . . . . . . . . . . . . . . . . . 13

S3 Nested loop plots for the proportion of Viterbi-decoded states that were correctly classified after accounting for chance agreement (top row) and the proportion of estimated state probabilities in which the true state received at least 0.50 (middle row) or 0.20 (bottom row) probability from 135 simulated scenarios without covariates that included "higher" (left column) or "lower" (right column) state persistence. Scenarios are ordered from outer to inner loops by the degree of state-dependent distribution overlap ("Overlap"), individual heterogeneity ("Sigma"), number of individuals ("nbInds"), and length of time series ("nbObs"). Comparisons are for the null (light green), fixed (dark green), mix2 (pink), mix3 (light orange), mix 4 (dark orange), BW (blue), and TMB (purple) models. . . . . . . . . . . . . . . 19

S4 Nested loop plots for mean bias (top row), 95% confidence interval coverage (middle row), and standard error (SE; bottom row) for $\gamma_{m,1,2}$ and $\gamma_{m,2,1}$ from 135 simulated scenarios without covariates. . . . . . . 22

S5 Nested loop plots for mean bias (top row), 95% confidence interval coverage (middle row), and standard error (SE; bottom row) for $\gamma_{m,1,2}$ and $\gamma_{m,2,1}$ from simulated scenarios without covariates, including 135 scenarios with "higher" state persistence (left column) and 90 scenarios with "mixA" or "mixB" finite mixtures (right column). Comparisons are for the AIC$_c$ model-averaged modMix (light orange), modFix (dark orange), modBW (light blue), and modTMB (light purple) models. 23

S6 Nested loop plots for mean bias (top row), 95% confidence interval coverage (middle row), and standard error (SE; bottom row) for $\gamma_{m,1,2}$ and $\gamma_{m,2,1}$ from 135 simulated scenarios without covariates. . . . . . . 24

S7 Nested loop plots for mean percent relative bias (top row), 95% confidence interval coverage (middle row), and percent standard error (SE; bottom row) for $\mu_1^y$ (left column) and $\mu_2^y$ (right column) from 225 simulated scenarios without covariate effects. Continuous random effect scenarios are limited to those with "higher" state persistence. . . . . 29
S8 Nested loop plots for mean percent relative bias (top row), 95% confidence interval coverage (middle row), and percent standard error (SE; bottom row) for $\mu_1^y$ from 135 simulated scenarios without covariate effects. . . . . 30
S9 Nested loop plots for mean percent relative bias (top row), 95% confidence interval coverage (middle row), and percent standard error (SE; bottom row) for $\mu_2^y$ from 135 simulated without covariate effects. . . . 31
S10 Nested loop plots for mean percent relative bias (top row), 95% confidence interval coverage (middle row), and percent standard error (SE; bottom row) for $\sigma_1^y$ (left column) and $\sigma_2^y$ (right column) from 225 simulated scenarios without covariate effects. Continuous random effect scenarios are limited to those with "higher" state persistence. . . . . 32
S11 Nested loop plots for mean percent relative bias (top row), 95% confidence interval coverage (middle row), and percent standard error (SE; bottom row) for $\sigma_1^y$ from 135 simulated scenarios without covariate effects. . . . . 33
S12 Nested loop plots for mean percent relative bias (top row), 95% confidence interval coverage (middle row), and percent standard error (SE; bottom row) for $\sigma_2^y$ from 135 simulated scenarios without covariate effects. . . . . 34
S13 Nested loop plots for mean $\text{AIC}_c$ weights in candidate model sets from simulated scenarios without covariates, including 135 scenarios with "higher" state persistence (left column) and 90 scenarios with "mixA" or "mixB" finite mixtures (right column). . . . . 35

S14 Nested loop plots for mean $AIC_c$ weights in candidate model sets from 135 simulated scenarios without covariates. . . . . . . . . . . . . . . . . . 36
S15 Nested loop plots for mean $AIC_c$ weights based on the "modTMB" candidate model set (top row) and proportion of likelihood ratio tests (LRT) supporting TMB over the null model (bottom row) from simulated scenarios without covariates, including 135 scenarios with "higher" state persistence (left column) and 90 scenarios with "mixA" or "mixB" finite mixtures (right column). . . . . . . . . . . . . . . . . . . . . . . . 37
S16 Nested loop plots for mean $AIC_c$ weights based on the "modTMB" candidate model set (top row) and proportion of likelihood ratio tests (LRT) supporting TMB over the null model (bottom row) from 135 simulated scenarios without covariates. . . . . . . . . . . . . . . . . . . 38
S17 Nested loop plots for the proportion of Viterbi-decoded states that were correctly classified after accounting for chance agreement (top) and the proportion of estimated state probabilities in which the true state received at least 0.50 (middle) or 0.20 (bottom) probability from 270 simulated scenarios with covariate effects $\mu_{1,1,2} = \mu_{1,2,1} = 0$ (left column) or $\mu_{1,1,2} = 0.5, \mu_{1,2,1} = -0.5$ (right column). Scenarios are ordered from outer to inner loops by state-dependent distribution overlap, state persistence, individual heterogeneity ("Sigma"), number of individuals ("nbInds"), and length of time series ("nbObs"). Dashed lines and "c" affix indicate models that included covariate effects. . . 41
S18 Nested loop plots for mean bias (top row), 95% confidence interval coverage (middle row), and confidence interval length (bottom row) for $\sigma_{1,2}$ and $\sigma_{2,1}$ from 270 simulated scenarios with covariate effects. Comparisons are for the BW (blue), BW including covariate effects ("BWc"; light blue), TMB (purple), and TMB including covariate effects ("TMBc"; light purple). . . . . . . . . . . . . . . . . . . . . . . 42

S19 Nested loop plots for mean bias (top row), 95% confidence interval coverage (middle row), and standard error (SE; bottom row) for $\gamma_{m,1,2}$ and $\gamma_{m,2,1}$ from 270 simulated scenarios with covariate effects. . . . . 43
S20 Nested loop plots for mean bias (top row), 95% confidence interval coverage (middle row), and standard error (SE; bottom row) for $\gamma_{m,1,2}$ and $\gamma_{m,2,1}$ from 270 simulated scenarios with covariate effects. . . . . 44
S21 Nested loop plots for mean percent relative bias (top row), 95% confidence interval coverage (middle row), and percent standard error (SE; bottom row) for $\mu_1^y$ from 270 simulated scenarios with covariate effects. 45
S22 Nested loop plots for mean percent relative bias (top row), 95% confidence interval coverage (middle row), and percent standard error (SE; bottom row) for $\mu_2^y$ from 270 simulated scenarios with covariate effects. 46
S23 Nested loop plots for mean percent relative bias (top row), 95% confidence interval coverage (middle row), and percent standard error (SE; bottom row) for $\sigma_1^y$ from 270 simulated scenarios with covariate effects. 47
S24 Nested loop plots for mean percent relative bias (top row), 95% confidence interval coverage (middle row), and percent standard error (SE; bottom row) for $\sigma_2^y$ from 270 simulated scenarios with covariate effects. 48
S25 Nested loop plots for mean $\text{AIC}_c$ weights in candidate model sets from 270 simulated scenarios with covariate effects. . . . . . . . . . . . . . 49
S26 Nested loop plots for median bias (top row), mean 95% confidence interval coverage (middle row), and median standard error (SE; bottom row) for covariate effects $\mu_{1,1,2}$ and $\mu_{1,2,1}$ from 270 simulated scenarios. Comparisons are for the null (light green), mix2 (pink), mix3 (light orange), mix4 (dark orange), BW (blue), and TMB (purple) models. 52

S27 Nested loop plots for median bias (top row), mean 95% confidence interval coverage (middle row), and median standard error (SE; bottom row) for covariate effects $\mu_{1,1,2}$ and $\mu_{1,2,1}$ from 270 simulated scenarios. Comparisons are for the AIC$_c$ model-averaged modMix (light orange), modBW (light blue), and modTMB (light purple) models. . . . . . . 53

S8 Overall mean proportion of Viterbi-decoded states that were correctly classified after accounting for chance agreement and the proportion of estimated state probabilities in which the true state received at least 0.50 or 0.20 probability by simulation design points for state-dependent distribution overlap, state persistence, covariate effects $(\mu_{1,1,2}, \mu_{1,2,1})$, individual heterogeneity $(\boldsymbol{\sigma})$, number of individuals, and time series length from 540 scenarios with covariate effects. All models (except "fixed") included terms for the covariate effects. . . . . . . . . . . . . 40

S9 Overall mean bias, 95% confidence interval coverage, and standard error (SE) for covariate effect $\mu_{1,1,2}$ by simulation design points for the covariate effect, state-dependent distribution overlap, state persistence, individual heterogeneity ($\boldsymbol{\sigma}$), number of individuals, and time series length from 540 scenarios with covariate effects. . . . . . . . . . 50
S10 Overall mean bias, 95% confidence interval coverage, and standard error (SE) for covariate effect $\mu_{1,2,1}$ by simulation design points for the covariate effect, state-dependent distribution overlap, state persistence, individual heterogeneity ($\boldsymbol{\sigma}$), number of individuals, and time series length from 540 scenarios with covariate effects. . . . . . . . . . 51

### S1 Approach of Burnham & White (2002)

Here I briefly describe the approximate approach of Burnham & White (2002) in the context of continuous individual-level random effects for the state transition probabilities in a $N$-state HMM (Eq. 4 in main text). Let $\hat{\boldsymbol{\beta}}_{i,j} = \left(\hat{\beta}_{1,i,j}, \ldots, \hat{\beta}_{M,i,j}\right)$ denote the maximum likelihood estimates for the (logit scale) state transition probability parameters from the individual fixed effects model (Eq. 2) corresponding to switches from state $i$ to state $j$. These can be represented as $\hat{\boldsymbol{\beta}}_{i,j} = \mathbf{z}_{i,j} + \boldsymbol{\epsilon}_{i,j}$, where the stochastic component $\boldsymbol{\epsilon}_{i,j} = (\epsilon_{1,i,j}, \ldots, \epsilon_{M,i,j})$ has unconditional (on $\mathbf{z}_{i,j}$) sampling variance-covariance matrix $\mathbf{W}_{i,j}$. From generalized least squares theory, for given $\sigma^2_{i,j}$, we have

$$\mu_{i,j} = \left(\mathbf{x}'\mathbf{D}^{-1}_{i,j}\mathbf{x}\right)^{-1}\mathbf{x}'\mathbf{D}^{-1}_{i,j}\hat{\boldsymbol{\beta}}_{i,j}, \tag{S1}$$

where $\mathbf{x}$ is a column vector of $M$ ones, $\mathbf{D}_{i,j} = \sigma^2_{i,j}\mathbf{I} + \mathbf{W}_{i,j}$, and $\mathbf{I}$ is a $M \times M$ identity matrix. Assuming (approximate) normality of $\hat{\boldsymbol{\beta}}_{i,j}$, we can obtain an estimate for $\sigma^2_{i,j}$ by numerically solving the equation

$$M - 1 = \left(\hat{\boldsymbol{\beta}}_{i,j} - \mathbf{x}\mu_{i,j}\right)'\mathbf{D}^{-1}_{i,j}\left(\hat{\boldsymbol{\beta}}_{i,j} - \mathbf{x}\mu_{i,j}\right), \tag{S2}$$

and the shrinkage estimator of $\mathbf{z}_{i,j}$ is

$$\tilde{\mathbf{z}}_{i,j} = \mathbf{H}_{i,j}\left(\hat{\boldsymbol{\beta}}_{i,j} - \mathbf{x}\mu_{i,j}\right) + \mathbf{x}\mu_{i,j} \tag{S3}$$

evaluated at the solution $\hat{\sigma}^2_{i,j}$ from Eq. S2, where $\mathbf{H}_{i,j} = \sigma_{i,j}\mathbf{D}^{-1/2}_{i,j}$. An equivalent formula, $\tilde{\mathbf{z}}_{i,j} = \mathbf{G}_{i,j}\hat{\boldsymbol{\beta}}_{i,j}$, is based on the projection matrix $\mathbf{G}_{i,j} = \mathbf{H}_{i,j} + (\mathbf{I} - \mathbf{H}_{i,j})\,\mathbf{A}_{i,j}\mathbf{D}^{-1}_{i,j}$, where $\mathbf{A}_{i,j} = \mathbf{x}\left(\mathbf{x}'\mathbf{D}^{-1}_{i,j}\mathbf{x}\right)^{-1}\mathbf{x}'$. To include $q$ individual covariate effects, we simply expand $\mathbf{x}$ and $\mu_{i,j}$ in Eqs S1–S3 to a $M \times q$ design matrix ($\mathbf{X}$) and $\boldsymbol{\mu}_{i,j} = (\mu_{0,i,j}, \mu_{1,i,j}, \ldots, \mu_{q,i,j})$, respectively, with $M - q$ degrees of freedom on the left-hand side of Eq. S2.

This approach is approximate because $\mathbf{W}_{i,j}$ is not known, and the estimated variance-covariance matrix for $\boldsymbol{\beta}_{i,j}$ ($\hat{\mathbf{W}}_{i,j}$) from the fixed effects model is used in place of $\mathbf{W}_{i,j}$. After this procedure is completed for all $N(N-1)$ random effects, a bias-corrected conditional Akaike's Information Criterion (AIC$_c$; Burnham & Anderson 2002) can be calculated as $-2\log\tilde{\mathcal{L}}_{fix} + 2p + 2\frac{p(p+1)}{n-p-1}$, where $\tilde{\mathcal{L}}_{fix}$ is the value of the re-optimized likelihood (Eq. 2) with the state transition probabilities fixed to $\tilde{\boldsymbol{\Gamma}}_m = (\tilde{\gamma}_{m,i,j})$, $\tilde{\gamma}_{m,i,j} = \frac{\exp(\tilde{z}_{m,i,j})}{\sum_{l=1}^{N}\exp\left(\tilde{z}_{m,i,l}\right)}$, $\tilde{z}_{m,i,j} = 0$ for $i = j$, $p = \sum_{i=1}^{N}\sum_{j\neq i}\mathrm{tr}(\mathbf{G}_{i,j}) + r$, $r$ is the number of free parameters that were re-optimized at $\tilde{\mathbf{\Gamma}}_m$, and $n$ is the sample size. If the random effects model is better supported than null or fixed effects models, then $\hat{\mu}_{i,j}$, $\hat{\sigma}_{i,j}$, $\hat{\tilde{\mathbf{z}}}_{i,j}$, and $\hat{\tilde{\gamma}}_{m,i,j}$ can be used for inferences about state transition probabilities, but inferences about any other model parameters can be drawn from the fixed effects model (because the estimated variance-covariance matrix for the re-optimized likelihood is not valid). One known issue with this approach is that $\hat{\mathbf{W}}_{i,j}$ is unreliable when any $\hat{\gamma}_{m,i,j}$ from the fixed effects model is near 0 or 1, and I suspect that such boundary issues will become more likely as $M$ increases, $T_m$ decreases, and $N$ increases. Bayesian analogues to the two-stage approach of Burnham & White (2002) could be less susceptible to these boundary issues (Hooten *et al.* 2016, 2019).

### S2 Simulatation study

#### S2.1 Simulation methods

##### S2.1.1 Continuous-valued random effects

Continuous-valued random effects with "moderate" heterogeneity were simulated with $\sigma_{1,2} = \sigma_{2,1} = 0.202$ and $\text{logit}^{-1}(\mu_{1,2}) = \text{logit}^{-1}(\mu_{2,1}) \in \{0.5, 0.25\}$, corresponding to a mean of 0.5 and standard deviation of 0.05 on the state transition probability scale when $\text{logit}^{-1}(\mu_{1,2}) = \text{logit}^{-1}(\mu_{2,1}) = 0.5$. Those with "high" heterogeneity were simulated with $\sigma_{1,2} = \sigma_{2,1} = 0.416$ and $\text{logit}^{-1}(\mu_{1,2}) = \text{logit}^{-1}(\mu_{2,1}) \in \{0.5, 0.25\}$, corresponding to a mean of 0.5 and standard deviation of 0.10 on the state transition probability scale when $\text{logit}^{-1}(\mu_{1,2}) = \text{logit}^{-1}(\mu_{2,1}) = 0.5$ (Fig. S1).

##### S2.1.2 Continuous-valued covariate effects

For simulations including measurable covariate effects, two covariate scenarios were included: 1) $\text{logit}^{-1}(\mu_{0,1,2}) = \text{logit}^{-1}(\mu_{0,2,1}) \in \{0.5, 0.25\}$, $\mu_{1,1,2} = 0$, and $\mu_{1,2,1} = 0$; and 2) $\text{logit}^{-1}(\mu_{0,1,2}) = \text{logit}^{-1}(\mu_{0,2,1}) \in \{0.5, 0.25\}$, $\mu_{1,1,2} = 0.5$, and $\mu_{1,2,1} = -0.5$, where $z_{m,i,j} \overset{\text{iid}}{\sim} \mathcal{N}\left(\mu_{0,i,j} + x_m \mu_{1,i,j}, \sigma^2_{i,j}\right)$ for $i \neq j$, $z_{m,i,j} = 0$ for $i = j$, $x_m$ is a measurable individual covariate drawn from a standard normal distribution, and $\gamma_{m,i,j} = \frac{\exp(z_{m,i,j})}{\sum_{l=1}^{N} \exp\left(z_{m,i,l}\right)}$ (Fig. S2).

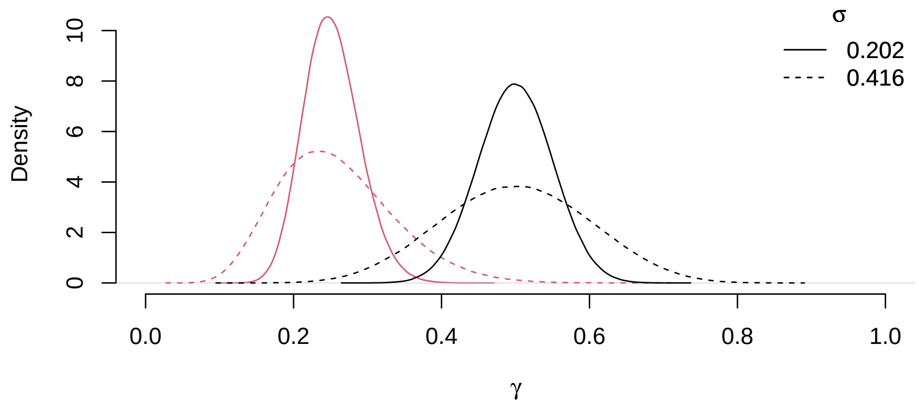

***Figure S1.*** *State transition probability distributions for* $\gamma_{m,1,2}$ *and* $\gamma_{m,2,1}$ *with "moderate"* ($\sigma_{1,2} = \sigma_{2,1} = 0.202$*; solid lines) and "high"* ($\sigma_{1,2} = \sigma_{2,1} = 0.416$*; dashed lines) individual heterogeneity, where* $logit^{-1}(\mu_{1,2}) = logit^{-1}(\mu_{2,1}) = 0.5$ *("lower" state persistence; black lines) or* $logit^{-1}(\mu_{1,2}) = logit^{-1}(\mu_{2,1}) = 0.25$ *("higher" state persistence; red lines).*

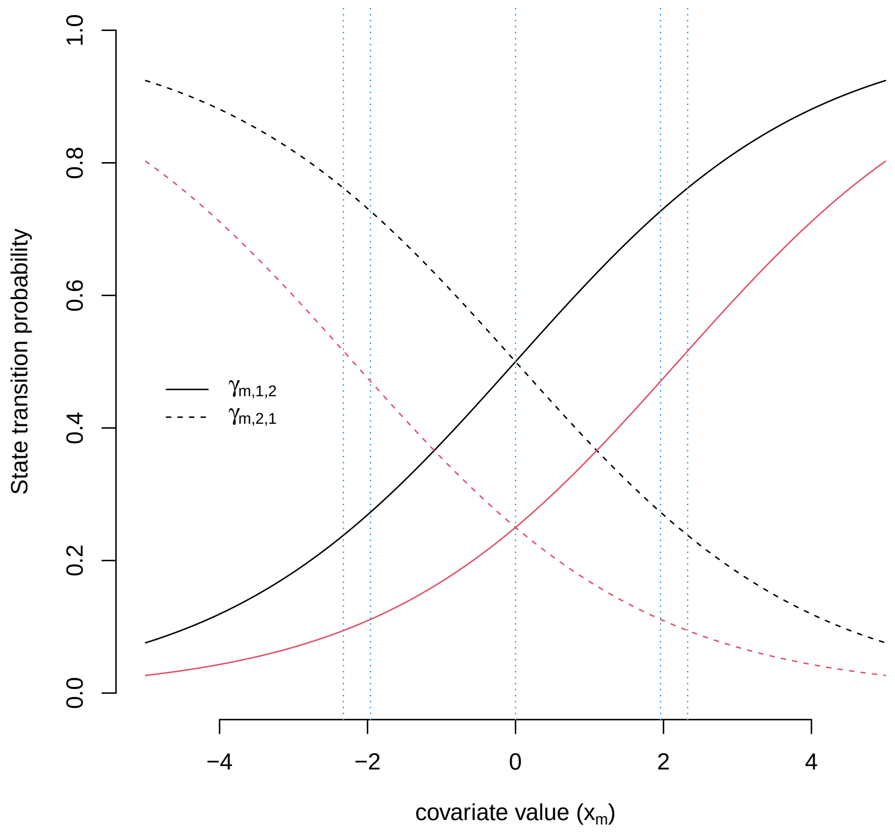

***Figure S2.*** *State transition probabilities as a function of the covariate* $(x_m)$ *for* $logit^{-1}(\mu_{0,1,2}) = logit^{-1}(\mu_{0,2,1}) = 0.5$ *("lower" state persistence; black lines) and* $logit^{-1}(\mu_{0,1,2}) = logit^{-1}(\mu_{0,2,1}) = 0.25$ *("higher" state persistence; red lines), where* $\mu_{1,1,2} = 0.5$ *and* $\mu_{1,2,1} = -0.5$*. From left to right, vertical blue lines respectively indicate standard normal 0.01, 0.025, 0.5, 0.975, and 0.99 quantiles.*

##### S2.1.3 Derivations of $\gamma_{m,i,j}$ and $\mu_{1,i,j}$ for finite mixture models

To facilitate comparisons across models, individual-level state transition probabilities were derived from the maximum likelihood estimates for the finite mixture models as $\gamma_{m,i,j} = \sum_{k=1}^{K} \omega_{m,k} \gamma_{i,j}^{(k)}$, where

$$\omega_{m,k} = \frac{\boldsymbol{\delta}^{(k)} \boldsymbol{\Gamma}^{(k)} \mathbf{P}(\mathbf{y}_{m,1}) \boldsymbol{\Gamma}^{(k)} \mathbf{P}(\mathbf{y}_{m,2}) \cdots \boldsymbol{\Gamma}^{(k)} \mathbf{P}(\mathbf{y}_{m,T_m-1}) \boldsymbol{\Gamma}^{(k)} \mathbf{P}(\mathbf{y}_{m,T_m}) \mathbf{1} \pi^{(k)}}{\sum_{l=1}^{K} \boldsymbol{\delta}^{(l)} \boldsymbol{\Gamma}^{(l)} \mathbf{P}(\mathbf{y}_{m,1}) \boldsymbol{\Gamma}^{(l)} \mathbf{P}(\mathbf{y}_{m,2}) \cdots \boldsymbol{\Gamma}^{(l)} \mathbf{P}(\mathbf{y}_{m,T_m-1}) \boldsymbol{\Gamma}^{(l)} \mathbf{P}(\mathbf{y}_{m,T_m}) \mathbf{1} \pi^{(l)}}$$

is the probability that individual $m$ belongs to mixture $k$. Population-level covariate effects were derived as $\mu_{1,i,j} = \sum_{k=1}^{K} \pi^{(k)} \mu_{1,i,j}^{(k)}$. Standard errors were derived using the delta method and finite-difference approximations of the first derivative for these transformations (e.g. Ver Hoef 2012).

##### S2.1.4 Modified forward-backward algorithm for finite mixture models

For a standard HMM, local state decoding involves both the forward probabilities

$$\alpha_{m,t}(j) = \Pr(\mathbf{y}_{m,1}, \ldots, \mathbf{y}_{m,t}, S_{m,t} = j)$$

and the backward probabilities

$$\beta_{m,t}(j) = \Pr(\mathbf{y}_{m,t+1}, \ldots, \mathbf{y}_{m,T_m} \mid S_{m,t} = j),$$

where

$$\boldsymbol{\alpha}_{m,t} = \boldsymbol{\alpha}_{m,t-1} \boldsymbol{\Gamma} \mathbf{P}(\mathbf{y}_{m,t}),$$

$$\boldsymbol{\beta}'_{m,t} = \boldsymbol{\Gamma} \mathbf{P}(\mathbf{y}_{m,t+1}) \boldsymbol{\beta}'_{m,t+1},$$

$\boldsymbol{\alpha}_{m,1} = \boldsymbol{\delta} \mathbf{P}(\mathbf{y}_{m,1})$, and $\boldsymbol{\beta}_{m,T_m} = \mathbf{1}$. Local decoding involves maximising the quantity

$$\Pr\left(S_{m,t} = s_{m,t} \mid \mathbf{y}_{m,1}, \ldots, \mathbf{y}_{m,T_m}\right) = \frac{\alpha_{m,t}(s_{m,t}) \beta_{m,t}(s_{m,t})}{\mathcal{L}_m}, \tag{S4}$$

with respect to $s_{m,t}$ for every individual $m$ and time $t = 1, \ldots, T_m$, where

$$\mathcal{L}_m = \boldsymbol{\delta} \boldsymbol{\Gamma} \mathbf{P}(\mathbf{y}_{m,1}) \boldsymbol{\Gamma} \mathbf{P}(\mathbf{y}_{m,2}) \cdots \boldsymbol{\Gamma} \mathbf{P}(\mathbf{y}_{m,T_m-1}) \boldsymbol{\Gamma} \mathbf{P}(\mathbf{y}_{m,T_m}) \mathbf{1}$$

is the likelihood for individual $m$. At any time point $t$, Eq. S4 can be evaluated for $S_{m,t} = 1, \ldots, N$, yielding the state probabilities for individual $m$.

With $K$ mixtures, the state probabilities for each mixture are weighted by the mixture probability for individual $m$:

$$\Pr\left(S_{m,t} = s_{m,t} \mid \mathbf{y}_{m,1}, \ldots, \mathbf{y}_{m,T_m}\right) = \sum_{k=1}^{K} \omega_{m,k} \frac{\alpha_{m,t}^{(k)}(s_{m,t}) \beta_{m,t}^{(k)}(s_{m,t})}{\mathcal{L}_m^{(k)}},$$

where $\boldsymbol{\alpha}_{m,t}^{(k)} = \boldsymbol{\alpha}_{m,t-1}^{(k)} \boldsymbol{\Gamma}^{(k)} \mathbf{P}(\mathbf{y}_{m,t})$, $\boldsymbol{\beta}_{m,t}^{(k)\prime} = \boldsymbol{\Gamma}^{(k)} \mathbf{P}(\mathbf{y}_{m,t+1}) \boldsymbol{\beta}_{m,t+1}^{(k)\prime}$, $\boldsymbol{\alpha}_{m,1}^{(k)} = \boldsymbol{\delta}^{(k)} \mathbf{P}(\mathbf{y}_{m,1})$, $\boldsymbol{\beta}_{m,T_m}^{(k)} = \mathbf{1}$, and

$$\mathcal{L}_m^{(k)} = \boldsymbol{\delta}^{(k)} \boldsymbol{\Gamma}^{(k)} \mathbf{P}(\mathbf{y}_{m,1}) \boldsymbol{\Gamma}^{(k)} \mathbf{P}(\mathbf{y}_{m,2}) \cdots \boldsymbol{\Gamma}^{(k)} \mathbf{P}(\mathbf{y}_{m,T_m-1}) \boldsymbol{\Gamma}^{(k)} \mathbf{P}(\mathbf{y}_{m,T_m}) \mathbf{1}.$$

##### S2.1.5 Modified Viterbi algorithm for finite mixture models

Let $q_{m,t}(j) = \Pr(S_{m,1} = s_{m,1}, \ldots, S_{m,t-1} = s_{m,t-1}, S_{m,t} = j, \mathbf{y}_{m,1}, \ldots, \mathbf{y}_{m,T_m})$ be the joint probability (with the data and the optimal state sequence up to time $t-1$) that the state for individual $m$ at time $t$ is $j$. The Viterbi algorithm for a standard HMM makes use of the following recurrence: $q_{m,t}(j) = \max_i \, q_{m,t-1}(i) \gamma_{i,j} f(\mathbf{y}_{m,t} \mid S_{m,t} = j)$, where $q_{m,1}(j) = \delta_j f(\mathbf{y}_{m,1} \mid S_{m,1} = j)$. The globally decoded states can then be determined by going backwards in time: choose $s_{m,T_m} = \operatorname{argmax}_j q_{m,T_m}(j)$ and then $s_{m,t} = \operatorname{argmax}_j q_{m,t}(j) \gamma_{j,s_{m,t+1}}$ for $t < T$.

With $K$ mixtures, I modified the Viterbi algorithm by weighting the initial distribution and state transitions probabilities for each mixture based on the mixture probabilities for individual $m$:

$$q_{m,t}(j) = \max_i \, q_{m,t-1}(i) \sum_{k=1}^{K} \omega_{m,k} \gamma_{i,j}^{(k)} f(\mathbf{y}_{m,t} \mid S_{m,t} = j),$$

where $q_{m,1}(j) = \sum_{k=1}^{K} \omega_{m,k} \delta_j^{(k)} f(\mathbf{y}_{m,1} \mid S_{m,1} = j)$ for $k = 1, \ldots, K$. The globally decoded states can then be determined by choosing

$$s_{m,t} = \begin{cases} \operatorname{argmax}_j q_{m,T_m}(j) & \text{if } t = T_m \\ \operatorname{argmax}_j q_{m,t}(j) \sum_{k=1}^{K} \omega_{m,k} \gamma_{j,s_{m,t+1}}^{(k)} & \text{if } 1 \leq t < T_m \end{cases}. \tag{S5}$$

An alternative approach kindly suggested by an anonymous reviewer is

$$s_{m,t} = s_{m,t}^{(l_m)},$$

where

$$s^{(k)}_{m,t} = \begin{cases} \text{argmax}_j \; q^{(k)}_{m,T_m}(j) & \text{if } t = T_m \\ \text{argmax}_j \; q^{(k)}_{m,t}(j)\, \gamma^{(k)}_{j,s_{m,t+1}} & \text{if } 1 \leq t < T_m \end{cases},$$

$$q^{(k)}_{m,t}(j) = \text{max}_i \; q^{(k)}_{m,t-1}(i)\gamma^{(k)}_{i,j} f(\mathbf{y}_{m,t} \mid S_{m,t} = j),$$

$q^{(k)}_{m,1}(j) = \delta^{(k)}_j f(\mathbf{y}_{m,1} \mid S_{m,1} = j)$, and

$$l_m = \underset{k}{\text{argmax}} \; \omega_{m,k} q^{(k)}_{m,T_m}.$$

While this alternative approach also worked well, it resulted in (marginally) fewer correct state assignments in 87% of the "without covariates" and 89% of the "with covariates" simulation scenarios. Hence I only report Viterbi-decoded state assignment results for finite mixture models based on Eq. S5, but the mean increase in the percentage of correct state assignments was only 0.12% (SD = 0.18%) in the "without covariates" and 0.11% (SD = 0.17%) in the "with covariates" scenarios.

#### S2.2 Simulation results

##### S2.2.1 Without covariates

***Table S1.*** *Overall mean proportion of Viterbi-decoded states that were correctly classified after accounting for chance agreement and the proportion of estimated state probabilities in which the true state received at least 0.50 or 0.20 probability by simulation design points for state-dependent distribution overlap, continuous or discrete random effects, individual heterogeneity, number of individuals, and time series length from 360 scenarios without covariate effects.*

| Model | Overlap | | | Random effects | | | | | | | No. individuals ($M$) | | | | | Time series length ($T_m$) | | | Overall |
|---|---|---|---|---|---|---|---|---|---|---|---|---|---|---|---|---|---|---|---|
| | | | | Continuous | | | | | Discrete | | | | | | | | | | |
| | | | | State persistence | | $\boldsymbol{\sigma}$ | | | | | | | | | | | | | |
| | Little | Some | Much | Higher | Lower | 0 | 0.202 | 0.416 | mixA | mixB | 100 | 50 | 30 | 15 | 5 | 250 | 110 | 30 - 250 | |
| | | | | | | | | | Prop. states correctly classified | | | | | | | | | | |
| null | 0.99 | 0.81 | 0.50 | 0.79 | 0.75 | 0.77 | 0.77 | 0.77 | 0.76 | 0.75 | 0.77 | 0.77 | 0.77 | 0.77 | 0.76 | 0.77 | 0.77 | 0.77 | 0.77 |
| fixed | 0.99 | 0.81 | 0.49 | 0.79 | 0.74 | 0.76 | 0.76 | 0.77 | 0.76 | 0.78 | 0.77 | 0.77 | 0.77 | 0.77 | 0.76 | 0.77 | 0.76 | 0.76 | 0.77 |
| mix2 | 0.99 | 0.82 | 0.51 | 0.79 | 0.75 | 0.77 | 0.77 | 0.77 | 0.77 | 0.79 | 0.78 | 0.78 | 0.78 | 0.77 | 0.76 | 0.78 | 0.77 | 0.77 | 0.77 |
| mix3 | 0.99 | 0.82 | 0.51 | 0.79 | 0.75 | 0.77 | 0.77 | 0.77 | 0.77 | 0.79 | 0.78 | 0.78 | 0.78 | 0.77 | 0.76 | 0.78 | 0.77 | 0.77 | 0.77 |
| mix4 | 0.99 | 0.82 | 0.51 | 0.79 | 0.75 | 0.77 | 0.77 | 0.77 | 0.77 | 0.79 | 0.78 | 0.78 | 0.78 | 0.77 | 0.76 | 0.78 | 0.77 | 0.77 | 0.77 |
| BW | 0.99 | 0.82 | 0.51 | 0.81 | 0.79 | 0.79 | 0.80 | 0.80 | 0.81 | 0.83 | 0.84 | 0.83 | 0.81 | 0.79 | 0.77 | 0.78 | 0.79 | 0.85 | 0.80 |
| TMB | 0.99 | 0.82 | 0.51 | 0.79 | 0.75 | 0.77 | 0.77 | 0.78 | 0.77 | 0.78 | 0.78 | 0.78 | 0.78 | 0.77 | 0.76 | 0.78 | 0.77 | 0.77 | 0.77 |
| | | | | | | | | | Prop. state probabilities > 0.50 | | | | | | | | | | |
| null | 1.00 | 0.91 | 0.76 | 0.90 | 0.88 | 0.89 | 0.89 | 0.89 | 0.88 | 0.88 | 0.89 | 0.89 | 0.89 | 0.89 | 0.88 | 0.89 | 0.89 | 0.89 | 0.89 |
| fixed | 1.00 | 0.91 | 0.75 | 0.90 | 0.87 | 0.88 | 0.88 | 0.89 | 0.89 | 0.89 | 0.89 | 0.89 | 0.89 | 0.89 | 0.88 | 0.89 | 0.88 | 0.89 | 0.89 |
| mix2 | 1.00 | 0.90 | 0.75 | 0.90 | 0.88 | 0.89 | 0.89 | 0.89 | 0.88 | 0.86 | 0.89 | 0.89 | 0.88 | 0.88 | 0.88 | 0.88 | 0.88 | 0.88 | 0.88 |
| mix3 | 1.00 | 0.90 | 0.75 | 0.90 | 0.88 | 0.89 | 0.89 | 0.89 | 0.88 | 0.86 | 0.89 | 0.89 | 0.88 | 0.88 | 0.88 | 0.88 | 0.88 | 0.88 | 0.88 |
| mix4 | 1.00 | 0.90 | 0.75 | 0.90 | 0.87 | 0.89 | 0.89 | 0.89 | 0.88 | 0.86 | 0.89 | 0.88 | 0.88 | 0.88 | 0.88 | 0.88 | 0.88 | 0.88 | 0.88 |
| BW | 1.00 | 0.91 | 0.76 | 0.91 | 0.89 | 0.90 | 0.90 | 0.90 | 0.91 | 0.92 | 0.92 | 0.91 | 0.91 | 0.90 | 0.89 | 0.89 | 0.90 | 0.93 | 0.90 |
| TMB | 1.00 | 0.91 | 0.76 | 0.90 | 0.88 | 0.89 | 0.89 | 0.89 | 0.89 | 0.89 | 0.89 | 0.89 | 0.89 | 0.89 | 0.88 | 0.89 | 0.89 | 0.89 | 0.89 |
| | | | | | | | | | Prop. state probabilities > 0.20 | | | | | | | | | | |
| null | 1.00 | 0.96 | 0.95 | 0.97 | 0.97 | 0.97 | 0.97 | 0.97 | 0.97 | 0.97 | 0.98 | 0.98 | 0.97 | 0.97 | 0.96 | 0.97 | 0.97 | 0.97 | 0.97 |
| fixed | 1.00 | 0.96 | 0.93 | 0.97 | 0.96 | 0.96 | 0.96 | 0.96 | 0.96 | 0.97 | 0.97 | 0.97 | 0.97 | 0.96 | 0.95 | 0.97 | 0.96 | 0.96 | 0.96 |
| mix2 | 1.00 | 0.96 | 0.95 | 0.97 | 0.97 | 0.97 | 0.97 | 0.97 | 0.97 | 0.96 | 0.97 | 0.97 | 0.97 | 0.97 | 0.96 | 0.97 | 0.97 | 0.97 | 0.97 |
| mix3 | 1.00 | 0.96 | 0.95 | 0.97 | 0.97 | 0.97 | 0.97 | 0.97 | 0.97 | 0.96 | 0.97 | 0.97 | 0.97 | 0.97 | 0.95 | 0.97 | 0.97 | 0.97 | 0.97 |
| mix4 | 1.00 | 0.96 | 0.94 | 0.97 | 0.97 | 0.97 | 0.97 | 0.97 | 0.97 | 0.96 | 0.97 | 0.97 | 0.97 | 0.97 | 0.95 | 0.97 | 0.97 | 0.97 | 0.97 |
| BW | 1.00 | 0.96 | 0.94 | 0.97 | 0.97 | 0.97 | 0.97 | 0.97 | 0.97 | 0.97 | 0.98 | 0.97 | 0.97 | 0.97 | 0.96 | 0.97 | 0.97 | 0.97 | 0.97 |
| TMB | 1.00 | 0.96 | 0.95 | 0.97 | 0.97 | 0.97 | 0.97 | 0.97 | 0.97 | 0.97 | 0.98 | 0.97 | 0.97 | 0.97 | 0.96 | 0.97 | 0.97 | 0.97 | 0.97 |

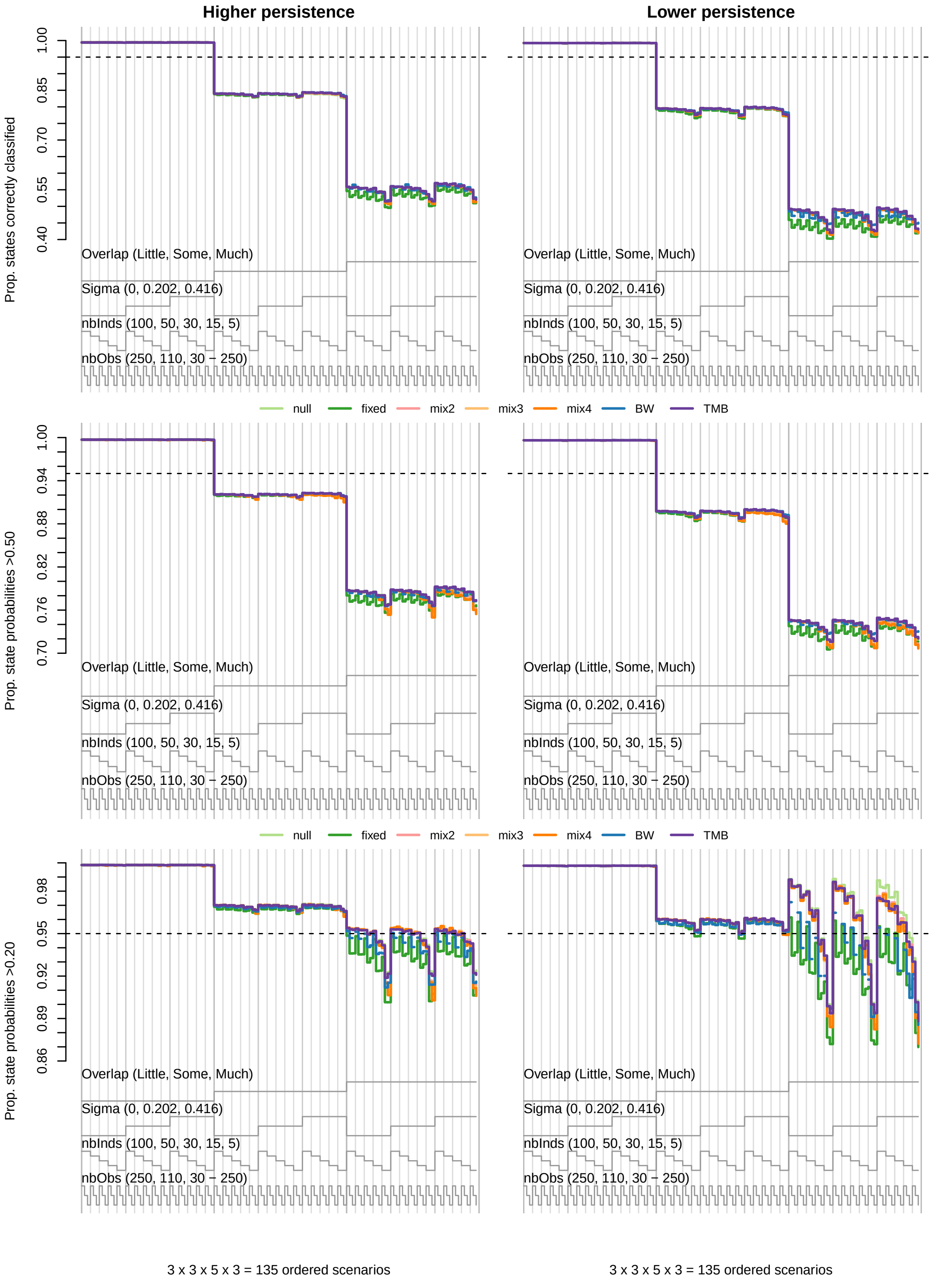

***Figure S3.*** *Nested loop plots for the proportion of Viterbi-decoded states that were correctly classified after accounting for chance agreement (top row) and the proportion of estimated state probabilities in which the true state received at least 0.50 (middle row) or 0.20 (bottom row) probability from 135 simulated scenarios without covariates that included "higher" (left column) or "lower" (right column) state persistence. Scenarios are ordered from outer to inner loops by the degree of state-dependent distribution overlap ("Overlap"), individual heterogeneity ("Sigma"), number of individuals ("nbInds"), and length of time series ("nbObs"). Comparisons are for the null (light green), fixed (dark green), mix2 (pink), mix3 (light orange), mix 4 (dark orange), BW (blue), and TMB (purple) models.*

***Table S2.*** *Overall mean bias, 95% confidence interval coverage, and standard error (SE) for $\gamma_{m,1,2}$ by simulation design points for state-dependent distribution overlap, continuous or discrete random effects, number of individuals, and time series length from 360 scenarios without covariate effects.*

| Model | Overlap | | | Random effects | | | | | | | No. individuals ($M$) | | | | | Time series length ($T_m$) | | | Overall |
|---|---|---|---|---|---|---|---|---|---|---|---|---|---|---|---|---|---|---|---|
| | | | | Continuous | | | | | Discrete | | | | | | | | | | |
| | | | | State persistence | | $\boldsymbol{\sigma}$ | | | | | | | | | | | | | |
| | Little | Some | Much | Higher | Lower | 0 | 0.202 | 0.416 | mixA | mixB | 100 | 50 | 30 | 15 | 5 | 250 | 110 | 30 - 250 | |
| | | | | | | | | | Bias | | | | | | | | | | |
| null | -0.01 | -0.01 | -0.02 | -0.01 | -0.01 | 0.00 | -0.01 | -0.02 | -0.05 | 0.00 | -0.01 | -0.01 | -0.01 | -0.01 | -0.02 | -0.01 | -0.01 | -0.01 | -0.01 |
| fixed | 0.01 | 0.01 | 0.01 | 0.02 | 0.00 | 0.01 | 0.01 | 0.01 | 0.01 | 0.00 | 0.01 | 0.01 | 0.01 | 0.01 | 0.00 | 0.00 | 0.01 | 0.02 | 0.01 |
| mix2 | 0.00 | 0.00 | -0.01 | 0.00 | -0.01 | 0.00 | 0.00 | -0.01 | 0.00 | 0.00 | 0.00 | 0.00 | 0.00 | 0.00 | -0.01 | 0.00 | 0.00 | 0.00 | 0.00 |
| modMix | 0.00 | 0.00 | -0.01 | 0.00 | -0.01 | 0.00 | 0.00 | -0.01 | 0.00 | 0.00 | 0.00 | 0.00 | 0.00 | 0.00 | -0.01 | 0.00 | 0.00 | 0.00 | 0.00 |
| modFix | 0.00 | 0.00 | -0.01 | 0.00 | -0.01 | 0.00 | 0.00 | 0.00 | 0.00 | 0.00 | 0.00 | 0.00 | 0.00 | 0.00 | -0.01 | 0.00 | 0.00 | 0.00 | 0.00 |
| BW | 0.00 | 0.01 | -0.05 | 0.01 | -0.02 | -0.01 | -0.01 | -0.01 | -0.01 | -0.01 | 0.00 | -0.01 | -0.01 | -0.01 | -0.02 | -0.01 | -0.01 | 0.00 | -0.01 |
| modBW | 0.00 | 0.00 | -0.02 | 0.00 | -0.01 | 0.00 | 0.00 | 0.00 | -0.01 | 0.00 | 0.00 | 0.00 | 0.00 | 0.00 | -0.01 | 0.00 | 0.00 | 0.00 | 0.00 |
| TMB | 0.00 | 0.00 | -0.01 | 0.00 | -0.01 | 0.00 | 0.00 | 0.00 | -0.01 | 0.00 | 0.00 | 0.00 | 0.00 | 0.00 | -0.01 | 0.00 | 0.00 | 0.00 | 0.00 |
| modTMB | 0.00 | 0.00 | -0.01 | 0.00 | -0.01 | 0.00 | 0.00 | -0.01 | 0.00 | 0.00 | 0.00 | 0.00 | 0.00 | 0.00 | -0.01 | 0.00 | 0.00 | 0.00 | 0.00 |
| | | | | | | | | | Coverage | | | | | | | | | | |
| null | 0.40 | 0.51 | 0.59 | 0.61 | 0.64 | 0.94 | 0.57 | 0.37 | 0.14 | 0.11 | 0.37 | 0.42 | 0.47 | 0.55 | 0.68 | 0.45 | 0.52 | 0.52 | 0.50 |
| fixed | 0.95 | 0.95 | 0.92 | 0.95 | 0.93 | 0.94 | 0.94 | 0.94 | 0.95 | 0.93 | 0.95 | 0.95 | 0.95 | 0.94 | 0.92 | 0.95 | 0.94 | 0.94 | 0.94 |
| mix2 | 0.70 | 0.77 | 0.80 | 0.69 | 0.72 | 0.93 | 0.66 | 0.52 | 0.91 | 0.91 | 0.67 | 0.72 | 0.75 | 0.79 | 0.84 | 0.73 | 0.78 | 0.76 | 0.75 |
| modMix | 0.76 | 0.81 | 0.82 | 0.74 | 0.77 | 0.95 | 0.68 | 0.63 | 0.92 | 0.91 | 0.73 | 0.77 | 0.79 | 0.82 | 0.86 | 0.79 | 0.81 | 0.79 | 0.79 |
| modFix | 0.86 | 0.86 | 0.83 | 0.81 | 0.84 | 0.95 | 0.72 | 0.81 | 0.93 | 0.93 | 0.78 | 0.82 | 0.85 | 0.89 | 0.92 | 0.87 | 0.85 | 0.83 | 0.85 |
| BW | 0.96 | 0.96 | 0.91 | 0.96 | 0.94 | 0.97 | 0.94 | 0.93 | 0.94 | 0.94 | 0.96 | 0.95 | 0.95 | 0.95 | 0.93 | 0.94 | 0.95 | 0.95 | 0.95 |
| modBW | 0.96 | 0.95 | 0.92 | 0.95 | 0.94 | 0.98 | 0.91 | 0.93 | 0.95 | 0.95 | 0.93 | 0.95 | 0.95 | 0.95 | 0.94 | 0.95 | 0.94 | 0.95 | 0.94 |
| TMB | 0.93 | 0.93 | 0.89 | 0.92 | 0.92 | 0.96 | 0.89 | 0.91 | 0.94 | 0.90 | 0.93 | 0.93 | 0.92 | 0.91 | 0.89 | 0.93 | 0.91 | 0.91 | 0.92 |
| modTMB | 0.90 | 0.90 | 0.87 | 0.88 | 0.89 | 0.95 | 0.83 | 0.87 | 0.92 | 0.92 | 0.90 | 0.89 | 0.89 | 0.89 | 0.89 | 0.91 | 0.89 | 0.87 | 0.89 |
| | | | | | | | | | SE | | | | | | | | | | |
| null | 0.01 | 0.03 | 0.04 | 0.02 | 0.03 | 0.03 | 0.03 | 0.03 | 0.03 | 0.04 | 0.01 | 0.02 | 0.02 | 0.03 | 0.05 | 0.02 | 0.03 | 0.03 | 0.03 |
| fixed | 0.06 | 0.08 | 0.13 | 0.08 | 0.10 | 0.09 | 0.09 | 0.09 | 0.10 | 0.09 | 0.09 | 0.09 | 0.09 | 0.09 | 0.09 | 0.07 | 0.10 | 0.11 | 0.09 |
| mix2 | 0.02 | 0.03 | 0.05 | 0.03 | 0.04 | 0.03 | 0.03 | 0.04 | 0.03 | 0.03 | 0.02 | 0.02 | 0.03 | 0.04 | 0.06 | 0.03 | 0.04 | 0.04 | 0.03 |
| modMix | 0.02 | 0.03 | 0.05 | 0.03 | 0.04 | 0.03 | 0.03 | 0.04 | 0.04 | 0.04 | 0.02 | 0.02 | 0.03 | 0.04 | 0.06 | 0.03 | 0.04 | 0.04 | 0.03 |
| modFix | 0.03 | 0.04 | 0.06 | 0.03 | 0.05 | 0.03 | 0.04 | 0.06 | 0.05 | 0.04 | 0.02 | 0.03 | 0.04 | 0.05 | 0.08 | 0.04 | 0.05 | 0.05 | 0.04 |
| BW | 0.05 | 0.07 | 0.12 | 0.06 | 0.09 | 0.07 | 0.07 | 0.08 | 0.08 | 0.08 | 0.07 | 0.07 | 0.08 | 0.08 | 0.09 | 0.06 | 0.08 | 0.09 | 0.08 |
| modBW | 0.04 | 0.06 | 0.08 | 0.05 | 0.07 | 0.05 | 0.06 | 0.07 | 0.06 | 0.05 | 0.04 | 0.04 | 0.05 | 0.06 | 0.08 | 0.04 | 0.06 | 0.07 | 0.06 |
| TMB | 0.04 | 0.05 | 0.07 | 0.04 | 0.06 | 0.03 | 0.05 | 0.06 | 0.08 | 0.08 | 0.05 | 0.05 | 0.05 | 0.06 | 0.07 | 0.04 | 0.06 | 0.06 | 0.06 |
| modTMB | 0.03 | 0.04 | 0.06 | 0.03 | 0.05 | 0.03 | 0.04 | 0.06 | 0.04 | 0.04 | 0.03 | 0.03 | 0.04 | 0.04 | 0.06 | 0.03 | 0.04 | 0.05 | 0.04 |

***Table S3.*** *Overall mean bias, 95% confidence interval coverage, and standard error (SE) for $\gamma_{m,2,1}$ by simulation design points for state-dependent distribution overlap, continuous or discrete random effects, number of individuals, and time series length from 360 scenarios without covariate effects.*

| Model | Overlap | | | Random effects: Continuous: State persistence | | Random effects: Continuous: $\sigma$ | | | Random effects: Discrete | | No. individuals ($M$) | | | | | Time series length ($T_m$) | | | Overall |
|---|---|---|---|---|---|---|---|---|---|---|---|---|---|---|---|---|---|---|---|
| | Little | Some | Much | Higher | Lower | 0 | 0.202 | 0.416 | mixA | mixB | 100 | 50 | 30 | 15 | 5 | 250 | 110 | 30 - 250 | |
| **Bias** | | | | | | | | | | | | | | | | | | | |
| null | -0.01 | -0.01 | 0.00 | 0.00 | 0.00 | 0.00 | 0.00 | -0.01 | -0.04 | 0.01 | -0.01 | -0.01 | -0.01 | 0.00 | 0.00 | -0.01 | 0.00 | 0.00 | 0.00 |
| fixed | 0.01 | 0.01 | 0.05 | 0.03 | 0.02 | 0.02 | 0.02 | 0.02 | 0.02 | 0.01 | 0.02 | 0.02 | 0.02 | 0.02 | 0.03 | 0.01 | 0.02 | 0.03 | 0.02 |
| mix2 | 0.00 | 0.00 | 0.01 | 0.00 | 0.01 | 0.01 | 0.01 | 0.00 | 0.00 | 0.00 | 0.00 | 0.00 | 0.00 | 0.01 | 0.02 | 0.00 | 0.01 | 0.01 | 0.00 |
| modMix | 0.00 | 0.00 | 0.01 | 0.00 | 0.01 | 0.00 | 0.00 | 0.00 | 0.00 | 0.00 | 0.00 | 0.00 | 0.00 | 0.00 | 0.01 | 0.00 | 0.00 | 0.00 | 0.00 |
| modFix | 0.00 | 0.00 | 0.01 | 0.00 | 0.01 | 0.01 | 0.00 | 0.01 | 0.01 | 0.00 | 0.00 | 0.00 | 0.00 | 0.01 | 0.02 | 0.00 | 0.01 | 0.01 | 0.01 |
| BW | 0.00 | -0.01 | 0.02 | 0.01 | -0.01 | 0.00 | 0.00 | 0.00 | 0.01 | 0.00 | 0.00 | 0.00 | 0.00 | 0.00 | 0.01 | 0.00 | 0.00 | 0.00 | 0.00 |
| modBW | 0.00 | 0.00 | 0.02 | 0.01 | 0.00 | 0.00 | 0.00 | 0.00 | 0.00 | 0.00 | 0.00 | 0.00 | 0.00 | 0.00 | 0.01 | 0.00 | 0.00 | 0.00 | 0.00 |
| TMB | 0.00 | 0.00 | 0.02 | 0.00 | 0.01 | 0.01 | 0.00 | 0.00 | 0.00 | 0.02 | 0.00 | 0.00 | 0.00 | 0.01 | 0.01 | 0.00 | 0.01 | 0.01 | 0.00 |
| modTMB | 0.00 | 0.00 | 0.01 | 0.00 | 0.01 | 0.01 | 0.00 | 0.00 | 0.00 | 0.00 | 0.00 | 0.00 | 0.00 | 0.00 | 0.01 | 0.00 | 0.00 | 0.00 | 0.00 |
| **Coverage** | | | | | | | | | | | | | | | | | | | |
| null | 0.40 | 0.50 | 0.64 | 0.61 | 0.64 | 0.94 | 0.57 | 0.37 | 0.24 | 0.11 | 0.39 | 0.44 | 0.49 | 0.56 | 0.69 | 0.47 | 0.54 | 0.54 | 0.51 |
| fixed | 0.95 | 0.95 | 0.92 | 0.94 | 0.94 | 0.94 | 0.95 | 0.94 | 0.94 | 0.94 | 0.95 | 0.95 | 0.95 | 0.94 | 0.93 | 0.95 | 0.95 | 0.94 | 0.94 |
| mix2 | 0.70 | 0.76 | 0.80 | 0.69 | 0.72 | 0.93 | 0.66 | 0.52 | 0.91 | 0.91 | 0.67 | 0.72 | 0.75 | 0.79 | 0.84 | 0.73 | 0.78 | 0.76 | 0.75 |
| modMix | 0.76 | 0.80 | 0.83 | 0.74 | 0.77 | 0.95 | 0.68 | 0.63 | 0.92 | 0.91 | 0.73 | 0.77 | 0.79 | 0.82 | 0.86 | 0.79 | 0.81 | 0.79 | 0.79 |
| modFix | 0.86 | 0.85 | 0.84 | 0.81 | 0.84 | 0.95 | 0.72 | 0.81 | 0.93 | 0.93 | 0.78 | 0.82 | 0.85 | 0.88 | 0.92 | 0.88 | 0.85 | 0.83 | 0.85 |
| BW | 0.96 | 0.95 | 0.94 | 0.96 | 0.95 | 0.98 | 0.95 | 0.94 | 0.94 | 0.94 | 0.96 | 0.96 | 0.95 | 0.95 | 0.94 | 0.95 | 0.95 | 0.95 | 0.95 |
| modBW | 0.96 | 0.95 | 0.92 | 0.95 | 0.94 | 0.98 | 0.92 | 0.93 | 0.95 | 0.95 | 0.93 | 0.95 | 0.95 | 0.95 | 0.94 | 0.95 | 0.94 | 0.95 | 0.95 |
| TMB | 0.92 | 0.92 | 0.90 | 0.92 | 0.92 | 0.96 | 0.89 | 0.91 | 0.91 | 0.90 | 0.93 | 0.93 | 0.92 | 0.91 | 0.89 | 0.93 | 0.91 | 0.91 | 0.92 |
| modTMB | 0.90 | 0.90 | 0.88 | 0.88 | 0.89 | 0.95 | 0.84 | 0.87 | 0.92 | 0.92 | 0.90 | 0.90 | 0.89 | 0.89 | 0.89 | 0.91 | 0.89 | 0.88 | 0.89 |
| **SE** | | | | | | | | | | | | | | | | | | | |
| null | 0.01 | 0.02 | 0.05 | 0.02 | 0.03 | 0.03 | 0.03 | 0.03 | 0.03 | 0.04 | 0.01 | 0.02 | 0.02 | 0.03 | 0.06 | 0.02 | 0.03 | 0.03 | 0.03 |
| fixed | 0.06 | 0.08 | 0.14 | 0.09 | 0.10 | 0.09 | 0.09 | 0.09 | 0.09 | 0.09 | 0.09 | 0.09 | 0.09 | 0.09 | 0.09 | 0.07 | 0.10 | 0.11 | 0.09 |
| mix2 | 0.02 | 0.03 | 0.06 | 0.03 | 0.04 | 0.03 | 0.03 | 0.04 | 0.03 | 0.03 | 0.02 | 0.02 | 0.03 | 0.04 | 0.06 | 0.03 | 0.04 | 0.04 | 0.03 |
| modMix | 0.02 | 0.03 | 0.06 | 0.03 | 0.04 | 0.03 | 0.03 | 0.04 | 0.04 | 0.03 | 0.02 | 0.02 | 0.03 | 0.04 | 0.06 | 0.03 | 0.04 | 0.04 | 0.04 |
| modFix | 0.03 | 0.04 | 0.06 | 0.04 | 0.05 | 0.03 | 0.04 | 0.06 | 0.05 | 0.04 | 0.02 | 0.03 | 0.04 | 0.05 | 0.08 | 0.04 | 0.05 | 0.05 | 0.04 |
| BW | 0.05 | 0.07 | 0.12 | 0.07 | 0.08 | 0.07 | 0.07 | 0.08 | 0.08 | 0.08 | 0.07 | 0.07 | 0.07 | 0.08 | 0.09 | 0.06 | 0.09 | 0.09 | 0.08 |
| modBW | 0.04 | 0.06 | 0.09 | 0.06 | 0.07 | 0.05 | 0.06 | 0.07 | 0.05 | 0.05 | 0.04 | 0.04 | 0.05 | 0.06 | 0.08 | 0.04 | 0.06 | 0.07 | 0.06 |
| TMB | 0.04 | 0.05 | 0.08 | 0.04 | 0.06 | 0.03 | 0.05 | 0.06 | 0.07 | 0.08 | 0.05 | 0.05 | 0.05 | 0.06 | 0.07 | 0.04 | 0.06 | 0.06 | 0.06 |
| modTMB | 0.03 | 0.04 | 0.06 | 0.04 | 0.05 | 0.03 | 0.04 | 0.06 | 0.04 | 0.03 | 0.03 | 0.03 | 0.04 | 0.05 | 0.07 | 0.03 | 0.05 | 0.05 | 0.04 |

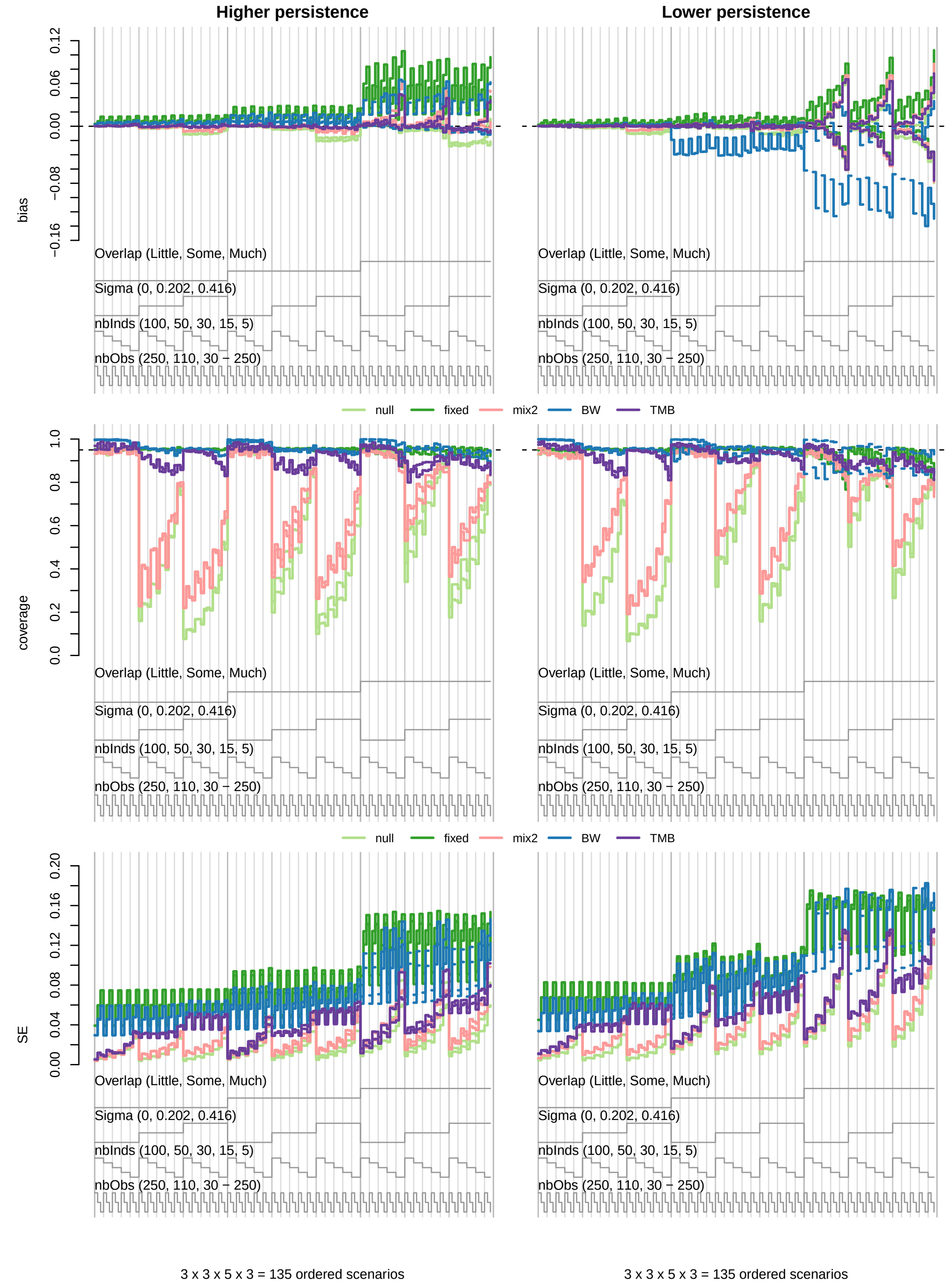

***Figure S4.*** *Nested loop plots for mean bias (top row), 95% confidence interval coverage (middle row), and standard error (SE; bottom row) for $\gamma_{m,1,2}$ and $\gamma_{m,2,1}$ from 135 simulated scenarios without covariates.*

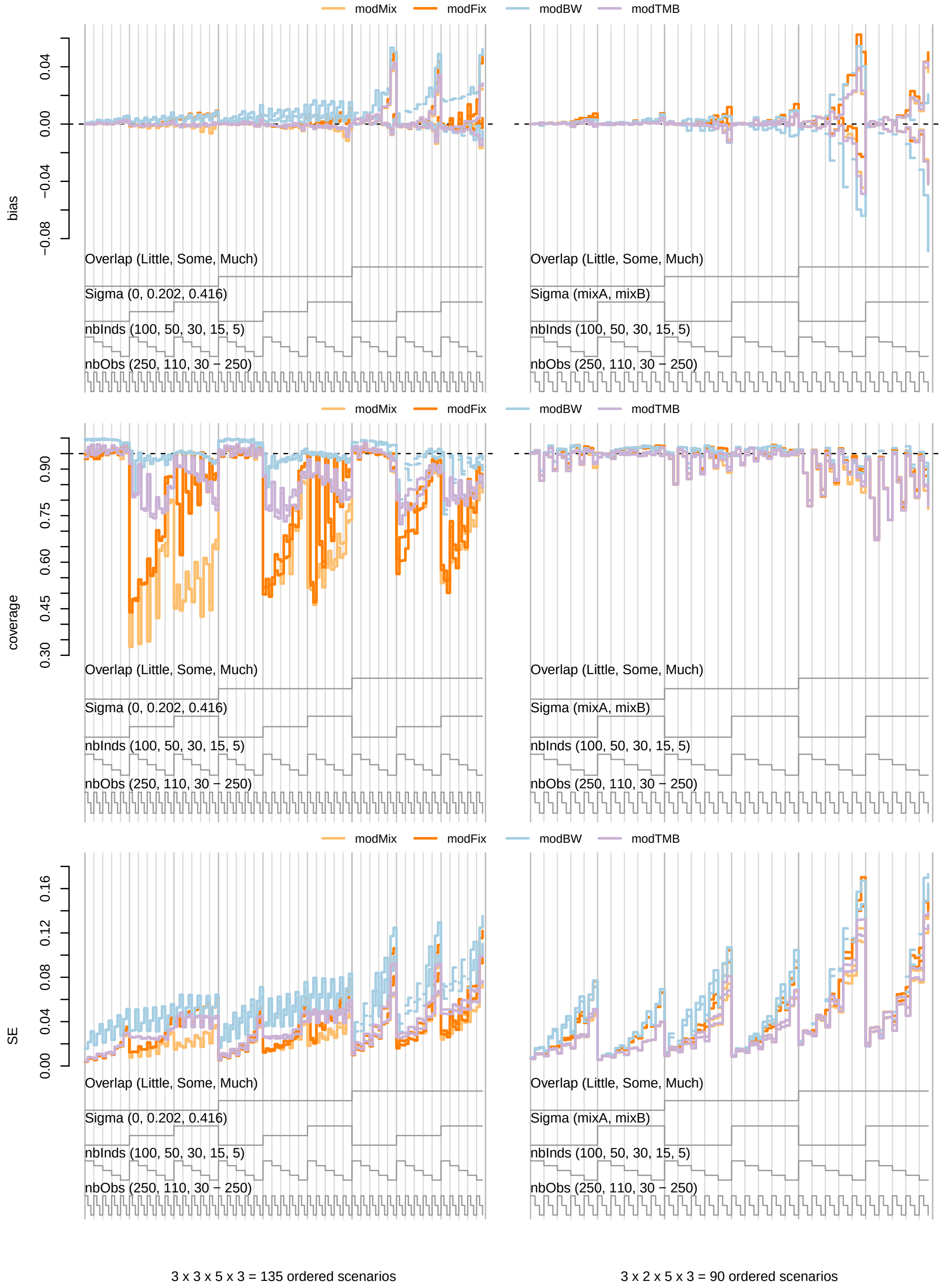

***Figure S5.*** *Nested loop plots for mean bias (top row), 95% confidence interval coverage (middle row), and standard error (SE; bottom row) for* $\gamma_{m,1,2}$ *and* $\gamma_{m,2,1}$ *from simulated scenarios without covariates, including 135 scenarios with "higher" state persistence (left column) and 90 scenarios with "mixA" or "mixB" finite mixtures (right column). Comparisons are for the* $AIC_c$ *model-averaged modMix (light orange), modFix (dark orange), modBW (light blue), and modTMB (light purple) models.*

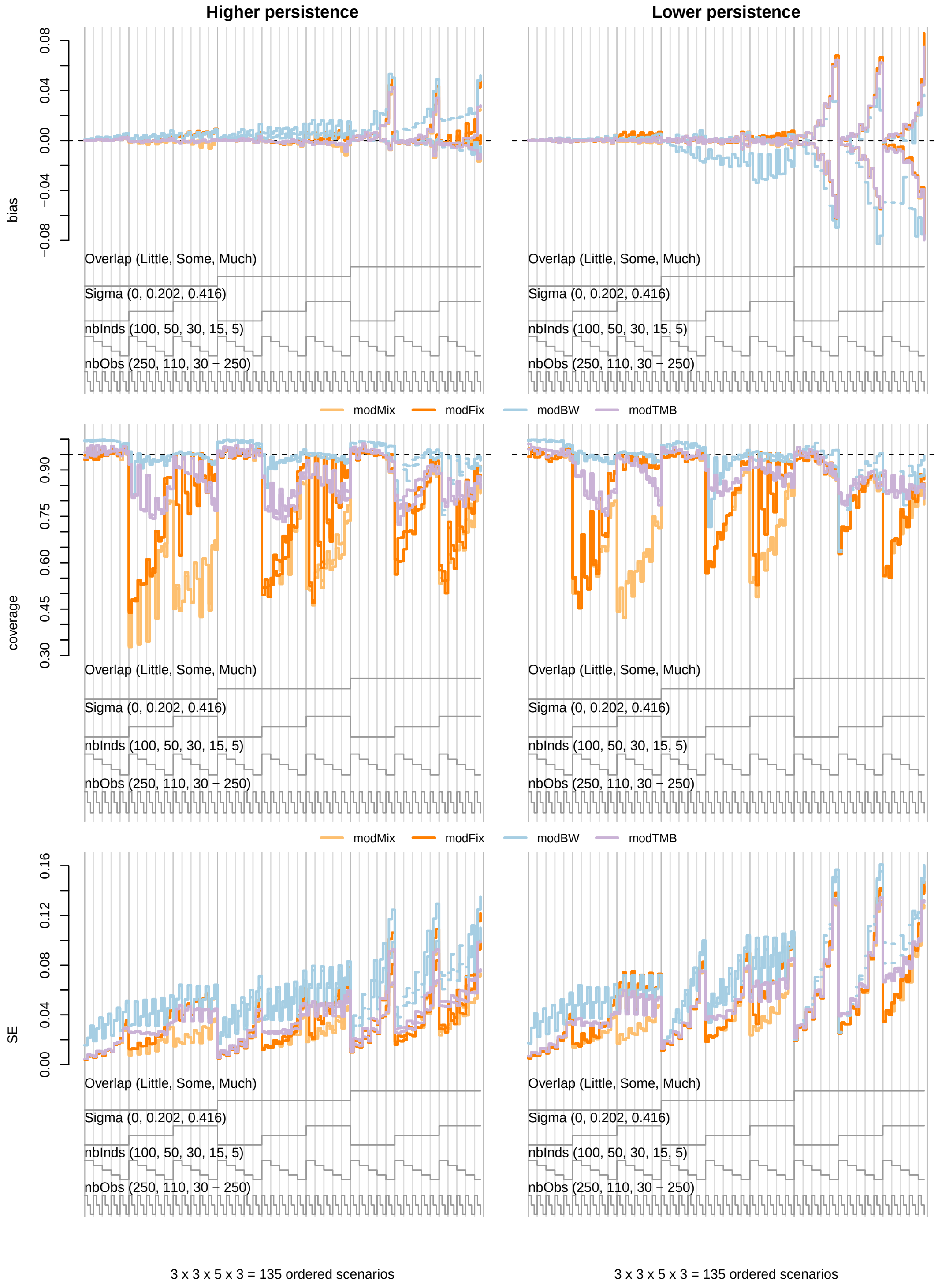

***Figure S6.*** *Nested loop plots for mean bias (top row), 95% confidence interval coverage (middle row), and standard error (SE; bottom row) for* $\gamma_{m,1,2}$ *and* $\gamma_{m,2,1}$ *from 135 simulated scenarios without covariates.*

***Table S4.*** *Overall mean percent relative bias, 95% confidence interval coverage, and percent standard error (SE) for $\mu_1^y$ by simulation design points for state-dependent distribution overlap, continuous or discrete random effects, number of individuals, and time series length from 360 scenarios without covariate effects.*

| Model | Overlap | | | Random effects: Continuous: State persistence | | Random effects: Continuous: $\boldsymbol{\sigma}$ | | | Random effects: Discrete | | No. individuals ($M$) | | | | | Time series length ($T_m$) | | | Overall |
|---|---|---|---|---|---|---|---|---|---|---|---|---|---|---|---|---|---|---|---|
| | Little | Some | Much | Higher | Lower | 0 | 0.202 | 0.416 | mixA | mixB | 100 | 50 | 30 | 15 | 5 | 250 | 110 | 30 - 250 | |
| **Bias** | | | | | | | | | | | | | | | | | | | |
| null | 0 | 0 | 0 | 0 | 0 | 0 | 0 | 0 | 1 | 1 | 0 | 0 | 0 | 0 | 1 | 0 | 0 | 0 | 0 |
| fixed | 0 | 0 | 0 | 0 | 0 | 0 | 0 | 0 | 0 | 0 | 0 | 0 | 0 | 0 | 0 | 0 | 0 | 0 | 0 |
| mix2 | 0 | 0 | 0 | 0 | 0 | 0 | 0 | 0 | 0 | 0 | 0 | 0 | 0 | 0 | 0 | 0 | 0 | 0 | 0 |
| modMix | 0 | 0 | 0 | 0 | 0 | 0 | 0 | 0 | 0 | 0 | 0 | 0 | 0 | 0 | 0 | 0 | 0 | 0 | 0 |
| modFix | 0 | 0 | 0 | 0 | 0 | 0 | 0 | 0 | 0 | 0 | 0 | 0 | 0 | 0 | 0 | 0 | 0 | 0 | 0 |
| modBW | 0 | 0 | 0 | 0 | 0 | 0 | 0 | 0 | 0 | 0 | 0 | 0 | 0 | 0 | 0 | 0 | 0 | 0 | 0 |
| TMB | 0 | 0 | 0 | 0 | 0 | 0 | 0 | 0 | 1 | 0 | 0 | 0 | 0 | 0 | 0 | 0 | 0 | 0 | 0 |
| modTMB | 0 | 0 | 0 | 0 | 0 | 0 | 0 | 0 | 0 | 0 | 0 | 0 | 0 | 0 | 0 | 0 | 0 | 0 | 0 |
| **Coverage** | | | | | | | | | | | | | | | | | | | |
| null | 0.95 | 0.92 | 0.94 | 0.94 | 0.94 | 0.94 | 0.94 | 0.94 | 0.93 | 0.93 | 0.94 | 0.94 | 0.94 | 0.94 | 0.92 | 0.94 | 0.94 | 0.94 | 0.94 |
| fixed | 0.95 | 0.92 | 0.91 | 0.93 | 0.91 | 0.92 | 0.92 | 0.92 | 0.93 | 0.93 | 0.93 | 0.93 | 0.93 | 0.93 | 0.91 | 0.94 | 0.92 | 0.92 | 0.93 |
| mix2 | 0.95 | 0.92 | 0.94 | 0.94 | 0.94 | 0.94 | 0.94 | 0.93 | 0.92 | 0.92 | 0.94 | 0.94 | 0.94 | 0.94 | 0.92 | 0.94 | 0.93 | 0.93 | 0.93 |
| modMix | 0.95 | 0.94 | 0.94 | 0.94 | 0.94 | 0.94 | 0.94 | 0.94 | 0.94 | 0.94 | 0.95 | 0.95 | 0.95 | 0.94 | 0.93 | 0.95 | 0.94 | 0.94 | 0.94 |
| modFix | 0.95 | 0.94 | 0.94 | 0.94 | 0.94 | 0.94 | 0.94 | 0.94 | 0.94 | 0.94 | 0.95 | 0.95 | 0.95 | 0.94 | 0.92 | 0.95 | 0.94 | 0.94 | 0.94 |
| modBW | 0.95 | 0.94 | 0.94 | 0.94 | 0.94 | 0.94 | 0.94 | 0.94 | 0.94 | 0.95 | 0.94 | 0.95 | 0.95 | 0.95 | 0.93 | 0.95 | 0.94 | 0.94 | 0.94 |
| TMB | 0.95 | 0.93 | 0.94 | 0.95 | 0.94 | 0.94 | 0.94 | 0.94 | 0.93 | 0.93 | 0.94 | 0.94 | 0.95 | 0.94 | 0.92 | 0.94 | 0.93 | 0.94 | 0.94 |
| modTMB | 0.95 | 0.94 | 0.94 | 0.94 | 0.94 | 0.94 | 0.94 | 0.94 | 0.94 | 0.94 | 0.95 | 0.95 | 0.95 | 0.94 | 0.93 | 0.95 | 0.94 | 0.94 | 0.94 |
| **Percent SE** | | | | | | | | | | | | | | | | | | | |
| null | 3 | 3 | 2 | 2 | 3 | 3 | 3 | 3 | 3 | 3 | 1 | 2 | 2 | 3 | 6 | 2 | 3 | 3 | 3 |
| fixed | 3 | 3 | 2 | 2 | 3 | 3 | 3 | 3 | 3 | 2 | 1 | 2 | 2 | 3 | 5 | 2 | 3 | 3 | 3 |
| mix2 | 3 | 3 | 2 | 2 | 3 | 3 | 3 | 3 | 3 | 2 | 1 | 2 | 2 | 3 | 5 | 2 | 3 | 3 | 3 |
| modMix | 3 | 3 | 2 | 2 | 3 | 3 | 3 | 3 | 3 | 2 | 1 | 2 | 2 | 3 | 5 | 2 | 3 | 3 | 3 |
| modFix | 3 | 3 | 2 | 2 | 3 | 3 | 3 | 3 | 3 | 2 | 1 | 2 | 2 | 3 | 5 | 2 | 3 | 3 | 3 |
| modBW | 3 | 3 | 3 | 2 | 3 | 3 | 3 | 3 | 3 | 3 | 1 | 2 | 2 | 3 | 5 | 2 | 3 | 3 | 3 |
| TMB | 3 | 3 | 2 | 2 | 3 | 3 | 3 | 3 | 3 | 2 | 1 | 2 | 2 | 3 | 5 | 2 | 3 | 3 | 3 |
| modTMB | 3 | 3 | 2 | 2 | 3 | 3 | 3 | 3 | 3 | 2 | 1 | 2 | 2 | 3 | 5 | 2 | 3 | 3 | 3 |

***Table S5.*** *Overall mean percent relative bias, 95% confidence interval coverage, and percent standard error (SE) for $\mu_2^y$ by simulation design points for state-dependent distribution overlap, continuous or discrete random effects, number of individuals, and time series length from 360 scenarios without covariate effects.*

| Model | Overlap | | | Random effects | | | | | | | No. individuals ($M$) | | | | | Time series length ($T_m$) | | | Overall |
|---|---|---|---|---|---|---|---|---|---|---|---|---|---|---|---|---|---|---|---|
| | | | | Continuous | | | | | Discrete | | | | | | | | | | |
| | | | | State persistence | | $\boldsymbol{\sigma}$ | | | | | | | | | | | | | |
| | Little | Some | Much | Higher | Lower | 0 | 0.202 | 0.416 | mixA | mixB | 100 | 50 | 30 | 15 | 5 | 250 | 110 | 30 - 250 | |
| | | | | | | | | | Bias | | | | | | | | | | |
| null | 0 | 0 | 2 | 0 | 1 | 1 | 1 | 1 | 0 | 0 | 0 | 0 | 0 | 1 | 2 | 0 | 1 | 1 | 1 |
| fixed | 0 | 0 | 3 | 1 | 1 | 1 | 1 | 1 | 1 | 1 | 0 | 0 | 1 | 1 | 2 | 0 | 1 | 1 | 1 |
| mix2 | 0 | 0 | 2 | 0 | 1 | 1 | 1 | 1 | 1 | 0 | 0 | 0 | 0 | 1 | 2 | 0 | 1 | 1 | 1 |
| modMix | 0 | 0 | 2 | 0 | 1 | 1 | 1 | 1 | 1 | 0 | 0 | 0 | 0 | 1 | 2 | 0 | 1 | 1 | 1 |
| modFix | 0 | 0 | 2 | 0 | 1 | 1 | 1 | 1 | 1 | 0 | 0 | 0 | 0 | 1 | 2 | 0 | 1 | 1 | 1 |
| modBW | 0 | 0 | 3 | 0 | 1 | 1 | 1 | 1 | 1 | 0 | 0 | 0 | 0 | 1 | 2 | 0 | 1 | 1 | 1 |
| TMB | 0 | 0 | 2 | 0 | 1 | 1 | 1 | 1 | 1 | 1 | 0 | 0 | 0 | 1 | 2 | 0 | 1 | 1 | 1 |
| modTMB | 0 | 0 | 2 | 0 | 1 | 1 | 1 | 1 | 1 | 0 | 0 | 0 | 0 | 1 | 2 | 0 | 1 | 1 | 1 |
| | | | | | | | | | Coverage | | | | | | | | | | |
| null | 0.95 | 0.94 | 0.93 | 0.95 | 0.94 | 0.94 | 0.94 | 0.94 | 0.94 | 0.93 | 0.95 | 0.95 | 0.95 | 0.94 | 0.92 | 0.94 | 0.94 | 0.94 | 0.94 |
| fixed | 0.95 | 0.93 | 0.89 | 0.93 | 0.91 | 0.92 | 0.92 | 0.93 | 0.93 | 0.92 | 0.93 | 0.93 | 0.93 | 0.92 | 0.91 | 0.94 | 0.92 | 0.92 | 0.93 |
| mix2 | 0.95 | 0.94 | 0.94 | 0.95 | 0.94 | 0.94 | 0.94 | 0.94 | 0.94 | 0.94 | 0.95 | 0.95 | 0.95 | 0.94 | 0.92 | 0.94 | 0.94 | 0.94 | 0.94 |
| modMix | 0.95 | 0.95 | 0.93 | 0.95 | 0.94 | 0.94 | 0.94 | 0.94 | 0.94 | 0.95 | 0.95 | 0.95 | 0.95 | 0.94 | 0.92 | 0.95 | 0.94 | 0.94 | 0.94 |
| modFix | 0.95 | 0.94 | 0.93 | 0.95 | 0.94 | 0.94 | 0.94 | 0.94 | 0.94 | 0.95 | 0.95 | 0.95 | 0.95 | 0.94 | 0.92 | 0.95 | 0.94 | 0.94 | 0.94 |
| modBW | 0.95 | 0.94 | 0.94 | 0.94 | 0.94 | 0.95 | 0.94 | 0.94 | 0.94 | 0.95 | 0.95 | 0.95 | 0.95 | 0.94 | 0.93 | 0.95 | 0.94 | 0.94 | 0.94 |
| TMB | 0.95 | 0.94 | 0.93 | 0.95 | 0.94 | 0.94 | 0.94 | 0.94 | 0.93 | 0.93 | 0.95 | 0.95 | 0.95 | 0.94 | 0.92 | 0.94 | 0.94 | 0.94 | 0.94 |
| modTMB | 0.95 | 0.95 | 0.94 | 0.95 | 0.94 | 0.94 | 0.94 | 0.94 | 0.95 | 0.95 | 0.95 | 0.95 | 0.95 | 0.94 | 0.92 | 0.95 | 0.94 | 0.94 | 0.94 |
| | | | | | | | | | Percent SE | | | | | | | | | | |
| null | 1 | 1 | 5 | 2 | 2 | 2 | 2 | 2 | 2 | 2 | 1 | 1 | 2 | 2 | 4 | 1 | 2 | 2 | 2 |
| fixed | 1 | 1 | 4 | 2 | 2 | 2 | 2 | 2 | 2 | 2 | 1 | 1 | 1 | 2 | 4 | 1 | 2 | 2 | 2 |
| mix2 | 1 | 1 | 4 | 2 | 2 | 2 | 2 | 2 | 2 | 2 | 1 | 1 | 1 | 2 | 4 | 1 | 2 | 2 | 2 |
| modMix | 1 | 1 | 4 | 2 | 2 | 2 | 2 | 2 | 2 | 2 | 1 | 1 | 1 | 2 | 4 | 1 | 2 | 2 | 2 |
| modFix | 1 | 1 | 4 | 2 | 2 | 2 | 2 | 2 | 2 | 2 | 1 | 1 | 1 | 2 | 4 | 1 | 2 | 2 | 2 |
| modBW | 1 | 1 | 5 | 2 | 2 | 2 | 2 | 2 | 2 | 2 | 1 | 1 | 1 | 2 | 4 | 1 | 2 | 2 | 2 |
| TMB | 1 | 1 | 5 | 2 | 2 | 2 | 2 | 2 | 2 | 2 | 1 | 1 | 2 | 2 | 4 | 1 | 2 | 2 | 2 |
| modTMB | 1 | 1 | 5 | 2 | 2 | 2 | 2 | 2 | 2 | 2 | 1 | 1 | 2 | 2 | 4 | 1 | 2 | 2 | 2 |

***Table S6.*** *Overall mean percent relative bias, 95% confidence interval coverage, and percent standard error (SE) for $\sigma_1^y$ by simulation design points for state-dependent distribution overlap, continuous or discrete random effects, number of individuals, and time series length from 360 scenarios without covariate effects.*

| Model | Overlap | | | Random effects | | | | | | | No. individuals ($M$) | | | | | Time series length ($T_m$) | | | Overall |
|---|---|---|---|---|---|---|---|---|---|---|---|---|---|---|---|---|---|---|---|
| | | | | Continuous | | | | | Discrete | | | | | | | | | | |
| | | | | State persistence | | $\boldsymbol{\sigma}$ | | | | | | | | | | | | | |
| | Little | Some | Much | Higher | Lower | 0 | 0.202 | 0.416 | mixA | mixB | 100 | 50 | 30 | 15 | 5 | 250 | 110 | 30 - 250 | |
| | Bias | | | | | | | | | | | | | | | | | | |
| null | 0 | 1 | 0 | 0 | 0 | 0 | 0 | 0 | 1 | 1 | 0 | 0 | 0 | 1 | 1 | 0 | 0 | 0 | 0 |
| fixed | 0 | -1 | 1 | 0 | 0 | 0 | 0 | 0 | 0 | 0 | 0 | 0 | 0 | 0 | 0 | 0 | 0 | 0 | 0 |
| mix2 | 0 | 0 | 0 | 0 | 0 | 0 | 0 | 0 | 0 | 0 | 0 | 0 | 0 | 0 | 0 | 0 | 0 | 0 | 0 |
| modMix | 0 | 0 | 0 | 0 | 0 | 0 | 0 | 0 | 0 | 0 | 0 | 0 | 0 | 0 | 0 | 0 | 0 | 0 | 0 |
| modFix | 0 | 0 | 0 | 0 | 0 | 0 | 0 | 0 | 0 | 0 | 0 | 0 | 0 | 0 | 0 | 0 | 0 | 0 | 0 |
| modBW | 0 | -1 | 0 | 0 | 0 | 0 | 0 | 0 | 0 | 0 | 0 | 0 | 0 | 0 | 0 | 0 | 0 | 0 | 0 |
| TMB | 0 | 0 | 1 | 0 | 0 | 0 | 0 | 0 | 1 | 0 | 0 | 0 | 0 | 0 | 0 | 0 | 0 | 0 | 0 |
| modTMB | 0 | 0 | 0 | 0 | 0 | 0 | 0 | 0 | 0 | 0 | 0 | 0 | 0 | 0 | 0 | 0 | 0 | 0 | 0 |
| | Coverage | | | | | | | | | | | | | | | | | | |
| null | 0.95 | 0.92 | 0.93 | 0.94 | 0.93 | 0.94 | 0.94 | 0.93 | 0.93 | 0.92 | 0.94 | 0.94 | 0.94 | 0.93 | 0.91 | 0.94 | 0.93 | 0.93 | 0.93 |
| fixed | 0.95 | 0.91 | 0.89 | 0.92 | 0.90 | 0.91 | 0.92 | 0.92 | 0.92 | 0.92 | 0.92 | 0.93 | 0.92 | 0.91 | 0.90 | 0.93 | 0.91 | 0.91 | 0.92 |
| mix2 | 0.95 | 0.91 | 0.93 | 0.94 | 0.93 | 0.94 | 0.94 | 0.93 | 0.92 | 0.92 | 0.94 | 0.94 | 0.94 | 0.93 | 0.91 | 0.93 | 0.93 | 0.93 | 0.93 |
| modMix | 0.95 | 0.94 | 0.93 | 0.94 | 0.93 | 0.94 | 0.94 | 0.94 | 0.94 | 0.94 | 0.95 | 0.95 | 0.94 | 0.94 | 0.92 | 0.94 | 0.94 | 0.94 | 0.94 |
| modFix | 0.95 | 0.94 | 0.93 | 0.94 | 0.93 | 0.94 | 0.94 | 0.93 | 0.94 | 0.94 | 0.95 | 0.95 | 0.94 | 0.93 | 0.92 | 0.94 | 0.93 | 0.94 | 0.94 |
| modBW | 0.95 | 0.93 | 0.93 | 0.94 | 0.94 | 0.94 | 0.94 | 0.93 | 0.94 | 0.94 | 0.95 | 0.94 | 0.94 | 0.94 | 0.93 | 0.94 | 0.94 | 0.94 | 0.94 |
| TMB | 0.95 | 0.93 | 0.92 | 0.94 | 0.93 | 0.94 | 0.94 | 0.94 | 0.92 | 0.92 | 0.94 | 0.94 | 0.94 | 0.93 | 0.91 | 0.94 | 0.93 | 0.93 | 0.93 |
| modTMB | 0.95 | 0.94 | 0.93 | 0.94 | 0.94 | 0.94 | 0.94 | 0.94 | 0.94 | 0.94 | 0.95 | 0.95 | 0.94 | 0.94 | 0.92 | 0.94 | 0.94 | 0.94 | 0.94 |
| | Percent SE | | | | | | | | | | | | | | | | | | |
| null | 4 | 6 | 6 | 4 | 6 | 5 | 5 | 5 | 5 | 6 | 2 | 3 | 4 | 6 | 10 | 4 | 6 | 6 | 5 |
| fixed | 4 | 5 | 5 | 4 | 5 | 5 | 5 | 5 | 5 | 4 | 2 | 3 | 4 | 5 | 9 | 4 | 5 | 5 | 5 |
| mix2 | 4 | 6 | 5 | 4 | 6 | 5 | 5 | 5 | 5 | 4 | 2 | 3 | 4 | 6 | 9 | 4 | 5 | 5 | 5 |
| modMix | 4 | 6 | 6 | 4 | 6 | 5 | 5 | 5 | 5 | 4 | 2 | 3 | 4 | 6 | 10 | 4 | 6 | 6 | 5 |
| modFix | 4 | 6 | 6 | 4 | 6 | 5 | 5 | 5 | 5 | 4 | 2 | 3 | 4 | 6 | 10 | 4 | 6 | 6 | 5 |
| modBW | 4 | 6 | 6 | 4 | 6 | 5 | 5 | 5 | 5 | 4 | 2 | 3 | 4 | 6 | 10 | 4 | 6 | 6 | 5 |
| TMB | 4 | 6 | 6 | 4 | 6 | 5 | 5 | 5 | 5 | 4 | 2 | 3 | 4 | 6 | 10 | 4 | 6 | 6 | 5 |
| modTMB | 4 | 6 | 6 | 4 | 6 | 5 | 5 | 5 | 5 | 4 | 2 | 3 | 4 | 6 | 10 | 4 | 6 | 6 | 5 |

***Table S7.*** *Overall mean percent relative bias, 95% confidence interval coverage, and percent standard error (SE) for $\sigma_2^y$ by simulation design points for state-dependent distribution overlap, continuous or discrete random effects, number of individuals, and time series length from 360 scenarios without covariate effects.*

| Model | Overlap | | | Random effects: Continuous, State persistence | | Random effects: Continuous, $\boldsymbol{\sigma}$ | | | Random effects: Discrete | | No. individuals ($M$) | | | | | Time series length ($T_m$) | | | Overall |
|---|---|---|---|---|---|---|---|---|---|---|---|---|---|---|---|---|---|---|---|
| | Little | Some | Much | Higher | Lower | 0 | 0.202 | 0.416 | mixA | mixB | 100 | 50 | 30 | 15 | 5 | 250 | 110 | 30 - 250 | |
| **Bias** | | | | | | | | | | | | | | | | | | | |
| null | 0 | 0 | 0 | 0 | 0 | 0 | 0 | 0 | 0 | 0 | 0 | 0 | 0 | 0 | -1 | 0 | 0 | 0 | 0 |
| fixed | 0 | 0 | 0 | 0 | 0 | 0 | 0 | 0 | 0 | 0 | 0 | 0 | 0 | 0 | -1 | 0 | 0 | 0 | 0 |
| mix2 | 0 | 0 | 0 | 0 | 0 | 0 | 0 | 0 | 0 | 0 | 0 | 0 | 0 | 0 | -1 | 0 | 0 | 0 | 0 |
| modMix | 0 | 0 | 0 | 0 | 0 | 0 | 0 | 0 | 0 | 0 | 0 | 0 | 0 | 0 | -1 | 0 | 0 | 0 | 0 |
| modFix | 0 | 0 | 0 | 0 | 0 | 0 | 0 | 0 | 0 | 0 | 0 | 0 | 0 | 0 | -1 | 0 | 0 | 0 | 0 |
| modBW | 0 | 0 | 0 | 0 | 0 | 0 | 0 | 0 | 0 | 0 | 0 | 0 | 0 | 0 | -1 | 0 | 0 | 0 | 0 |
| TMB | 0 | 0 | 0 | 0 | 0 | 0 | 0 | 0 | 0 | 0 | 0 | 0 | 0 | 0 | -1 | 0 | 0 | 0 | 0 |
| modTMB | 0 | 0 | 0 | 0 | 0 | 0 | 0 | 0 | 0 | 0 | 0 | 0 | 0 | 0 | -1 | 0 | 0 | 0 | 0 |
| **Coverage** | | | | | | | | | | | | | | | | | | | |
| null | 0.95 | 0.95 | 0.94 | 0.95 | 0.95 | 0.95 | 0.95 | 0.95 | 0.95 | 0.94 | 0.95 | 0.95 | 0.95 | 0.95 | 0.94 | 0.95 | 0.94 | 0.95 | 0.95 |
| fixed | 0.95 | 0.94 | 0.93 | 0.94 | 0.94 | 0.94 | 0.94 | 0.94 | 0.94 | 0.95 | 0.94 | 0.94 | 0.94 | 0.94 | 0.93 | 0.95 | 0.94 | 0.94 | 0.94 |
| mix2 | 0.95 | 0.95 | 0.94 | 0.95 | 0.94 | 0.94 | 0.95 | 0.95 | 0.95 | 0.95 | 0.95 | 0.95 | 0.95 | 0.95 | 0.94 | 0.95 | 0.94 | 0.95 | 0.95 |
| modMix | 0.95 | 0.95 | 0.95 | 0.95 | 0.95 | 0.95 | 0.95 | 0.95 | 0.95 | 0.95 | 0.95 | 0.95 | 0.95 | 0.95 | 0.94 | 0.95 | 0.95 | 0.95 | 0.95 |
| modFix | 0.95 | 0.95 | 0.94 | 0.95 | 0.95 | 0.95 | 0.95 | 0.95 | 0.95 | 0.95 | 0.95 | 0.95 | 0.95 | 0.95 | 0.94 | 0.95 | 0.94 | 0.95 | 0.95 |
| modBW | 0.95 | 0.95 | 0.95 | 0.95 | 0.95 | 0.95 | 0.95 | 0.95 | 0.95 | 0.95 | 0.95 | 0.95 | 0.95 | 0.95 | 0.94 | 0.95 | 0.95 | 0.95 | 0.95 |
| TMB | 0.95 | 0.95 | 0.94 | 0.95 | 0.95 | 0.95 | 0.95 | 0.95 | 0.94 | 0.95 | 0.95 | 0.95 | 0.95 | 0.95 | 0.94 | 0.95 | 0.94 | 0.95 | 0.95 |
| modTMB | 0.95 | 0.95 | 0.95 | 0.95 | 0.95 | 0.95 | 0.95 | 0.95 | 0.95 | 0.95 | 0.95 | 0.95 | 0.95 | 0.95 | 0.94 | 0.95 | 0.95 | 0.95 | 0.95 |
| **Percent SE** | | | | | | | | | | | | | | | | | | | |
| null | 2 | 3 | 3 | 3 | 3 | 3 | 3 | 3 | 3 | 3 | 1 | 2 | 2 | 3 | 6 | 2 | 3 | 3 | 3 |
| fixed | 2 | 3 | 3 | 3 | 3 | 3 | 3 | 3 | 3 | 3 | 1 | 2 | 2 | 3 | 5 | 2 | 3 | 3 | 3 |
| mix2 | 2 | 3 | 3 | 3 | 3 | 3 | 3 | 3 | 3 | 3 | 1 | 2 | 2 | 3 | 5 | 2 | 3 | 3 | 3 |
| modMix | 2 | 3 | 3 | 3 | 3 | 3 | 3 | 3 | 3 | 3 | 1 | 2 | 2 | 3 | 6 | 2 | 3 | 3 | 3 |
| modFix | 2 | 3 | 3 | 3 | 3 | 3 | 3 | 3 | 3 | 3 | 1 | 2 | 2 | 3 | 6 | 2 | 3 | 3 | 3 |
| modBW | 2 | 3 | 4 | 3 | 3 | 3 | 3 | 3 | 3 | 3 | 1 | 2 | 2 | 3 | 5 | 2 | 3 | 3 | 3 |
| TMB | 2 | 3 | 3 | 3 | 3 | 3 | 3 | 3 | 3 | 3 | 1 | 2 | 2 | 3 | 6 | 2 | 3 | 3 | 3 |
| modTMB | 2 | 3 | 3 | 3 | 3 | 3 | 3 | 3 | 3 | 3 | 1 | 2 | 2 | 3 | 6 | 2 | 3 | 3 | 3 |

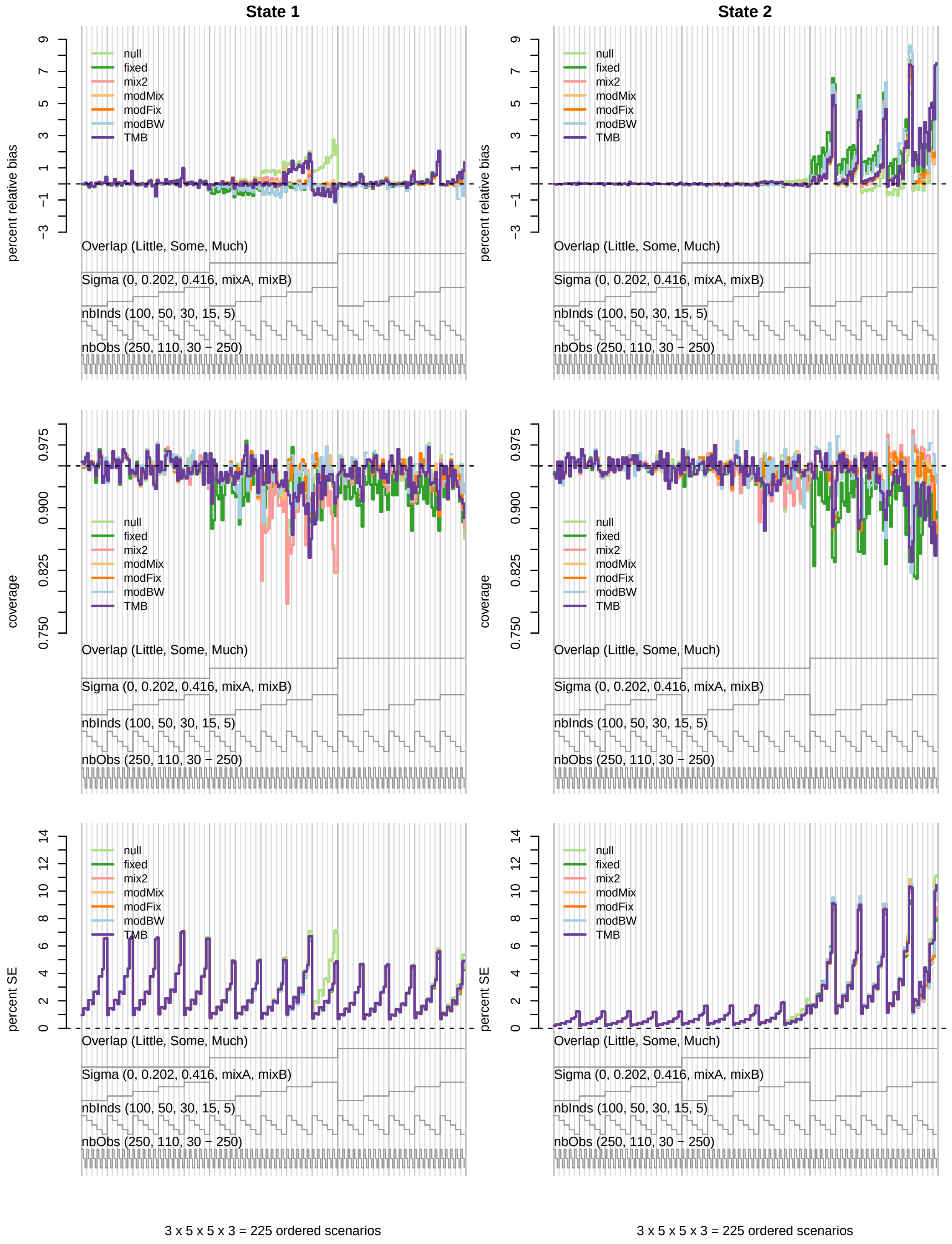

***Figure S7.*** *Nested loop plots for mean percent relative bias (top row), 95% confidence interval coverage (middle row), and percent standard error (SE; bottom row) for $\mu_1^y$ (left column) and $\mu_2^y$ (right column) from 225 simulated scenarios without covariate effects. Continuous random effect scenarios are limited to those with "higher" state persistence.*

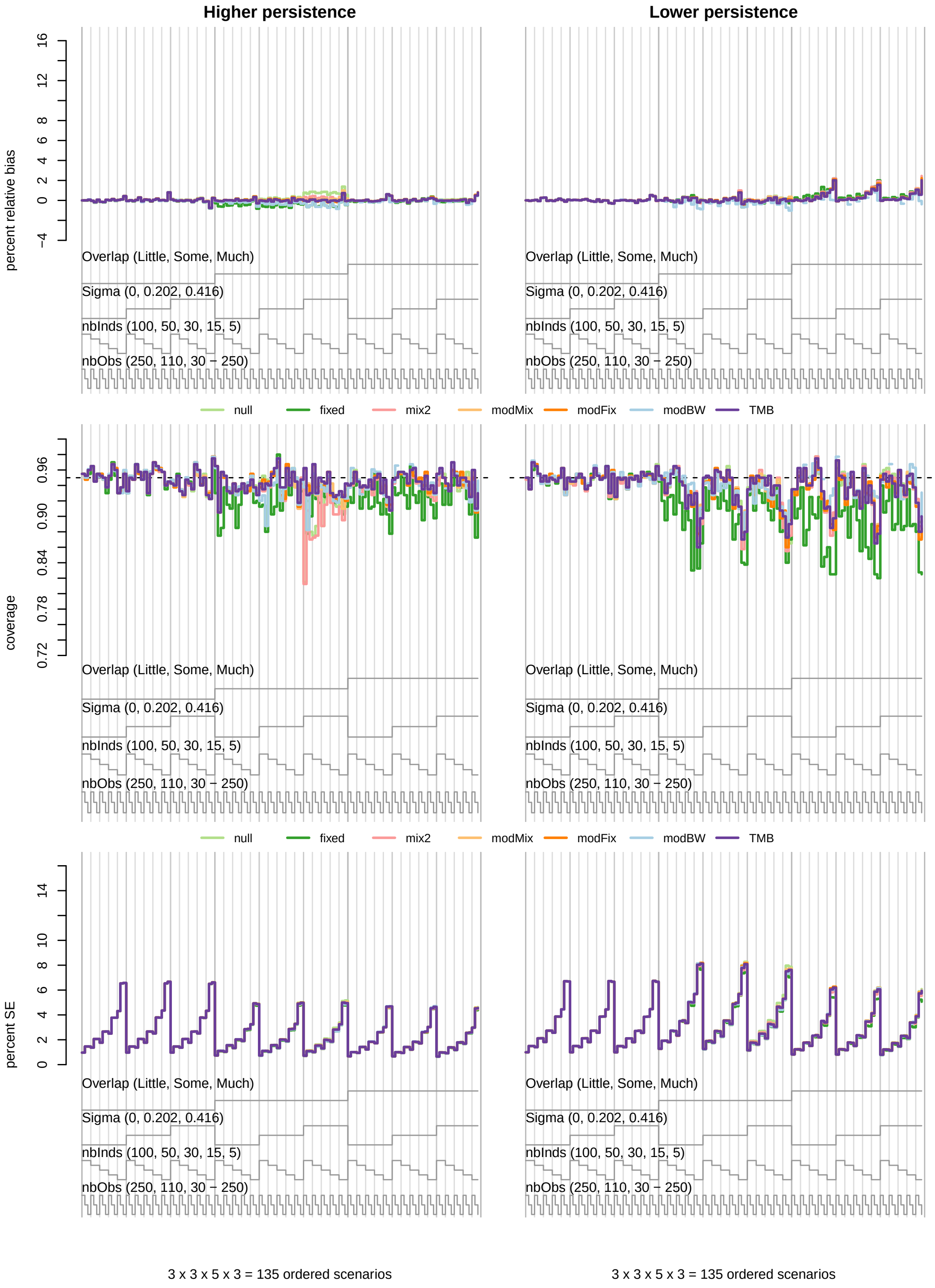

***Figure S8.*** *Nested loop plots for mean percent relative bias (top row), 95% confidence interval coverage (middle row), and percent standard error (SE; bottom row) for* $\mu_1^y$ *from 135 simulated scenarios without covariate effects.*

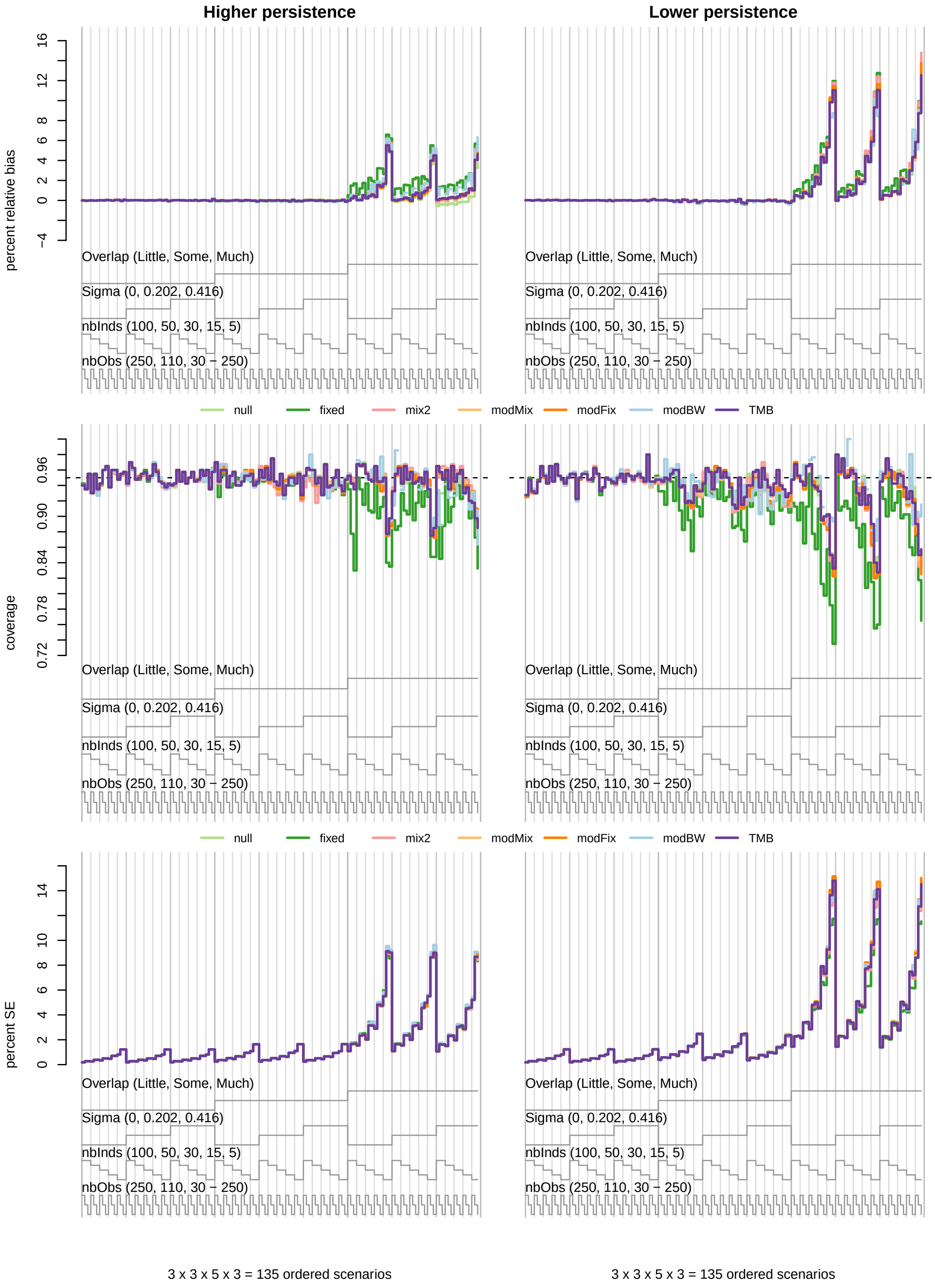

***Figure S9.*** *Nested loop plots for mean percent relative bias (top row), 95% confidence interval coverage (middle row), and percent standard error (SE; bottom row) for* $\mu_2^y$ *from 135 simulated without covariate effects.*

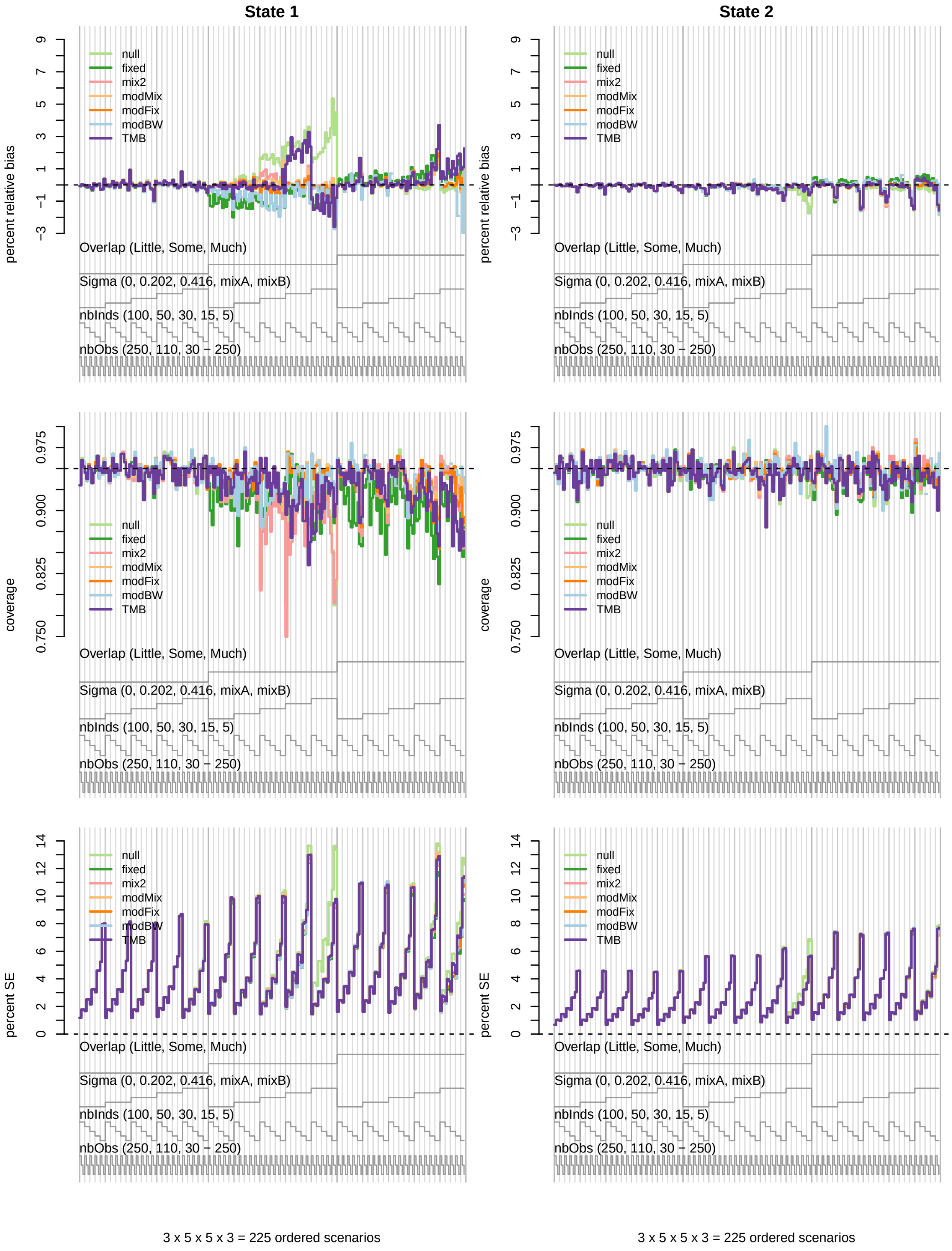

***Figure S10.*** *Nested loop plots for mean percent relative bias (top row), 95% confidence interval coverage (middle row), and percent standard error (SE; bottom row) for* $\sigma_1^y$ *(left column) and* $\sigma_2^y$ *(right column) from 225 simulated scenarios without covariate effects. Continuous random effect scenarios are limited to those with "higher" state persistence.*

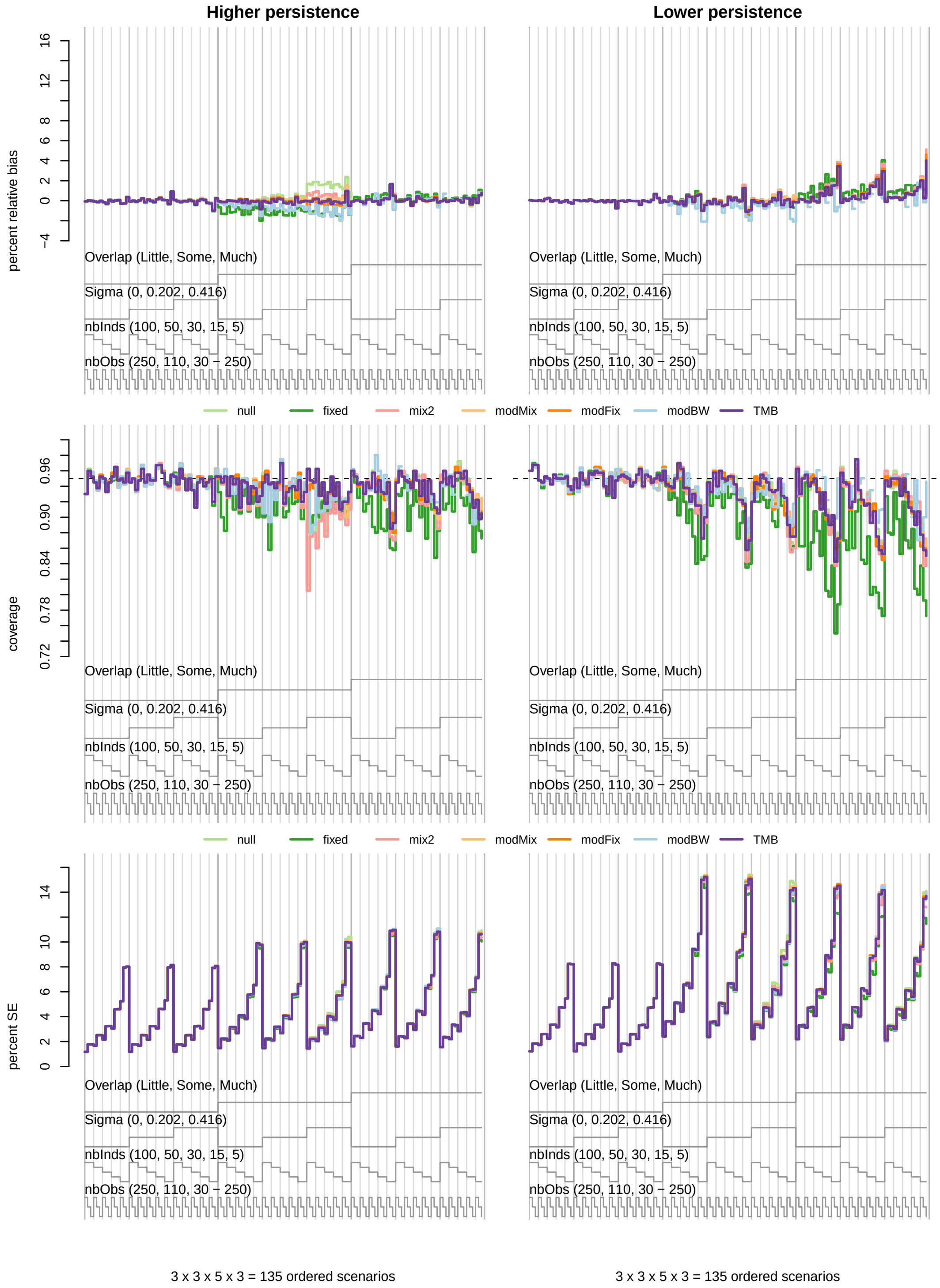

***Figure S11.*** *Nested loop plots for mean percent relative bias (top row), 95% confidence interval coverage (middle row), and percent standard error (SE; bottom row) for* $\sigma_1^y$ *from 135 simulated scenarios without covariate effects.*

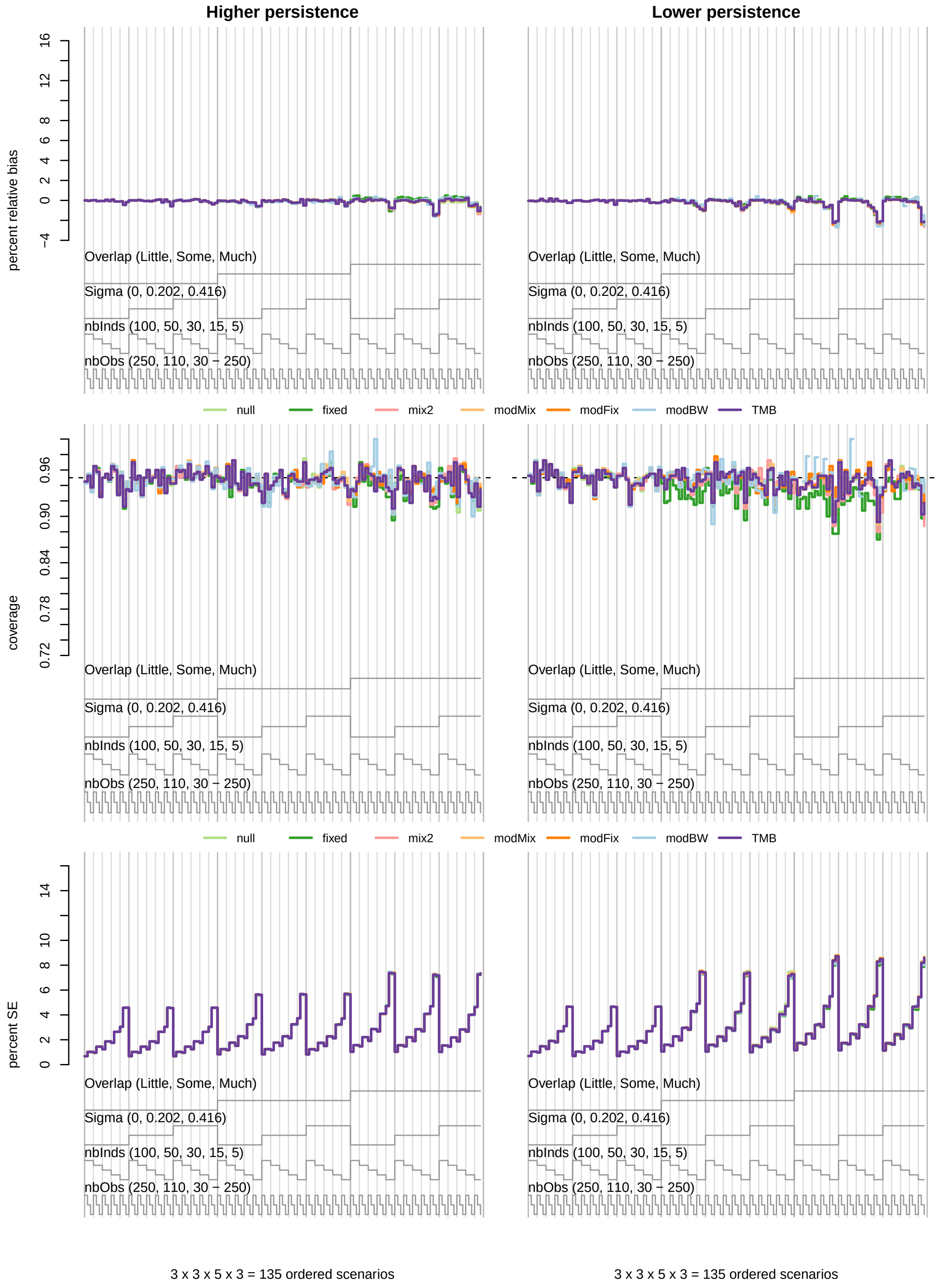

***Figure S12.*** *Nested loop plots for mean percent relative bias (top row), 95% confidence interval coverage (middle row), and percent standard error (SE; bottom row) for* $\sigma_2^y$ *from 135 simulated scenarios without covariate effects.*

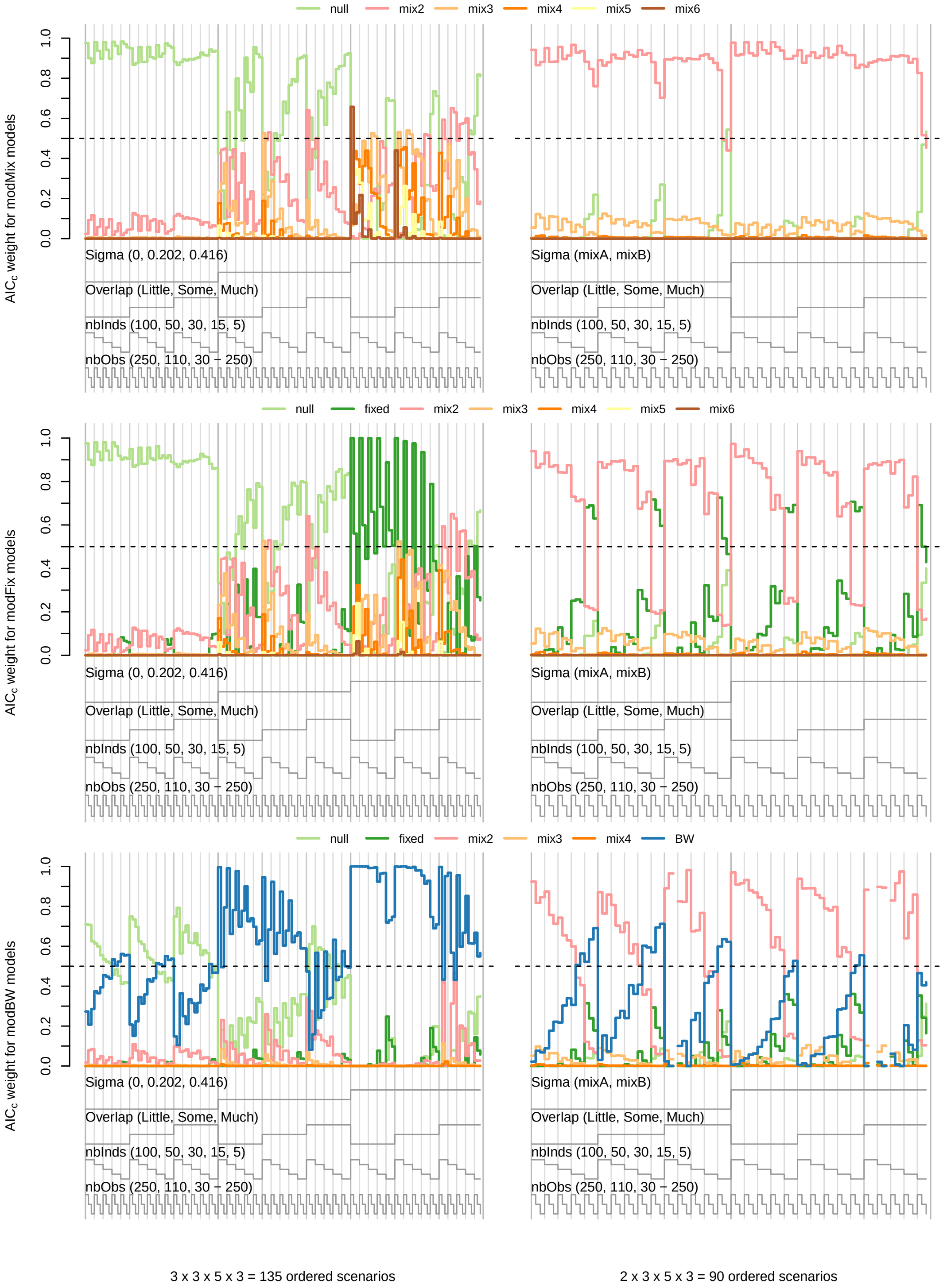

***Figure S13.*** *Nested loop plots for mean $AIC_c$ weights in candidate model sets from simulated scenarios without covariates, including 135 scenarios with "higher" state persistence (left column) and 90 scenarios with "mixA" or "mixB" finite mixtures (right column).*

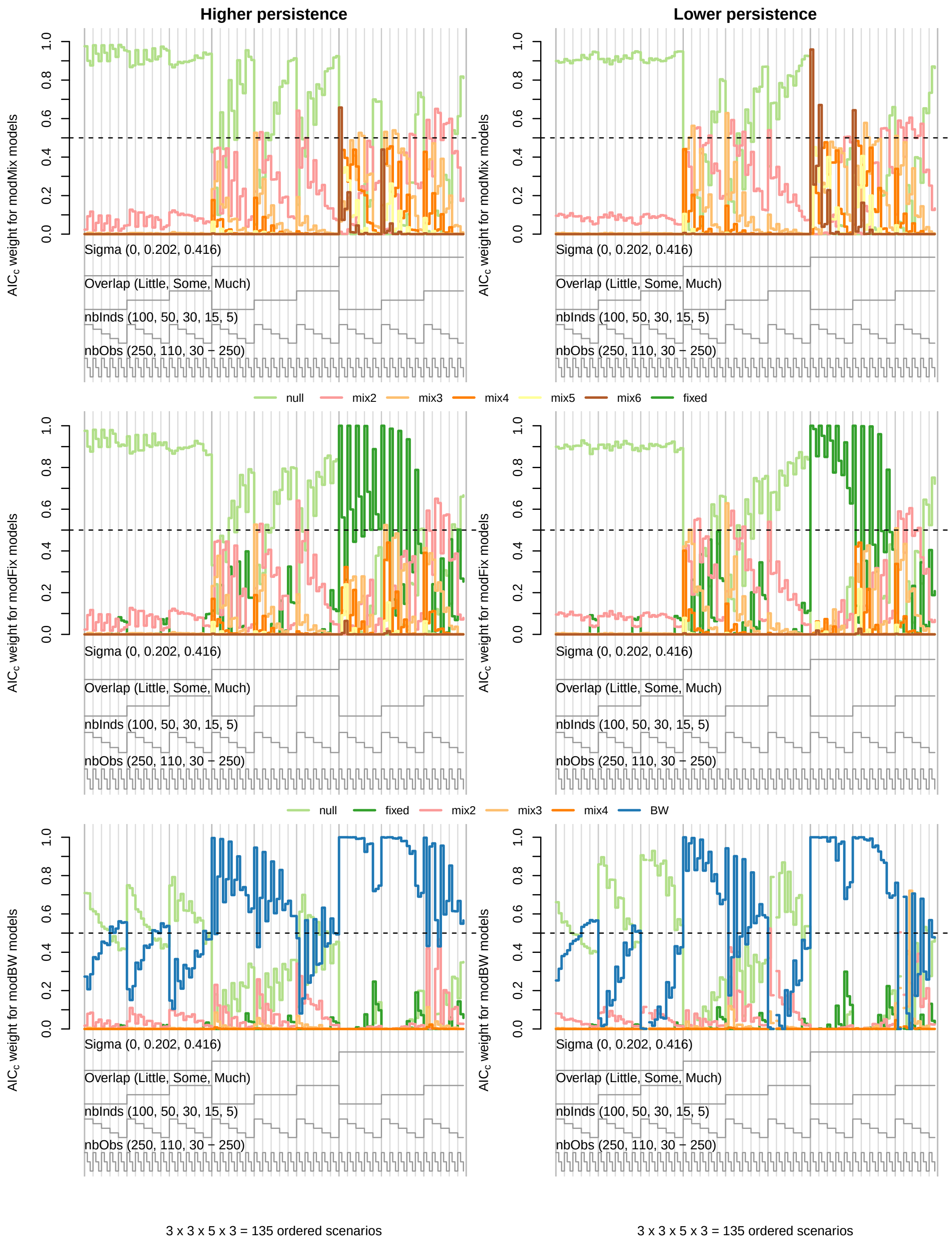

***Figure S14.*** *Nested loop plots for mean AIC$_c$ weights in candidate model sets from 135 simulated scenarios without covariates.*

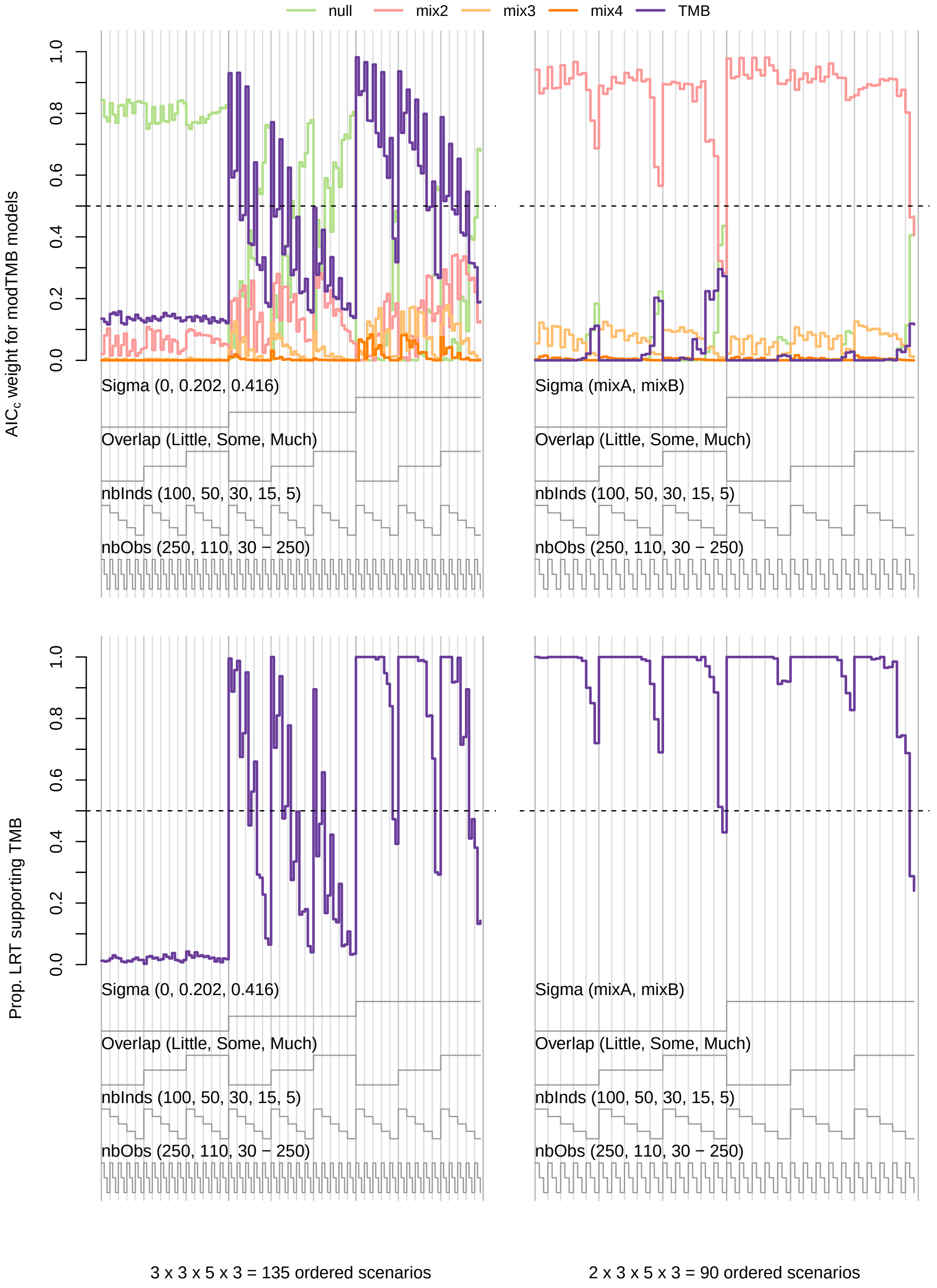

***Figure S15.*** *Nested loop plots for mean AIC$_c$ weights based on the "modTMB" candidate model set (top row) and proportion of likelihood ratio tests (LRT) supporting TMB over the null model (bottom row) from simulated scenarios without covariates, including 135 scenarios with "higher" state persistence (left column) and 90 scenarios with "mixA" or "mixB" finite mixtures (right column).*

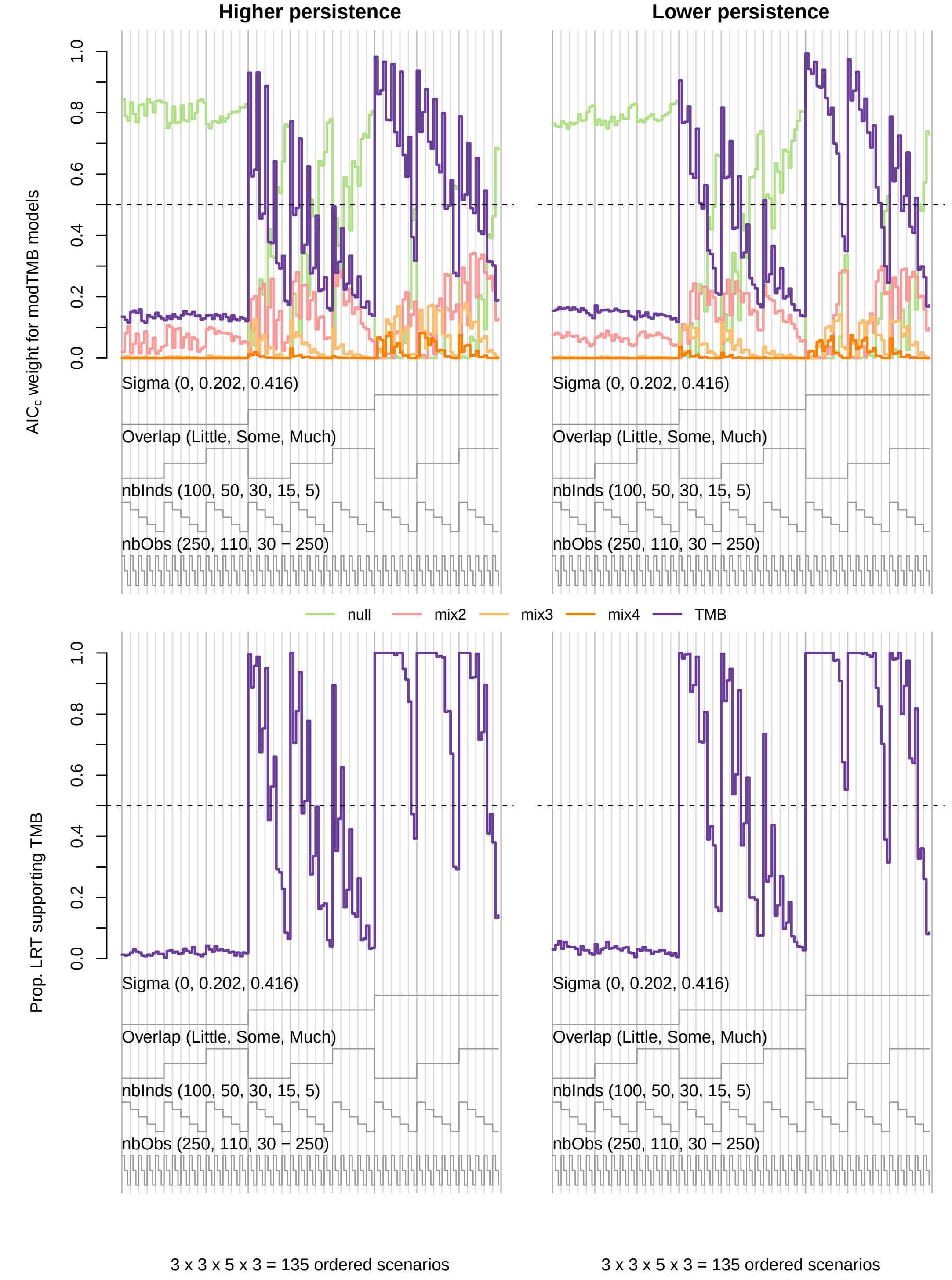

***Figure S16.*** *Nested loop plots for mean AIC$_c$ weights based on the "modTMB" candidate model set (top row) and proportion of likelihood ratio tests (LRT) supporting TMB over the null model (bottom row) from 135 simulated scenarios without covariates.*

##### S2.2.2 With covariates

***Table S8.*** *Overall mean proportion of Viterbi-decoded states that were correctly classified after accounting for chance agreement and the proportion of estimated state probabilities in which the true state received at least 0.50 or 0.20 probability by simulation design points for state-dependent distribution overlap, state persistence, covariate effects* ($\mu_{1,1,2}, \mu_{1,2,1}$)*, individual heterogeneity* ($\boldsymbol{\sigma}$)*, number of individuals, and time series length from 540 scenarios with covariate effects. All models (except "fixed") included terms for the covariate effects.*

| Model | Overlap | | | State persistence | | ($\mu_{1,1,2}, \mu_{1,2,1}$) | | $\boldsymbol{\sigma}$ | | | No. individuals ($M$) | | | | | Time series length ($T_m$) | | | Overall |
|---|---|---|---|---|---|---|---|---|---|---|---|---|---|---|---|---|---|---|---|
| | Little | Some | Much | Higher | Lower | (0, 0) | (0.5, −0.5) | 0 | 0.202 | 0.416 | 100 | 50 | 30 | 15 | 5 | 250 | 110 | 30 - 250 | |
| | Prop. states correctly classified | | | | | | | | | | | | | | | | | | |
| null | 0.99 | 0.82 | 0.52 | 0.80 | 0.76 | 0.77 | 0.78 | 0.78 | 0.78 | 0.78 | 0.78 | 0.78 | 0.78 | 0.78 | 0.76 | 0.78 | 0.78 | 0.78 | 0.78 |
| fixed | 0.99 | 0.82 | 0.50 | 0.80 | 0.75 | 0.76 | 0.78 | 0.77 | 0.77 | 0.77 | 0.78 | 0.77 | 0.77 | 0.77 | 0.76 | 0.78 | 0.77 | 0.77 | 0.77 |
| mix2 | 0.99 | 0.82 | 0.52 | 0.80 | 0.75 | 0.77 | 0.78 | 0.78 | 0.78 | 0.78 | 0.78 | 0.78 | 0.78 | 0.78 | 0.76 | 0.78 | 0.78 | 0.78 | 0.78 |
| mix3 | 0.99 | 0.82 | 0.52 | 0.80 | 0.75 | 0.77 | 0.78 | 0.78 | 0.78 | 0.78 | 0.78 | 0.78 | 0.78 | 0.78 | 0.76 | 0.78 | 0.78 | 0.77 | 0.78 |
| mix4 | 0.99 | 0.82 | 0.52 | 0.80 | 0.75 | 0.77 | 0.78 | 0.78 | 0.78 | 0.78 | 0.78 | 0.78 | 0.78 | 0.78 | 0.76 | 0.78 | 0.77 | 0.77 | 0.78 |
| BW | 0.99 | 0.82 | 0.52 | 0.82 | 0.79 | 0.79 | 0.82 | 0.80 | 0.81 | 0.81 | 0.85 | 0.83 | 0.82 | 0.79 | 0.77 | 0.78 | 0.80 | 0.84 | 0.81 |
| TMB | 0.99 | 0.82 | 0.52 | 0.80 | 0.76 | 0.77 | 0.78 | 0.78 | 0.78 | 0.78 | 0.79 | 0.78 | 0.78 | 0.78 | 0.76 | 0.78 | 0.78 | 0.78 | 0.78 |
| | Prop. state probabilities > 0.50 | | | | | | | | | | | | | | | | | | |
| null | 1.00 | 0.91 | 0.77 | 0.90 | 0.88 | 0.89 | 0.89 | 0.89 | 0.89 | 0.89 | 0.89 | 0.89 | 0.89 | 0.89 | 0.89 | 0.89 | 0.89 | 0.89 | 0.89 |
| fixed | 1.00 | 0.91 | 0.76 | 0.90 | 0.88 | 0.89 | 0.89 | 0.89 | 0.89 | 0.89 | 0.89 | 0.89 | 0.89 | 0.89 | 0.88 | 0.89 | 0.89 | 0.89 | 0.89 |
| mix2 | 1.00 | 0.91 | 0.76 | 0.90 | 0.88 | 0.89 | 0.89 | 0.89 | 0.89 | 0.89 | 0.89 | 0.89 | 0.89 | 0.89 | 0.88 | 0.89 | 0.89 | 0.89 | 0.89 |
| mix3 | 1.00 | 0.91 | 0.76 | 0.90 | 0.88 | 0.89 | 0.89 | 0.89 | 0.89 | 0.89 | 0.89 | 0.89 | 0.89 | 0.89 | 0.88 | 0.89 | 0.89 | 0.89 | 0.89 |
| mix4 | 1.00 | 0.91 | 0.76 | 0.90 | 0.88 | 0.89 | 0.89 | 0.89 | 0.89 | 0.89 | 0.89 | 0.89 | 0.89 | 0.89 | 0.88 | 0.89 | 0.89 | 0.89 | 0.89 |
| BW | 1.00 | 0.91 | 0.77 | 0.91 | 0.90 | 0.90 | 0.91 | 0.90 | 0.90 | 0.91 | 0.93 | 0.92 | 0.91 | 0.90 | 0.89 | 0.89 | 0.90 | 0.92 | 0.91 |
| TMB | 1.00 | 0.91 | 0.77 | 0.90 | 0.88 | 0.89 | 0.90 | 0.89 | 0.89 | 0.89 | 0.89 | 0.89 | 0.89 | 0.89 | 0.89 | 0.89 | 0.89 | 0.89 | 0.89 |
| | Prop. state probabilities > 0.20 | | | | | | | | | | | | | | | | | | |
| null | 1.00 | 0.96 | 0.95 | 0.97 | 0.97 | 0.97 | 0.97 | 0.97 | 0.97 | 0.97 | 0.98 | 0.98 | 0.97 | 0.97 | 0.96 | 0.97 | 0.97 | 0.97 | 0.97 |
| fixed | 1.00 | 0.96 | 0.93 | 0.97 | 0.96 | 0.96 | 0.96 | 0.96 | 0.96 | 0.96 | 0.97 | 0.97 | 0.97 | 0.96 | 0.96 | 0.97 | 0.96 | 0.96 | 0.96 |
| mix2 | 1.00 | 0.96 | 0.95 | 0.97 | 0.97 | 0.97 | 0.97 | 0.97 | 0.97 | 0.97 | 0.98 | 0.97 | 0.97 | 0.97 | 0.95 | 0.97 | 0.97 | 0.97 | 0.97 |
| mix3 | 1.00 | 0.96 | 0.94 | 0.97 | 0.97 | 0.97 | 0.97 | 0.97 | 0.97 | 0.97 | 0.98 | 0.97 | 0.97 | 0.97 | 0.95 | 0.97 | 0.97 | 0.97 | 0.97 |
| mix4 | 1.00 | 0.96 | 0.94 | 0.97 | 0.97 | 0.97 | 0.97 | 0.97 | 0.97 | 0.97 | 0.98 | 0.97 | 0.97 | 0.97 | 0.95 | 0.97 | 0.97 | 0.97 | 0.97 |
| BW | 1.00 | 0.96 | 0.94 | 0.97 | 0.97 | 0.97 | 0.97 | 0.97 | 0.97 | 0.97 | 0.98 | 0.97 | 0.97 | 0.97 | 0.96 | 0.97 | 0.97 | 0.97 | 0.97 |
| TMB | 1.00 | 0.96 | 0.95 | 0.97 | 0.97 | 0.97 | 0.97 | 0.97 | 0.97 | 0.97 | 0.98 | 0.97 | 0.97 | 0.97 | 0.96 | 0.97 | 0.97 | 0.97 | 0.97 |

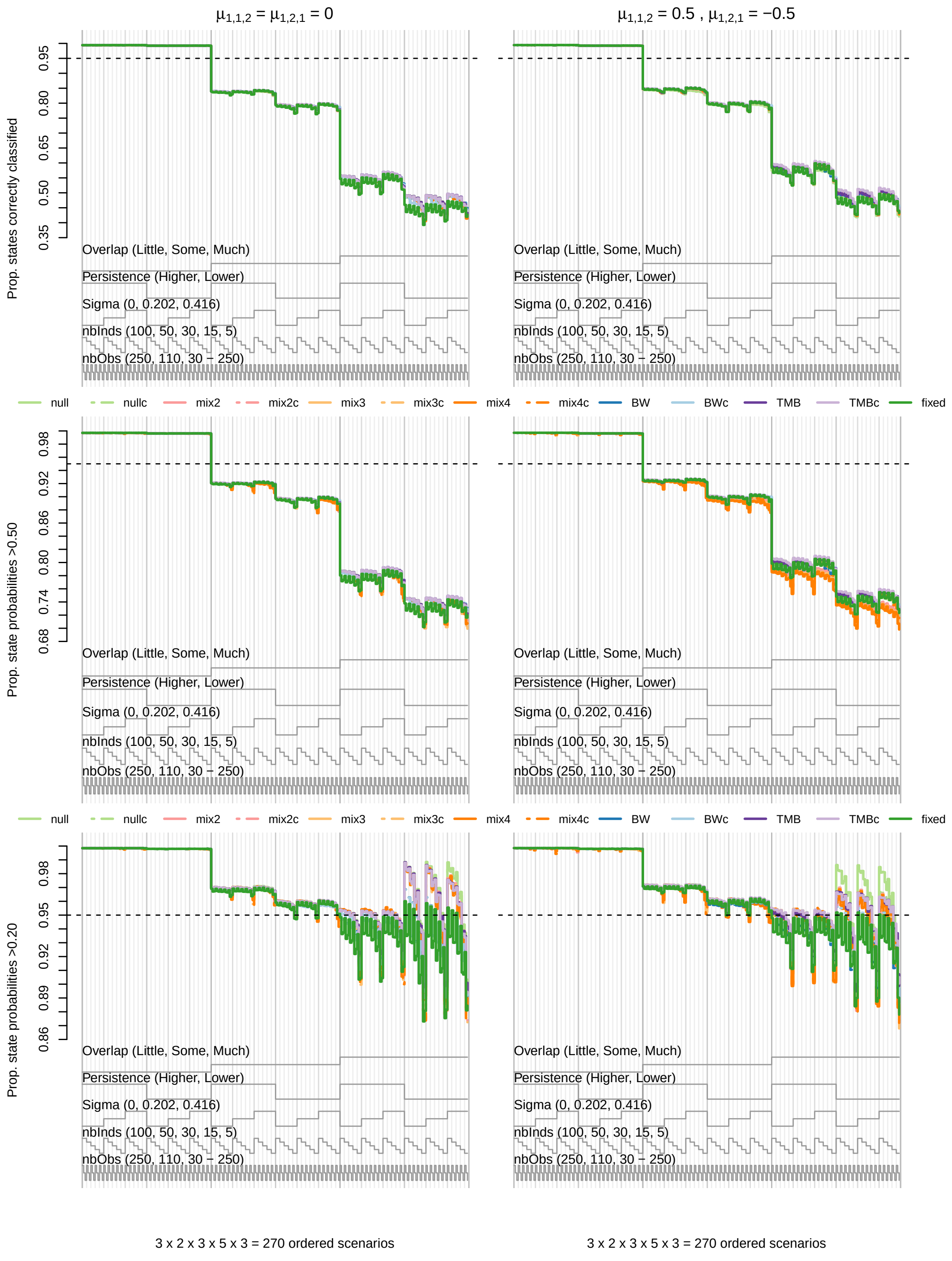

***Figure S17.*** *Nested loop plots for the proportion of Viterbi-decoded states that were correctly classified after accounting for chance agreement (top) and the proportion of estimated state probabilities in which the true state received at least 0.50 (middle) or 0.20 (bottom) probability from 270 simulated scenarios with covariate effects* $\mu_{1,1,2} = \mu_{1,2,1} = 0$ *(left column) or* $\mu_{1,1,2} = 0.5, \mu_{1,2,1} = -0.5$ *(right column). Scenarios are ordered from outer to inner loops by state-dependent distribution overlap, state persistence, individual heterogeneity ("Sigma"), number of individuals ("nbInds"), and length of time series ("nbObs"). Dashed lines and "c" affix indicate models that included covariate effects.*

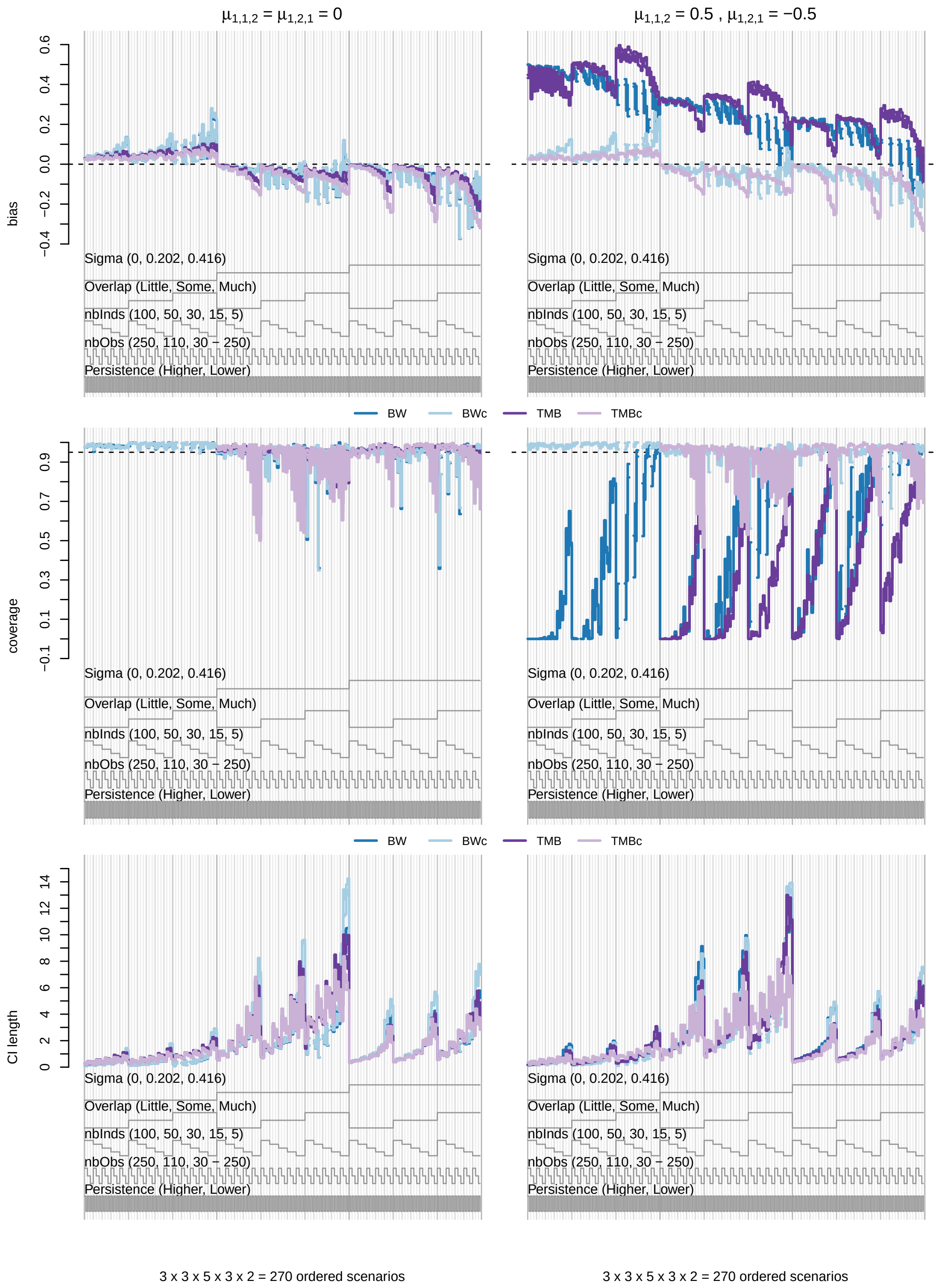

***Figure S18.*** *Nested loop plots for mean bias (top row), 95% confidence interval coverage (middle row), and confidence interval length (bottom row) for* $\sigma_{1,2}$ *and* $\sigma_{2,1}$ *from 270 simulated scenarios with covariate effects. Comparisons are for the BW (blue), BW including covariate effects ("BWc"; light blue), TMB (purple), and TMB including covariate effects ("TMBc"; light purple).*

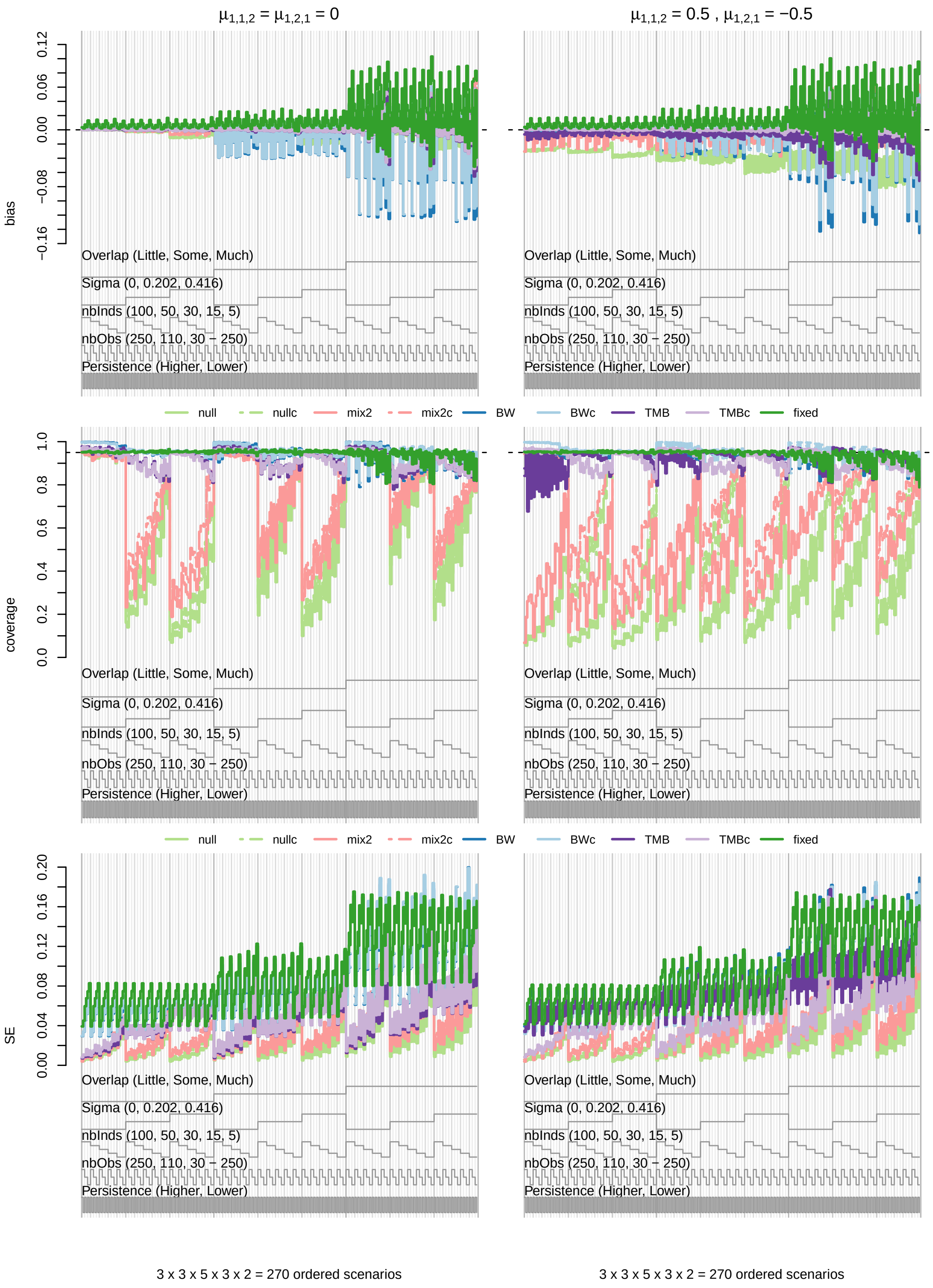

***Figure S19.*** *Nested loop plots for mean bias (top row), 95% confidence interval coverage (middle row), and standard error (SE; bottom row) for* $\gamma_{m,1,2}$ *and* $\gamma_{m,2,1}$ *from 270 simulated scenarios with covariate effects.*

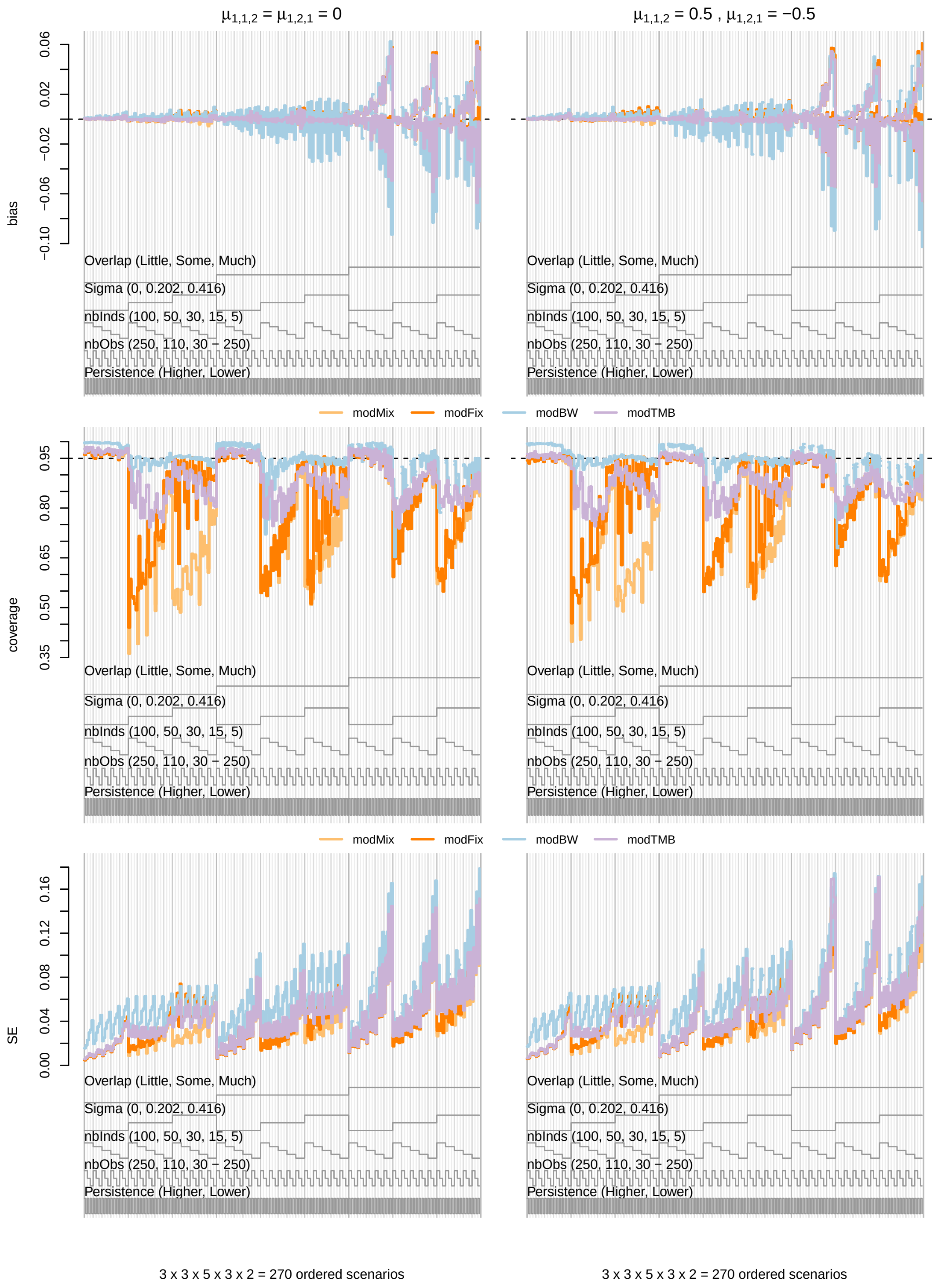

***Figure S20.*** *Nested loop plots for mean bias (top row), 95% confidence interval coverage (middle row), and standard error (SE; bottom row) for* $\gamma_{m,1,2}$ *and* $\gamma_{m,2,1}$ *from 270 simulated scenarios with covariate effects.*

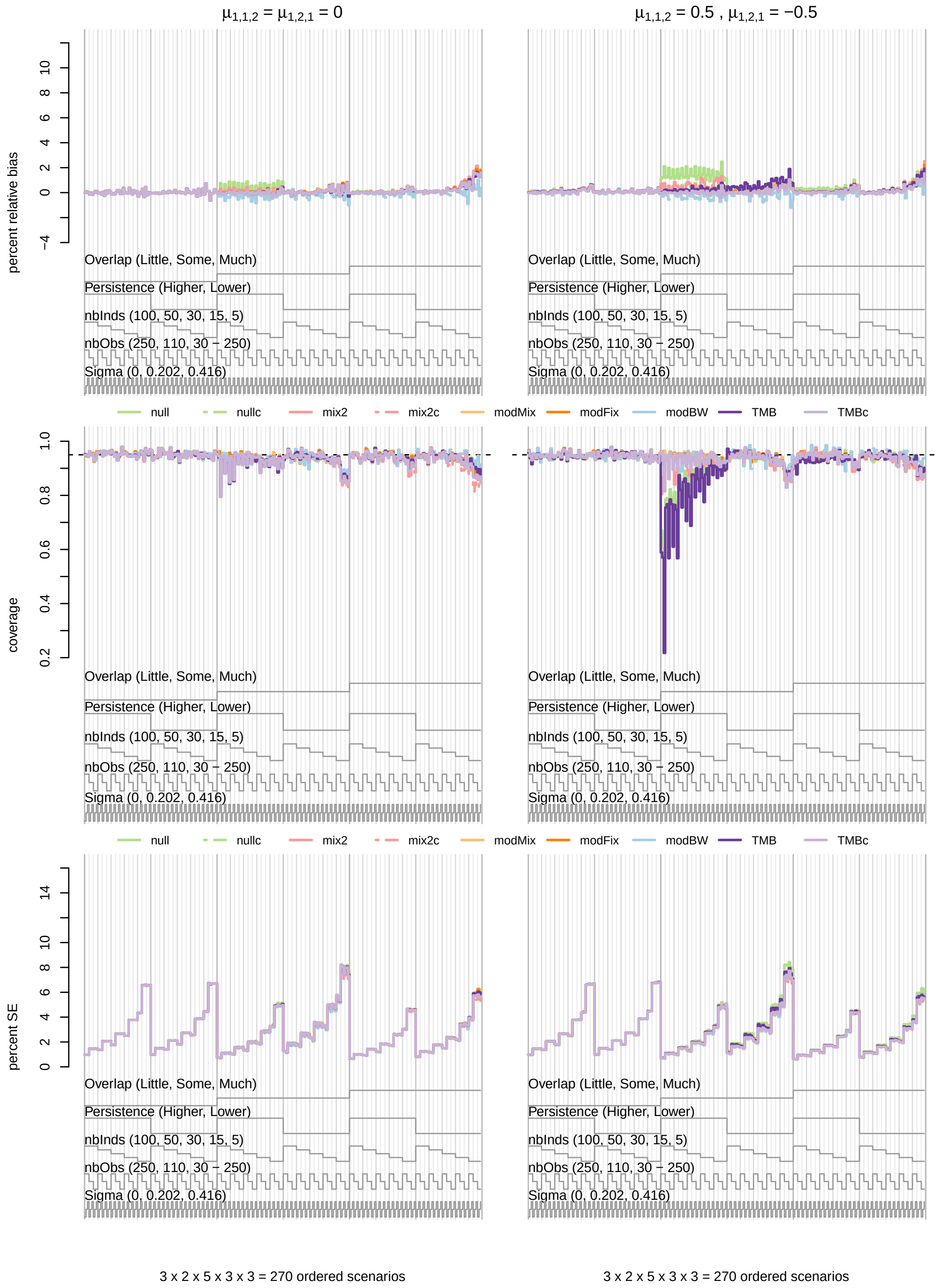

***Figure S21.*** *Nested loop plots for mean percent relative bias (top row), 95% confidence interval coverage (middle row), and percent standard error (SE; bottom row) for* $\mu_1^y$ *from 270 simulated scenarios with covariate effects.*

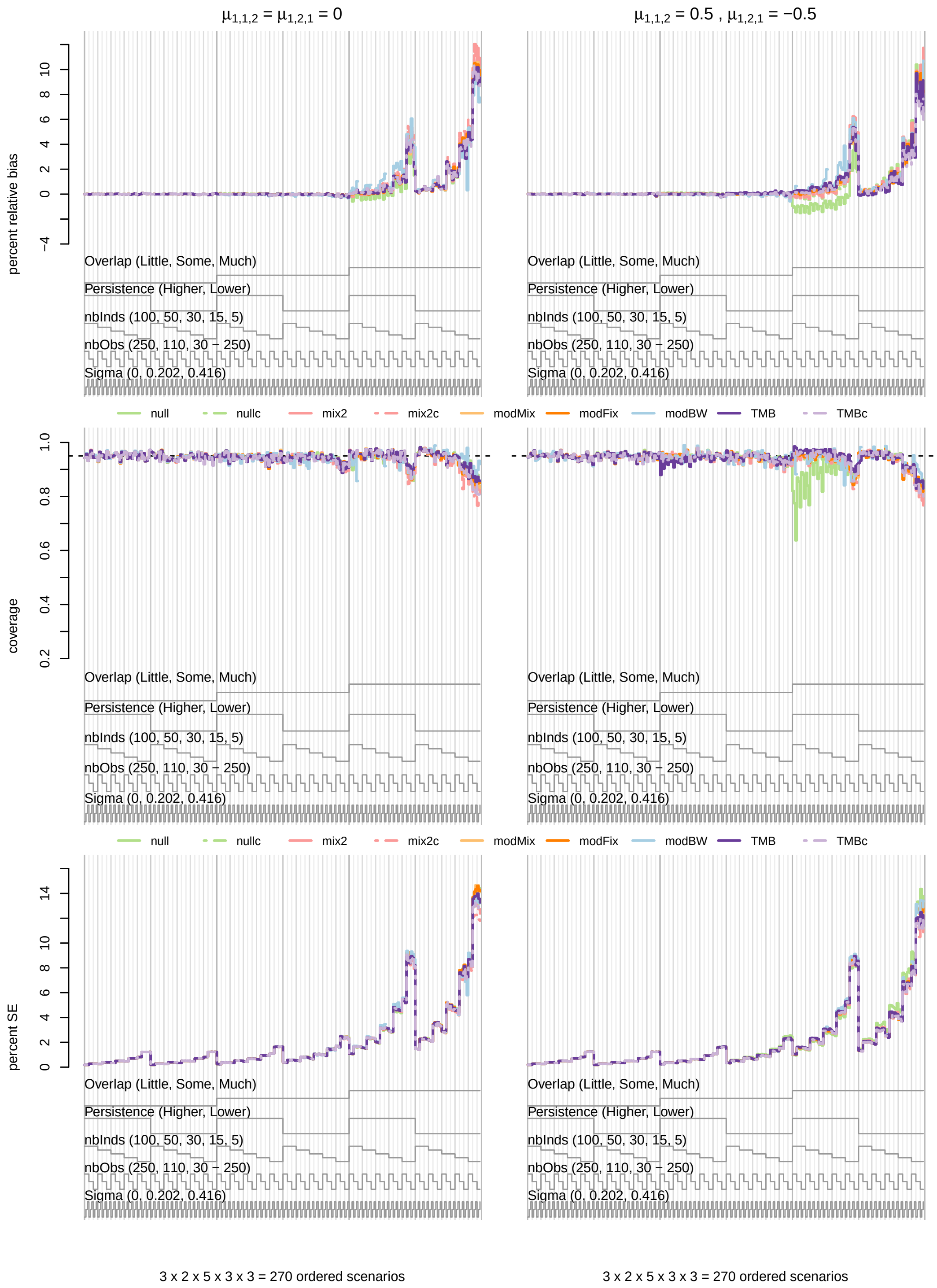

***Figure S22.*** *Nested loop plots for mean percent relative bias (top row), 95% confidence interval coverage (middle row), and percent standard error (SE; bottom row) for* $\mu_2^y$ *from 270 simulated scenarios with covariate effects.*

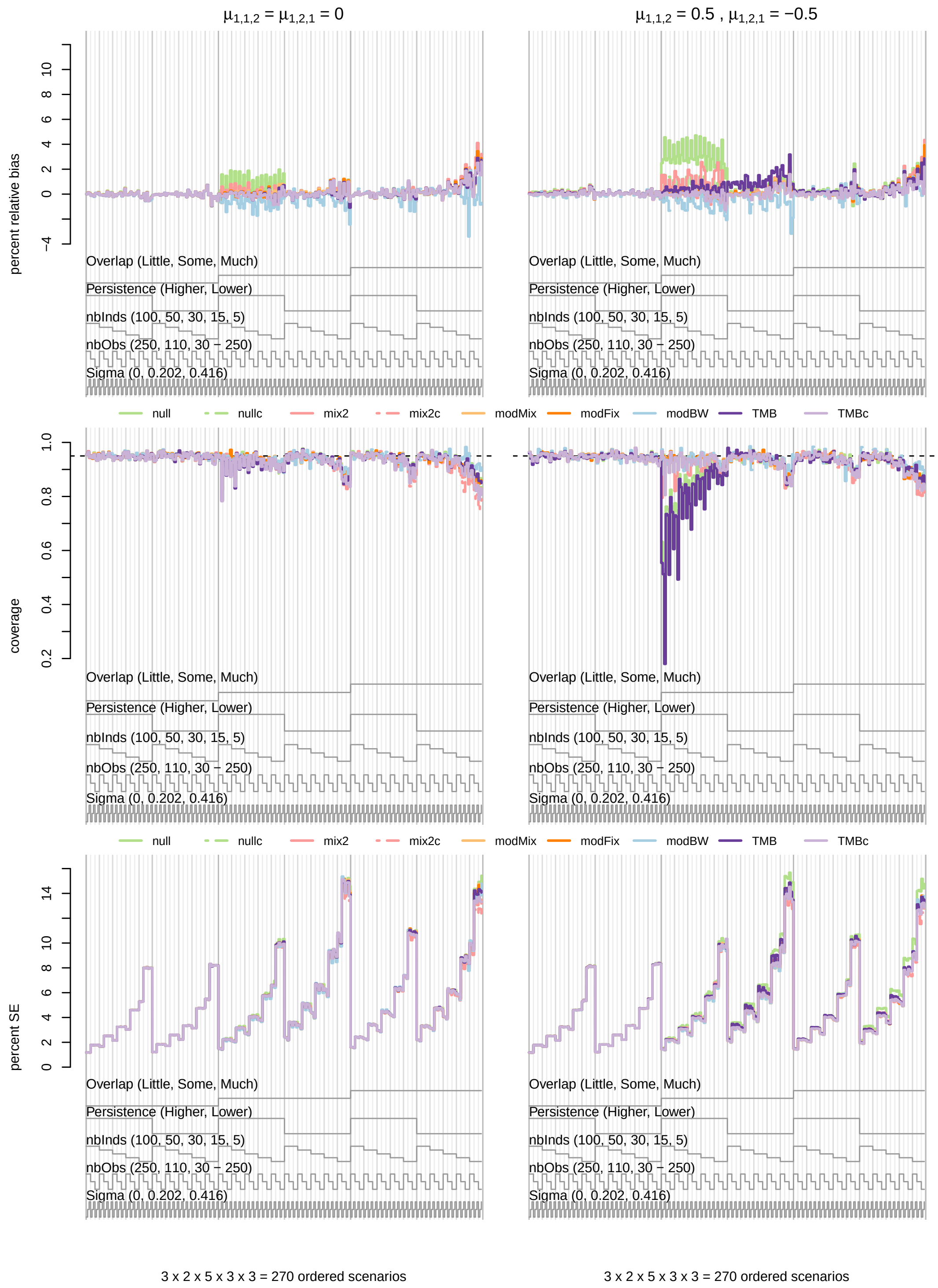

***Figure S23.*** *Nested loop plots for mean percent relative bias (top row), 95% confidence interval coverage (middle row), and percent standard error (SE; bottom row) for* $\sigma_1^y$ *from 270 simulated scenarios with covariate effects.*

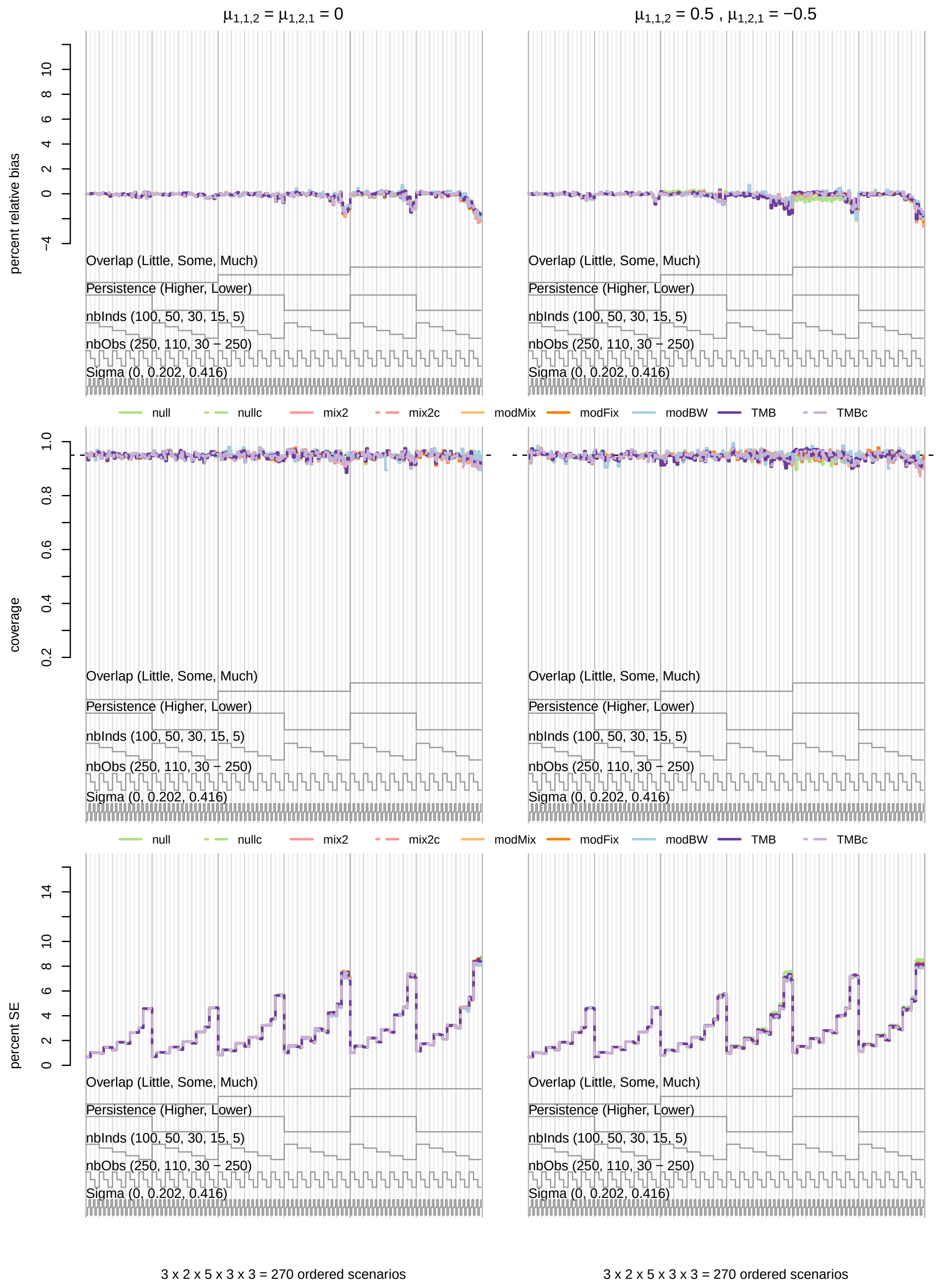

***Figure S24.*** *Nested loop plots for mean percent relative bias (top row), 95% confidence interval coverage (middle row), and percent standard error (SE; bottom row) for* $\sigma_2^y$ *from 270 simulated scenarios with covariate effects.*

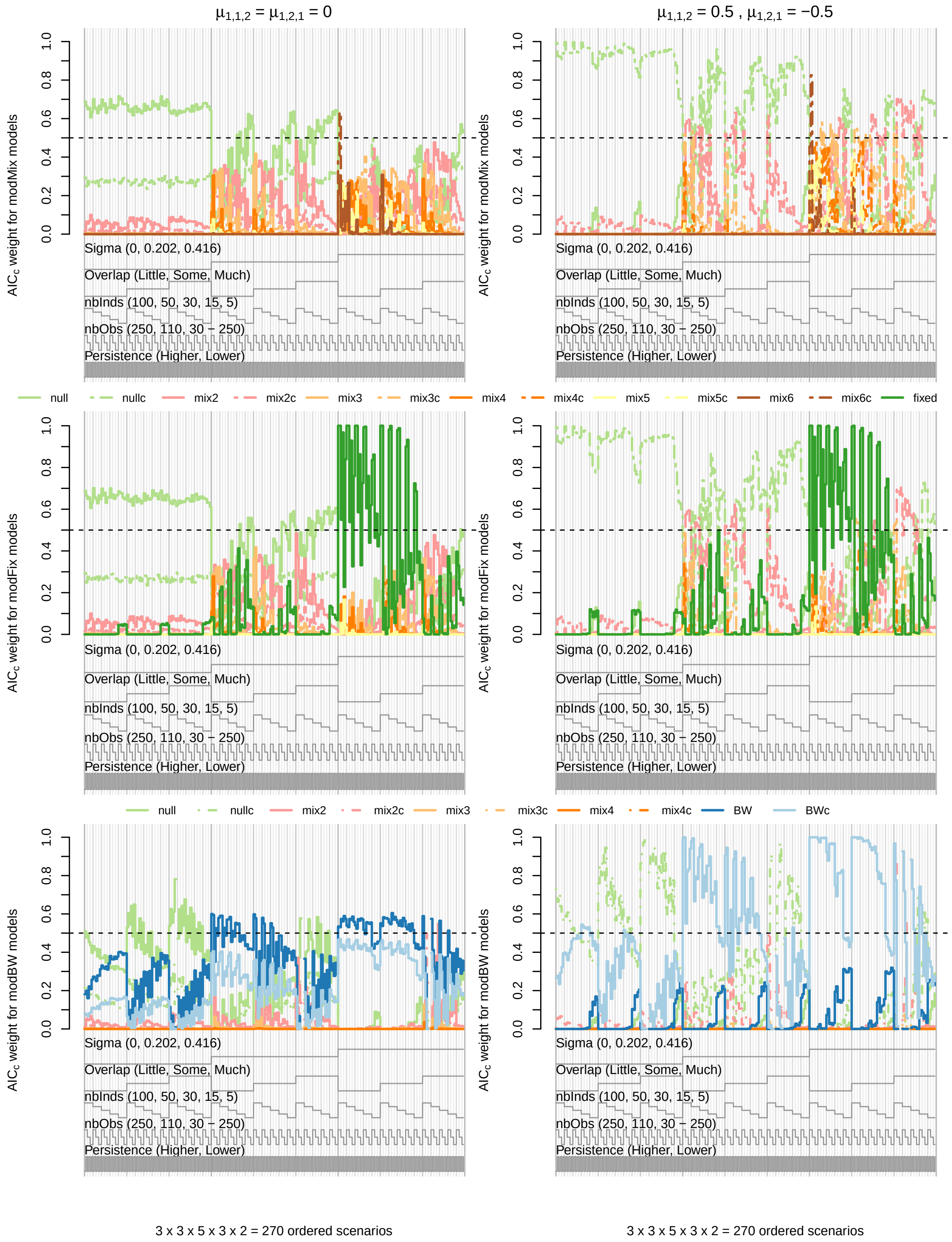

***Figure S25.*** *Nested loop plots for mean AIC$_c$ weights in candidate model sets from 270 simulated scenarios with covariate effects.*

***Table S9.*** *Overall mean bias, 95% confidence interval coverage, and standard error (SE) for covariate effect $\mu_{1,1,2}$ by simulation design points for the covariate effect, state-dependent distribution overlap, state persistence, individual heterogeneity ($\boldsymbol{\sigma}$), number of individuals, and time series length from 540 scenarios with covariate effects.*

| Model | $\mu_{1,1,2}$ | | Overlap | | | State persistence | | $\boldsymbol{\sigma}$ | | | No. individuals ($M$) | | | | | Time series length ($T_m$) | | | Overall |
|---|---|---|---|---|---|---|---|---|---|---|---|---|---|---|---|---|---|---|---|
| | 0 | 0.5 | Little | Some | Much | Higher | Lower | 0 | 0.202 | 0.416 | 100 | 50 | 30 | 15 | 5 | 250 | 110 | 30 - 250 | |
| | | | | | | | | | | Bias | | | | | | | | | |
| null | 0.00 | -0.01 | 0.00 | 0.00 | -0.01 | 0.00 | 0.00 | 0.00 | 0.00 | -0.01 | -0.01 | 0.00 | 0.00 | 0.00 | 0.00 | 0.00 | 0.00 | 0.00 | 0.00 |
| mix2 | 0.00 | -0.01 | -0.01 | -0.01 | 0.00 | 0.00 | -0.01 | 0.00 | 0.00 | -0.01 | 0.00 | 0.00 | 0.00 | 0.00 | -0.03 | -0.01 | 0.00 | -0.01 | -0.01 |
| mix3 | 0.00 | -0.01 | -0.01 | -0.01 | -0.01 | -0.01 | -0.01 | 0.00 | 0.00 | -0.02 | 0.00 | 0.00 | 0.00 | -0.01 | -0.03 | -0.01 | 0.00 | -0.01 | -0.01 |
| mix4 | 0.00 | -0.01 | -0.01 | -0.01 | 0.00 | -0.01 | -0.01 | 0.00 | 0.00 | -0.02 | 0.00 | 0.00 | 0.00 | -0.01 | -0.03 | -0.01 | 0.00 | 0.00 | -0.01 |
| modMix | 0.00 | 0.00 | 0.00 | 0.00 | 0.00 | 0.00 | 0.00 | 0.00 | 0.00 | -0.01 | 0.00 | 0.00 | 0.00 | 0.00 | 0.00 | -0.01 | 0.00 | 0.00 | 0.00 |
| BW | 0.00 | -0.03 | 0.00 | -0.02 | -0.04 | -0.01 | -0.02 | -0.01 | -0.02 | -0.02 | -0.01 | -0.02 | -0.02 | -0.02 | -0.02 | -0.02 | -0.02 | -0.02 | -0.02 |
| modBW | 0.00 | -0.02 | 0.00 | -0.01 | -0.02 | -0.01 | -0.01 | 0.00 | -0.01 | -0.02 | -0.01 | -0.01 | -0.01 | -0.01 | -0.01 | -0.01 | -0.01 | -0.01 | -0.01 |
| TMB | 0.00 | 0.00 | 0.00 | 0.00 | 0.00 | 0.00 | 0.00 | 0.00 | 0.00 | 0.00 | 0.00 | 0.00 | 0.00 | 0.00 | 0.00 | 0.00 | 0.00 | 0.00 | 0.00 |
| modTMB | 0.00 | 0.00 | 0.00 | 0.00 | 0.00 | 0.00 | 0.00 | 0.00 | 0.00 | 0.00 | 0.00 | 0.00 | 0.00 | 0.00 | 0.00 | 0.00 | 0.00 | 0.00 | 0.00 |
| | | | | | | | | | | Coverage | | | | | | | | | |
| null | 0.87 | 0.87 | 0.83 | 0.87 | 0.91 | 0.88 | 0.86 | 0.95 | 0.89 | 0.77 | 0.86 | 0.87 | 0.87 | 0.87 | 0.88 | 0.84 | 0.89 | 0.87 | 0.87 |
| mix2 | 0.90 | 0.90 | 0.87 | 0.90 | 0.94 | 0.91 | 0.90 | 0.96 | 0.93 | 0.83 | 0.89 | 0.90 | 0.90 | 0.91 | 0.92 | 0.87 | 0.92 | 0.91 | 0.90 |
| mix3 | 0.92 | 0.92 | 0.89 | 0.92 | 0.95 | 0.92 | 0.92 | 0.97 | 0.94 | 0.86 | 0.91 | 0.91 | 0.92 | 0.93 | 0.93 | 0.89 | 0.94 | 0.93 | 0.92 |
| mix4 | 0.93 | 0.93 | 0.90 | 0.93 | 0.96 | 0.93 | 0.93 | 0.97 | 0.95 | 0.88 | 0.92 | 0.93 | 0.93 | 0.94 | 0.93 | 0.90 | 0.94 | 0.94 | 0.93 |
| modMix | 0.91 | 0.91 | 0.88 | 0.91 | 0.93 | 0.91 | 0.90 | 0.95 | 0.92 | 0.86 | 0.91 | 0.91 | 0.91 | 0.90 | 0.90 | 0.89 | 0.92 | 0.91 | 0.91 |
| BW | 0.95 | 0.92 | 0.94 | 0.94 | 0.93 | 0.94 | 0.93 | 0.95 | 0.93 | 0.93 | 0.92 | 0.93 | 0.94 | 0.95 | 0.94 | 0.93 | 0.95 | 0.94 | 0.94 |
| modBW | 0.95 | 0.94 | 0.94 | 0.94 | 0.94 | 0.94 | 0.94 | 0.96 | 0.94 | 0.93 | 0.94 | 0.94 | 0.95 | 0.95 | 0.94 | 0.94 | 0.94 | 0.94 | 0.94 |
| TMB | 0.93 | 0.93 | 0.93 | 0.93 | 0.94 | 0.93 | 0.93 | 0.95 | 0.93 | 0.92 | 0.94 | 0.94 | 0.94 | 0.93 | 0.90 | 0.93 | 0.93 | 0.93 | 0.93 |
| modTMB | 0.93 | 0.93 | 0.92 | 0.93 | 0.94 | 0.93 | 0.93 | 0.95 | 0.93 | 0.91 | 0.94 | 0.94 | 0.94 | 0.93 | 0.91 | 0.93 | 0.93 | 0.93 | 0.93 |
| | | | | | | | | | | SE | | | | | | | | | |
| null | 0.10 | 0.12 | 0.07 | 0.09 | 0.16 | 0.11 | 0.11 | 0.11 | 0.11 | 0.11 | 0.04 | 0.06 | 0.08 | 0.12 | 0.24 | 0.08 | 0.12 | 0.13 | 0.11 |
| mix2 | 0.13 | 0.15 | 0.09 | 0.12 | 0.20 | 0.14 | 0.14 | 0.13 | 0.14 | 0.15 | 0.05 | 0.07 | 0.10 | 0.15 | 0.32 | 0.10 | 0.15 | 0.17 | 0.14 |
| mix3 | 0.14 | 0.16 | 0.10 | 0.13 | 0.21 | 0.15 | 0.15 | 0.13 | 0.15 | 0.17 | 0.05 | 0.08 | 0.11 | 0.17 | 0.33 | 0.11 | 0.16 | 0.18 | 0.15 |
| mix4 | 0.14 | 0.17 | 0.10 | 0.14 | 0.22 | 0.15 | 0.15 | 0.13 | 0.15 | 0.18 | 0.06 | 0.08 | 0.11 | 0.17 | 0.34 | 0.11 | 0.16 | 0.18 | 0.15 |
| modMix | 0.12 | 0.13 | 0.09 | 0.11 | 0.18 | 0.13 | 0.12 | 0.11 | 0.12 | 0.14 | 0.05 | 0.07 | 0.10 | 0.14 | 0.26 | 0.09 | 0.13 | 0.14 | 0.12 |
| BW | 0.12 | 0.14 | 0.10 | 0.12 | 0.19 | 0.13 | 0.13 | 0.11 | 0.12 | 0.15 | 0.04 | 0.06 | 0.09 | 0.13 | 0.28 | 0.10 | 0.14 | 0.16 | 0.13 |
| modBW | 0.12 | 0.14 | 0.10 | 0.12 | 0.20 | 0.13 | 0.13 | 0.11 | 0.12 | 0.16 | 0.04 | 0.07 | 0.09 | 0.14 | 0.29 | 0.10 | 0.14 | 0.16 | 0.13 |
| TMB | 0.12 | 0.13 | 0.09 | 0.11 | 0.18 | 0.13 | 0.12 | 0.11 | 0.12 | 0.14 | 0.05 | 0.07 | 0.10 | 0.14 | 0.27 | 0.10 | 0.13 | 0.14 | 0.13 |
| modTMB | 0.12 | 0.13 | 0.09 | 0.11 | 0.18 | 0.13 | 0.13 | 0.11 | 0.12 | 0.15 | 0.05 | 0.07 | 0.10 | 0.14 | 0.27 | 0.10 | 0.13 | 0.15 | 0.13 |

***Table S10.*** *Overall mean bias, 95% confidence interval coverage, and standard error (SE) for covariate effect $\mu_{1,2,1}$ by simulation design points for the covariate effect, state-dependent distribution overlap, state persistence, individual heterogeneity ($\boldsymbol{\sigma}$), number of individuals, and time series length from 540 scenarios with covariate effects.*

| Model | $\mu_{1,2,1}$ | | Overlap | | | State persistence | | $\boldsymbol{\sigma}$ | | | No. individuals ($M$) | | | | | Time series length ($T_m$) | | | Overall |
|---|---|---|---|---|---|---|---|---|---|---|---|---|---|---|---|---|---|---|---|
| | 0 | -0.5 | Little | Some | Much | Higher | Lower | 0 | 0.202 | 0.416 | 100 | 50 | 30 | 15 | 5 | 250 | 110 | 30 - 250 | |
| | | | | | | | | | Bias | | | | | | | | | | |
| null | 0.00 | 0.01 | 0.00 | 0.00 | 0.01 | 0.01 | 0.00 | 0.00 | 0.00 | 0.01 | 0.01 | 0.01 | 0.00 | 0.01 | 0.00 | 0.01 | 0.00 | 0.01 | 0.00 |
| mix2 | 0.00 | 0.01 | 0.01 | 0.01 | 0.00 | 0.01 | 0.01 | 0.00 | 0.00 | 0.02 | 0.00 | 0.00 | 0.00 | 0.01 | 0.02 | 0.01 | 0.00 | 0.01 | 0.01 |
| mix3 | 0.00 | 0.01 | 0.01 | 0.01 | 0.00 | 0.01 | 0.01 | 0.00 | 0.00 | 0.02 | 0.00 | 0.00 | 0.00 | 0.01 | 0.03 | 0.01 | 0.00 | 0.00 | 0.01 |
| mix4 | 0.00 | 0.01 | 0.01 | 0.01 | 0.00 | 0.01 | 0.01 | 0.00 | 0.00 | 0.02 | 0.00 | 0.00 | 0.00 | 0.01 | 0.03 | 0.01 | 0.00 | 0.00 | 0.01 |
| modMix | 0.00 | 0.01 | 0.00 | 0.00 | 0.00 | 0.00 | 0.00 | 0.00 | 0.00 | 0.01 | 0.00 | 0.00 | 0.00 | 0.01 | 0.01 | 0.01 | 0.00 | 0.00 | 0.00 |
| BW | 0.00 | 0.04 | 0.00 | 0.01 | 0.05 | 0.02 | 0.02 | 0.02 | 0.02 | 0.02 | 0.01 | 0.02 | 0.02 | 0.02 | 0.02 | 0.02 | 0.02 | 0.02 | 0.02 |
| modBW | 0.00 | 0.02 | 0.00 | 0.01 | 0.03 | 0.01 | 0.01 | 0.01 | 0.01 | 0.02 | 0.01 | 0.01 | 0.01 | 0.01 | 0.01 | 0.01 | 0.01 | 0.01 | 0.01 |
| TMB | 0.00 | 0.00 | 0.00 | 0.00 | 0.00 | 0.00 | 0.00 | 0.00 | 0.00 | 0.00 | 0.00 | 0.00 | 0.00 | 0.00 | 0.00 | 0.00 | 0.00 | 0.00 | 0.00 |
| modTMB | 0.00 | 0.00 | 0.00 | 0.00 | 0.00 | 0.00 | 0.00 | 0.00 | 0.00 | 0.00 | 0.00 | 0.00 | 0.00 | 0.00 | 0.00 | 0.00 | 0.00 | 0.00 | 0.00 |
| | | | | | | | | | Coverage | | | | | | | | | | |
| null | 0.87 | 0.87 | 0.83 | 0.87 | 0.91 | 0.88 | 0.86 | 0.94 | 0.89 | 0.77 | 0.86 | 0.87 | 0.87 | 0.87 | 0.88 | 0.84 | 0.89 | 0.87 | 0.87 |
| mix2 | 0.90 | 0.90 | 0.87 | 0.90 | 0.94 | 0.91 | 0.90 | 0.96 | 0.93 | 0.83 | 0.89 | 0.90 | 0.90 | 0.91 | 0.92 | 0.88 | 0.92 | 0.91 | 0.90 |
| mix3 | 0.92 | 0.92 | 0.89 | 0.92 | 0.95 | 0.92 | 0.92 | 0.96 | 0.94 | 0.86 | 0.91 | 0.92 | 0.92 | 0.93 | 0.93 | 0.89 | 0.94 | 0.93 | 0.92 |
| mix4 | 0.93 | 0.93 | 0.90 | 0.93 | 0.95 | 0.93 | 0.93 | 0.97 | 0.95 | 0.88 | 0.92 | 0.93 | 0.93 | 0.94 | 0.93 | 0.90 | 0.94 | 0.94 | 0.93 |
| modMix | 0.91 | 0.91 | 0.89 | 0.91 | 0.93 | 0.91 | 0.90 | 0.94 | 0.92 | 0.86 | 0.91 | 0.91 | 0.91 | 0.90 | 0.90 | 0.89 | 0.92 | 0.91 | 0.91 |
| BW | 0.95 | 0.92 | 0.94 | 0.94 | 0.93 | 0.94 | 0.94 | 0.95 | 0.94 | 0.93 | 0.93 | 0.94 | 0.94 | 0.94 | 0.94 | 0.93 | 0.94 | 0.94 | 0.94 |
| modBW | 0.95 | 0.94 | 0.94 | 0.94 | 0.94 | 0.94 | 0.94 | 0.95 | 0.94 | 0.93 | 0.94 | 0.94 | 0.94 | 0.94 | 0.93 | 0.94 | 0.94 | 0.94 | 0.94 |
| TMB | 0.93 | 0.93 | 0.93 | 0.93 | 0.93 | 0.93 | 0.93 | 0.95 | 0.93 | 0.92 | 0.94 | 0.94 | 0.94 | 0.93 | 0.91 | 0.93 | 0.94 | 0.93 | 0.93 |
| modTMB | 0.93 | 0.93 | 0.93 | 0.93 | 0.93 | 0.93 | 0.93 | 0.95 | 0.93 | 0.91 | 0.94 | 0.94 | 0.93 | 0.93 | 0.91 | 0.93 | 0.93 | 0.93 | 0.93 |
| | | | | | | | | | SE | | | | | | | | | | |
| null | 0.11 | 0.12 | 0.07 | 0.09 | 0.18 | 0.12 | 0.11 | 0.11 | 0.11 | 0.12 | 0.04 | 0.06 | 0.08 | 0.12 | 0.26 | 0.08 | 0.13 | 0.13 | 0.11 |
| mix2 | 0.13 | 0.16 | 0.09 | 0.12 | 0.22 | 0.14 | 0.15 | 0.13 | 0.14 | 0.16 | 0.05 | 0.07 | 0.10 | 0.16 | 0.34 | 0.10 | 0.16 | 0.17 | 0.14 |
| mix3 | 0.14 | 0.17 | 0.10 | 0.13 | 0.23 | 0.15 | 0.16 | 0.14 | 0.15 | 0.17 | 0.06 | 0.08 | 0.11 | 0.17 | 0.35 | 0.11 | 0.17 | 0.19 | 0.15 |
| mix4 | 0.15 | 0.17 | 0.10 | 0.14 | 0.24 | 0.16 | 0.16 | 0.14 | 0.16 | 0.18 | 0.06 | 0.09 | 0.12 | 0.18 | 0.36 | 0.11 | 0.17 | 0.19 | 0.16 |
| modMix | 0.12 | 0.14 | 0.09 | 0.11 | 0.19 | 0.13 | 0.13 | 0.12 | 0.12 | 0.15 | 0.05 | 0.07 | 0.10 | 0.14 | 0.28 | 0.10 | 0.14 | 0.15 | 0.13 |
| BW | 0.13 | 0.14 | 0.10 | 0.11 | 0.22 | 0.14 | 0.13 | 0.12 | 0.13 | 0.16 | 0.04 | 0.07 | 0.09 | 0.14 | 0.30 | 0.11 | 0.14 | 0.16 | 0.13 |
| modBW | 0.13 | 0.14 | 0.10 | 0.11 | 0.22 | 0.14 | 0.13 | 0.12 | 0.13 | 0.16 | 0.04 | 0.07 | 0.09 | 0.14 | 0.30 | 0.11 | 0.15 | 0.16 | 0.14 |
| TMB | 0.12 | 0.14 | 0.09 | 0.11 | 0.19 | 0.13 | 0.13 | 0.12 | 0.12 | 0.15 | 0.05 | 0.08 | 0.10 | 0.14 | 0.28 | 0.10 | 0.14 | 0.15 | 0.13 |
| modTMB | 0.12 | 0.14 | 0.09 | 0.11 | 0.19 | 0.13 | 0.13 | 0.12 | 0.12 | 0.15 | 0.05 | 0.08 | 0.10 | 0.14 | 0.28 | 0.10 | 0.14 | 0.15 | 0.13 |

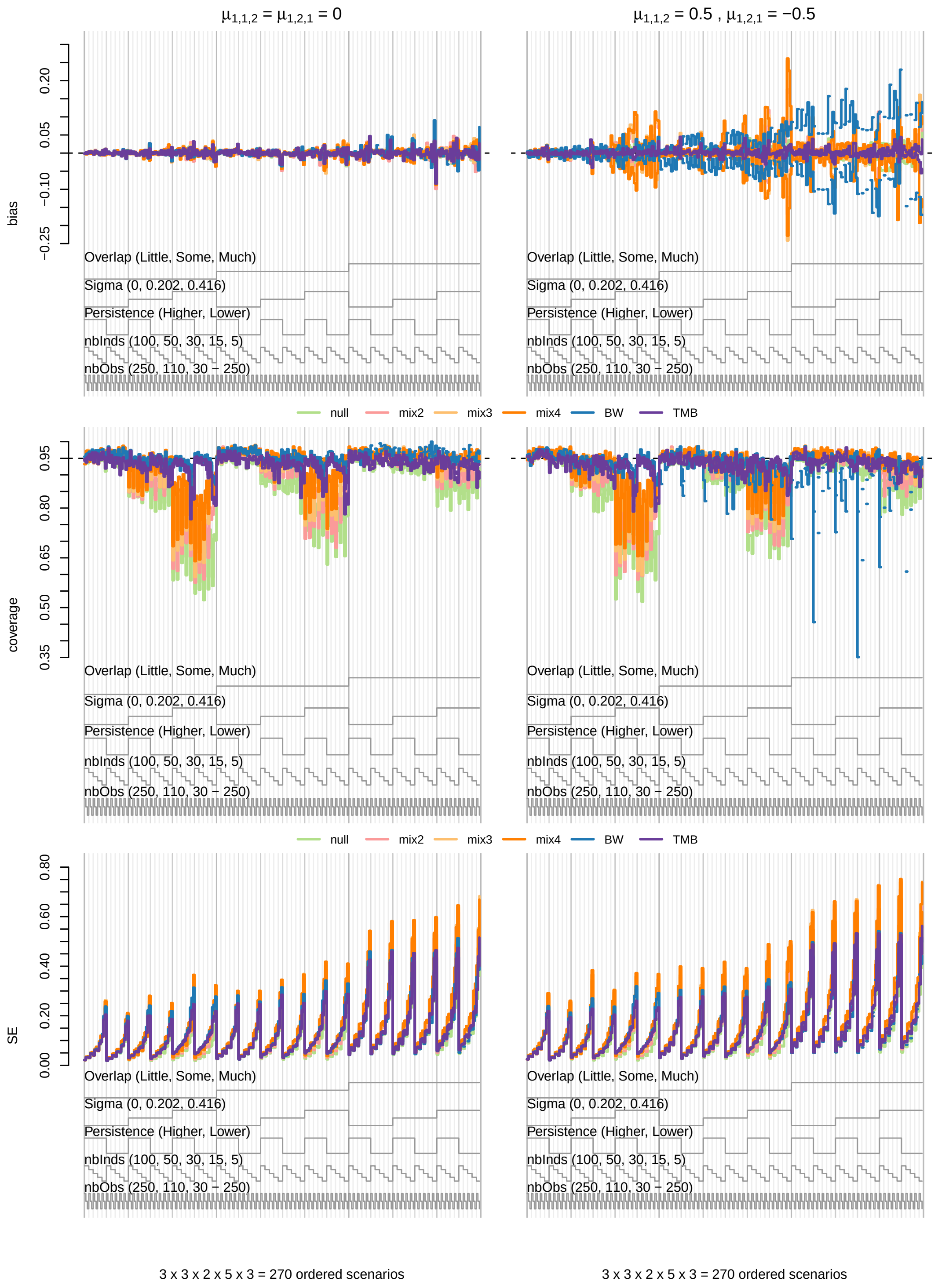

***Figure S26.*** *Nested loop plots for median bias (top row), mean 95% confidence interval coverage (middle row), and median standard error (SE; bottom row) for covariate effects* $\mu_{1,1,2}$ *and* $\mu_{1,2,1}$ *from 270 simulated scenarios. Comparisons are for the null (light green), mix2 (pink), mix3 (light orange), mix4 (dark orange), BW (blue), and TMB (purple) models.*

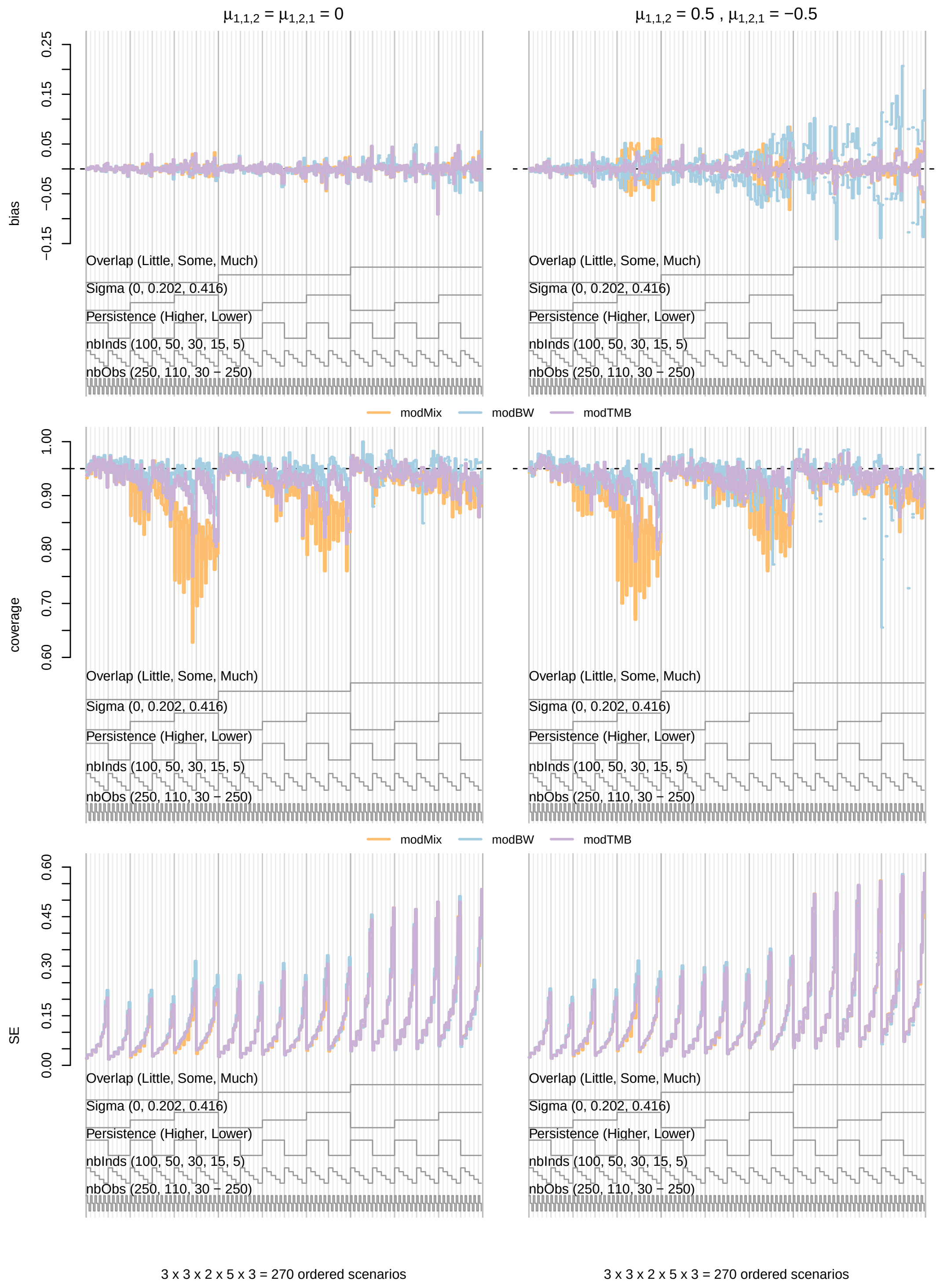

***Figure S27.*** *Nested loop plots for median bias (top row), mean 95% confidence interval coverage (middle row), and median standard error (SE; bottom row) for covariate effects* $\mu_{1,1,2}$ *and* $\mu_{1,2,1}$ *from 270 simulated scenarios. Comparisons are for the* $AIC_c$ *model-averaged modMix (light orange), modBW (light blue), and modTMB (light purple) models.*
