## Supplementary material for "Worth the effort? A practical examination of random effects in hidden Markov models for animal telemetry data": Code: HMM_recov.cpp

```
#include <TMB.hpp>
#include <cmath>

using namespace density;
using std::sqrt;

template <class Type>
int which_max(vector<Type> x){
  int first = 0;
  int last = x.size()-1;
  if (first==last) return last;
  int largest = first;
  
  while (++first<last+1)
    if (x(largest)<x(first))    
      largest=first;
    return largest;
}


/* Numerically stable forward algorithm based on http://bozeman.genome.washington.edu/compbio/mbt599_2006/hmm_scaling_revised.pdf */
template<class Type>
Type forward_alg(vector<Type> delta, array<Type> trMat, matrix<Type> lnProbs, int nbSteps, vector<int> aInd) {
  int nbStates = trMat.cols();
  Type logalpha;
  matrix<Type> ltrMat(nbStates,nbStates);
  vector<Type> ldeltaG(nbStates);
  vector<Type> lalpha(nbStates);
  vector<Type> lnewalpha(nbStates);
  Type jnll = 0.0;
  Type sumalpha;
  int nbAnimals = aInd.size();
  int animal = 0;
  for(int t=0; t < nbSteps; t++){
    if(animal<nbAnimals && t==aInd(animal)){
      sumalpha = -INFINITY;
      for(int j=0; j < nbStates; j++){
        ldeltaG(j) = -INFINITY;
        for(int i=0; i < nbStates; i++){
          ltrMat(i,j) = log(trMat(animal,i,j));
          ldeltaG(j) = logspace_add(ldeltaG(j),log(delta(i))+ltrMat(i,j));
        }
        lalpha(j) = ldeltaG(j)+lnProbs(t,j);
        sumalpha  = logspace_add(sumalpha,lalpha(j));
      }
      jnll -= sumalpha;
      lalpha -= sumalpha;
      animal++;
    } else {
      sumalpha = -INFINITY;
      for(int j=0; j < nbStates; j++){
        logalpha = -INFINITY;
        for(int i=0; i < nbStates; i++){
          logalpha = logspace_add(logalpha,lalpha(i)+ltrMat(i,j));
        }
        lnewalpha(j) = logalpha + lnProbs(t,j);
        sumalpha  = logspace_add(sumalpha,lnewalpha(j));
      }
      jnll -= sumalpha;
      lalpha = lnewalpha - sumalpha;
    }
  }
  return jnll;
}

/* Numerically stable viterbi based on Rabiner 1989 IEEE 77(2):257-286 */
template<class Type>
vector<int> viterbi(vector<Type> delta, array<Type> trMat, matrix<Type> lnProbs, int nbSteps, vector<int> aInd) {
  int nbStates = trMat.cols();
  matrix<Type> phi(nbSteps,nbStates);
  matrix<int> psi(nbSteps,nbStates);
  vector<Type> tmpphi(nbStates);
  vector<int> states(nbSteps);
  states.setOnes();
  matrix<Type> ltrMat(nbStates,nbStates);
  vector<Type> ldeltaG(nbStates);
  int nbAnimals = aInd.size();
  int animal = 0;
  for(int t=0; t < nbSteps; t++){
    if(animal<nbAnimals && t==aInd(animal)){
      for(int j=0; j < nbStates; j++){
        ldeltaG(j) = -INFINITY;
        for(int i=0; i < nbStates; i++){
          ltrMat(i,j) = log(trMat(animal,i,j));
          ldeltaG(j) = logspace_add(ldeltaG(j),log(delta(i))+ltrMat(i,j));
        }
        psi(t,j) = 0;
        phi(t,j) = ldeltaG(j)+lnProbs(t,j);
      }
      animal++;
    } else {
      for(int j=0; j < nbStates; j++){
        for(int i=0; i < nbStates; i++){
          tmpphi(i) = phi(t-1,i)+ltrMat(i,j);
        }
        psi(t,j) = which_max(tmpphi);
        phi(t,j) = tmpphi(psi(t,j))+lnProbs(t,j);
      }
    }
  }
  
  states(nbSteps-1) += which_max(vector<Type>(phi.row(nbSteps-1)));
  animal = nbAnimals-1;
  for(int t=(nbSteps-2); t >= 0 ; t--){
    if(animal>0 && t==(aInd(animal)-1)){
      states(t) += which_max(vector<Type>(phi.row(t)));
      animal--;
    } else {
      states(t) += psi(t+1,states(t+1)-1);
    }
  }
  return states;
}

/* Numerically stable forward log-probabilities based on http://bozeman.genome.washington.edu/compbio/mbt599_2006/hmm_scaling_revised.pdf */
template<class Type>
matrix<Type> logAlpha(vector<Type> delta, array<Type> trMat, matrix<Type> lnProbs, int nbSteps, vector<int> aInd) {
  int nbStates = trMat.cols();
  Type logalpha;
  matrix<Type> elnalpha(nbSteps,nbStates);
  matrix<Type> ltrMat(nbStates,nbStates);
  vector<Type> ldeltaG(nbStates);
  int nbAnimals = aInd.size();
  int animal = 0;
  for(int t=0; t < nbSteps; t++){
    if(animal<nbAnimals && t==aInd(animal)){
      for(int j=0; j < nbStates; j++){
        ldeltaG(j) = -INFINITY;
        for(int i=0; i < nbStates; i++){
          ltrMat(i,j) = log(trMat(animal,i,j));
          ldeltaG(j) = logspace_add(ldeltaG(j),log(delta(i))+ltrMat(i,j));
        }
        elnalpha(t,j) = ldeltaG(j)+lnProbs(t,j);
      }
      animal++;
    } else {
      for(int j=0; j < nbStates; j++){
        logalpha = -INFINITY;
        for(int i=0; i < nbStates; i++){
          logalpha = logspace_add(logalpha,elnalpha(t-1,i)+ltrMat(i,j));
        }
        elnalpha(t,j) = logalpha+lnProbs(t,j);
      }
    }
  }
  return elnalpha;
}

/* Numerically stable backward log-probabilities based on http://bozeman.genome.washington.edu/compbio/mbt599_2006/hmm_scaling_revised.pdf */
template<class Type>
matrix<Type> logBeta(vector<Type> delta, array<Type> trMat, matrix<Type> lnProbs, int nbSteps, vector<int> aInd) {
  int nbStates = trMat.cols();
  Type logbeta;
  matrix<Type> elnbeta(nbSteps,nbStates);
  matrix<Type> ltrMat(nbStates,nbStates);
  int nbAnimals = aInd.size();
  
  int animal = nbAnimals-1;
  for(int j=0; j < nbStates; j++){
    elnbeta(nbSteps-1,j) = Type(0.0);
    for(int i=0; i < nbStates; i++){
      ltrMat(i,j) = log(trMat(animal,i,j));
    }
  }
  for(int t=(nbSteps-2); t >= 0 ; t--){
    if(animal>0 && t==(aInd(animal)-1)){
      animal--;
      for(int j=0; j < nbStates; j++){
        elnbeta(t,j) = Type(0.0);
        for(int i=0; i < nbStates; i++){
          ltrMat(i,j) = log(trMat(animal,i,j));
        }
      }
    } else {
      for(int i=0; i < nbStates; i++){
        logbeta = -INFINITY;
        for(int j=0; j < nbStates; j++){
          logbeta = logspace_add(logbeta,ltrMat(i,j)+lnProbs(t+1,j)+elnbeta(t+1,j));
        }
        elnbeta(t,i) = logbeta;
      }
    }
  }
  return elnbeta;
}

/* Numerically stable state probabilities based on http://bozeman.genome.washington.edu/compbio/mbt599_2006/hmm_scaling_revised.pdf */
template<class Type>
matrix<Type> stProbs(matrix<Type> elnalpha, matrix<Type> elnbeta, int nbSteps) {
  int nbStates = elnalpha.cols();
  Type normalizer;
  matrix<Type> gamma(nbSteps,nbStates);
  
  for(int t=0; t < nbSteps; t++){
    normalizer = -INFINITY;
    for(int i=0; i < nbStates; i++){
      gamma(t,i) = elnalpha(t,i)+elnbeta(t,i);
      normalizer = logspace_add(normalizer,gamma(t,i));
    }
    for(int i=0; i < nbStates; i++){
      gamma(t,i) = exp(gamma(t,i)-normalizer);
    }
  }
  return gamma;
}

template<class Type>
Type objective_function<Type>::operator() ()
{
  // DATA
  DATA_VECTOR(data);	            //  observations
  DATA_VECTOR(cov);               // covariates
  DATA_INTEGER(nbStates);        // number of states
  DATA_INTEGER(nbSteps);          // number of observations
  DATA_IVECTOR(aInd);          // indicator for first observation of each individual
  
  // OBSERVATION PARAMETERS
  PARAMETER_VECTOR(l_mu); 				// gamma mean (log scale)
  PARAMETER_VECTOR(l_sd);     	// gamma sd (log scale)
  
  // initial state and t.p.m. random effect parameters
  PARAMETER_VECTOR(l_delta); // initial distribution (mlogit scale) - should be of length nbStates - 1
  PARAMETER_VECTOR(mu_0); // t.p.m. (mlogit scale) - should be 1 x (nbStates-1) * nbStates
  PARAMETER_VECTOR(mu_1); // covariate effect
  PARAMETER_VECTOR(l_sigma); // sd for random effects (log scale)
  
  // random effects
  PARAMETER_MATRIX(epsilon);

  
  // Transform parameters
  vector<Type> shape(nbStates);
  vector<Type> scale(nbStates);
  vector<Type> mu(nbStates);
  vector<Type> sd(nbStates);
  vector<Type> sigma(nbStates);
  for(int i=0; i < nbStates; i++){
    mu(i) = exp(l_mu(i));
    sd(i) = exp(l_sd(i));
    shape(i) = mu(i) * mu(i) / (sd(i) * sd(i));
    scale(i) = sd(i) * sd(i) / mu(i);
    sigma(i) = exp(l_sigma(i));
  }
  
  int nbAnimals = aInd.size();
  
  /* Define likelihood */
  Type nll=0.0;
  
  // initial distribution
  vector<Type> delta(nbStates);
  delta(0) = Type(1.0);
  for(int i=1; i < nbStates; i++){
    delta(i) = exp(l_delta(i-1));
  }
  delta /= delta.sum();
  
  // tpm
  array<Type> trMat(nbAnimals,nbStates,nbStates);
  Type denom;
  Type epsmean=0.0;
  int cpt;
  for(int s=0;s<nbAnimals;s++){
    cpt = 0;
    for(int i=0;i<nbStates;i++){
      denom = Type(0.0);
      for(int j=0;j<nbStates;j++){
        if(i==j) {
          trMat(s,i,j) = Type(1.0);
          cpt++;
        } else {
          trMat(s,i,j) = exp(epsilon(s,i*nbStates+j-cpt));
          epsmean = mu_0(i*nbStates+j-cpt)+mu_1(i*nbStates+j-cpt)*cov(s);
          nll -= dnorm(epsilon(s,i*nbStates+j-cpt),epsmean,sigma(i),true);
        }
        denom += trMat(s,i,j);
      }
      for(int j=0;j<nbStates;j++){
        trMat(s,i,j) /= denom;
      }
    }
  }
  
  matrix<Type> allProbs(nbSteps,nbStates);
  allProbs.setZero();
  
  // observations
  
  for(int state = 0; state < nbStates; state++){
    for(int t=0; t < nbSteps; t++){
      allProbs(t,state) = dgamma(data(t),shape(state),scale(state),1);
    }
  }
  
  REPORT(allProbs);
  
  // forward algorithm
  Type hmmll = forward_alg(delta, trMat, allProbs, nbSteps, aInd);
  REPORT(hmmll);
  nll += hmmll;
  
  ADREPORT(mu);
  ADREPORT(sd);
  
  if(nbStates>1){
    vector<int> states = viterbi(delta, trMat, allProbs, nbSteps, aInd);
    REPORT(states);
    matrix<Type> lalpha = logAlpha(delta, trMat, allProbs, nbSteps, aInd);
    matrix<Type> lbeta = logBeta(delta, trMat, allProbs, nbSteps, aInd);
    matrix<Type> stateProbs = stProbs(lalpha, lbeta, nbSteps); 
    REPORT(stateProbs);
    ADREPORT(delta);
    ADREPORT(trMat);
    ADREPORT(sigma);
  }
  
  return nll;
}
```
