## Supplementary material for "Worth the effort? A practical examination of random effects in hidden Markov models for animal telemetry data": Code: pilotWhale.R

```
## pilot whale example from Isojunno et al (2017; https://doi.org/10.1002/ecs2.2044)
# requires momentuHMM >=1.5.2
library(momentuHMM)
library(TMB)
compile("pilotWhale_re.cpp")
source("whaleFunctions.R")

## load and prepare data 
load("pilotData.RData")
pilotData <- momentuHMM::prepData(pilotData,coordNames = NULL)

## data summary
summary(pilotData,dataNames=names(pilotData)[-c(1,13:15)])

nbStates <- 4 # number of states
stateNames <- c("exploratory","foraging","crowded","directed")
nbAnimals <- length(unique(pilotData$ID))

## 11 data streams
dist <- list(dive.dur = "weibull",
             dive.depth = "gamma",
             GR.speed2 = "gamma",
             dive.pitchvar2 = "beta",
             breath.headchange = "vm",
             GR.size = "pois",
             GR.tight = "bern",
             dive.CS.pres = "bern",
             dive.SS.pres = "bern",
             presurf = "bern",
             postsurf = "bern")

## initial values
Par0 <- list(dive.dur = c(1.9, 2.72, 1.64, 4.21, 1.3, 8.14, 1.53, 0.79),
             dive.depth = c(10.23, 315.91, 10.74, 5.51, 4.93, 233.12, 6.32, 1.91),
             GR.speed2 = c(1.15, 1.32, 1.36, 1.57, 0.66, 0.51, 0.77, 0.76),
             dive.pitchvar2 = c(2.21, 2.88, 1.82, 3.18, 17.94, 6.04, 16.06, 55.36),
             breath.headchange = c(3.07, 5.65, 2.64, 18.02),
             GR.size = c(6, 7.39, 20.39, 9.52),
             GR.tight = c(0.89, 0.66, 0.76, 0.81),
             dive.CS.pres = c(0.76, 0.99, 0.41, 0.47),
             dive.SS.pres = c(0.72, 0.98, 0.39, 0.41),
             presurf = c(0.76, 0.99, 0.71, 0.71),
             postsurf = c(0.81, 0.96, 0.72, 0.66))

beta0 <- matrix(c(-2.38, -3.86, -1.22, 0.21, -1.6, -0.47, -3.56, -4.15, -2.29, -1.17, -3.05, -2.54),nrow=1)


## fit model with single mixture on TPM
fitmix1 <- momentuHMM::fitHMM(pilotData,nbStates=nbStates,dist=dist,Par0=Par0,beta0=beta0,stationary=TRUE,mixtures=1,stateNames=stateNames,nlmPar=list(print.level=2))

## fit covariate model 
Par0_cov <- momentuHMM::getPar0(fitmix1,formula=~ind.size2 + ind.calf)
fitcov <- momentuHMM::fitHMM(pilotData,nbStates=nbStates,dist=dist,formula=~ind.size2 + ind.calf,Par0=Par0_cov$Par,beta0=Par0_cov$beta,stationary=TRUE,mixtures=1,stateNames=stateNames,nlmPar=list(print.level=2))

## fit model with 2 mixtures on TPM
Par0_mix2 <- momentuHMM::getPar0(fitmix1,mixtures=2)
Par0_mix2$beta$beta[1,] <- c(-2.26, -3.93, -0.58, 0.03, -2.25, -0.26, -3.38, -4.79, -2.82, -1.06, -3.3, -3.43)
Par0_mix2$beta$beta[2,] <- c(-2.51, -3.32, -2.63, 0.03, -1.26, -0.12, -96.8, -3.62, -1.75, -1.76, -2.14, -1.38)
Par0_mix2$beta$pi <- c(0.73, 0.27)
fitmix2 <- momentuHMM::fitHMM(pilotData,nbStates=nbStates,dist=dist,Par0=Par0_mix2$Par,beta0=Par0_mix2$beta,stationary=TRUE,mixtures=2,stateNames=stateNames,nlmPar=list(print.level=2))

## fit model with 3 mixtures on TPM
Par0_mix3 <- momentuHMM::getPar0(fitmix2,mixtures=3)
Par0_mix3$beta$beta[1,] <- c(-2.15, -4.31, -1.09, 0.28, -1.88, -0.3, -3.5, -4.71, -3.11, -0.68, -2.49, -2.6)
Par0_mix3$beta$beta[2,] <- c(-2.5, -2.47, 0.63, -17.22, -13.18, 0.59, -3.92, -13.96, -2.27, -1.25, -3.57, -3.75)
Par0_mix3$beta$beta[3,] <- c(-2.71, -3.48, -3.01, -0.35, -1.12, -0.1, -96.8, -2.98, -1.53, -2.29, -2.07, -1.55)
Par0_mix3$beta$pi <- c(0.4, 0.4, 0.2)
fitmix3 <- momentuHMM::fitHMM(pilotData,nbStates=nbStates,dist=dist,Par0=Par0_mix3$Par,beta0=Par0_mix3$beta,stationary=TRUE,mixtures=3,stateNames=stateNames,nlmPar=list(print.level=2))

## fit model with 4 mixtures on TPM
Par0_mix4 <- momentuHMM::getPar0(fitmix3,mixtures=4)
Par0_mix4$beta$beta[1,] <- c(-2.28, -4.09, -1.21, 0.05, -1.74, -0.26, -3.32, -5.43, -2.99, -0.76, -2.31, -2.25)
Par0_mix4$beta$beta[2,] <- c(-2.47, -2.45, 0.68, -17.22, -13.18, 0.59, -3.97, -13.97, -2.27, -1.28, -3.57, -3.75)
Par0_mix4$beta$beta[3,] <- c(-2.73, -3.47, -3, -0.37, -1.14, -0.09, -96.8, -2.97, -1.53, -2.32, -2.07, -1.54)
Par0_mix4$beta$beta[4,] <- c(-1.65, -25.12, -0.36, 2.11, -13.24, -16.48, -2.9, -3.45, -3.56, -0.39, -3.71, -31.97)
Par0_mix4$beta$pi <- c(0.33, 0.4, 0.2, 0.07)
fitmix4 <- momentuHMM::fitHMM(pilotData,nbStates=nbStates,dist=dist,Par0=Par0_mix4$Par,beta0=Par0_mix4$beta,stationary=TRUE,mixtures=4,stateNames=stateNames,nlmPar=list(print.level=2))

## fixed effects model
Par0_fix <- momentuHMM::getPar0(fitmix4,formula=~0+ID,mixtures=1)
fitfix <- momentuHMM::fitHMM(pilotData,nbStates=nbStates,dist=dist,formula=~0+ID,stationary=TRUE,Par0=Par0_fix$Par,stateNames=stateNames,nlmPar=list(print.level=2))

## BW approximate continuous random effects model (fails)
fitBW <- momentuHMM::randomEffects(fitfix)

## fit continuous random effects model using TMB
Par0_tmb<- list(l_ddurshape=log(Par0$dive.dur[1:nbStates]),
              l_ddurscale=log(Par0$dive.dur[nbStates+1:nbStates]),
              l_ddepmu=log(Par0$dive.depth[1:nbStates]),
              l_ddepsd=log(Par0$dive.depth[nbStates+1:nbStates]),
              l_GRspeedmu=log(Par0$GR.speed2[1:nbStates]),
              l_GRspeedsd=log(Par0$GR.speed2[nbStates+1:nbStates]),
              l_dpitchshape1=log(Par0$dive.pitchvar2[1:nbStates]),
              l_dpitchshape2=log(Par0$dive.pitchvar2[nbStates+1:nbStates]),
              l_brkappa=log(Par0$breath.headchange),
              l_GRsizerate=log(Par0$GR.size),
              l_GRtightprob=stats::qlogis(Par0$GR.tight),
              l_diveCSprob=stats::qlogis(Par0$dive.CS.pres),
              l_diveSSprob=stats::qlogis(Par0$dive.SS.pres),
              l_presurfprob=stats::qlogis(Par0$presurf),
              l_postsurfprob=stats::qlogis(Par0$postsurf),
              l_gamma=matrix(c(-2.41, -4.08, -0.72,  0.03, -1.74, -0.18, -3.68, -4.15, -2.16, -0.97, -3.24, -2.69),1,nbStates*(nbStates-1)),
              l_sigma=c(-0.75,    0.14,    0.16,   -0.58,   -1.41,   -3.54,   -3.91,   -1.54,   -0.07,   -0.60,    0.14,   -0.23),
              epsilon=matrix(0,nbAnimals,nbStates*(nbStates-1)))

covIndex <- c(1,cumsum(table(pilotData$ID))[1:(nbAnimals-1)]+1) # first observation for each individual

dyn.load(dynlib("pilotWhale_re"))
fitTMB <- whaleTMB(as.matrix(pilotData[,-c(1,13:15)]), nbStates, nrow(pilotData), covIndex-1, Par0_tmb, "pilotWhale_re", "epsilon") 
dyn.unload(dynlib("pilotWhale_re"))

## AICc
aicResults <- AIC(fitmix1,fitcov,fitmix2,fitmix3,fitmix4,fitfix,momentuHMM:::momentuHMM(list(mle=fitTMB$mle, mod=fitTMB$mod ,data=fitfix$data, conditions=fitfix$conditions, stateNames=stateNames)),n=nrow(pilotData))
aicWeights <- AICweights(fitmix1,fitcov,fitmix2,fitmix3,fitmix4,fitfix,momentuHMM:::momentuHMM(list(mle=fitTMB$mle, mod=fitTMB$mod ,data=fitfix$data, conditions=fitfix$conditions, stateNames=stateNames)),n=nrow(pilotData))

## BIC
bicResults <- AIC(fitmix1,fitcov,fitmix2,fitmix3,fitmix4,fitfix,momentuHMM:::momentuHMM(list(mle=fitTMB$mle, mod=fitTMB$mod ,data=fitfix$data, conditions=fitfix$conditions, stateNames=stateNames)),k=log(nrow(pilotData)))
bicWeights <- AICweights(fitmix1,fitcov,fitmix2,fitmix3,fitmix4,fitfix,momentuHMM:::momentuHMM(list(mle=fitTMB$mle, mod=fitTMB$mod ,data=fitfix$data, conditions=fitfix$conditions, stateNames=stateNames)),k=log(nrow(pilotData)))

# likelihood ratio test
lrt(fitmix1,fitTMB)

# plot TMB random effect variances
plot(1:(nbStates*(nbStates-1)),unlist(lapply(fitTMB$varcomp,function(x) x$beta$sigma$est)),
     ylim=c(min(unlist(lapply(fitTMB$varcomp,function(x) x$beta$sigma$lower))),10),#max(unlist(lapply(fitTMB$varcomp,function(x) x$beta$sigma$upper)))), 
     xlab="state transition",ylab=expression(sigma),xaxt="n",pch=20)
axis(1, at=1:length(colnames(fitTMB$mle$beta)),labels=colnames(fitTMB$mle$beta))
abline(0,0,h=0,lty=2)
graphics::arrows(1:(nbStates*nbStates-1),unlist(lapply(fitTMB$varcomp,function(x) x$beta$sigma$lower)),1:(nbStates*nbStates-1),unlist(lapply(fitTMB$varcomp,function(x) x$beta$sigma$upper)),length=0.025, angle=90, code=3, col=gray(.5))

# plot TMB state transition probabilties
par(mar=c(4,4,1.5,2)-c(0,0,1.5,1))
par(mfrow=c(nbStates,nbStates))
for(i in 1:nbStates){
  for(j in 1:nbStates){
    #par(mar=c(1,1,1,1))
    plot(factor(1:nbAnimals),fitTMB$trProbs$est[i,j,],
         ylim=c(0,1),
         xlab=ifelse(i==nbStates,"Individual",NA),ylab=paste0(i," -> ",j),pch=20,bty="n")
    abline(h=0.5,lty=2)
    graphics::arrows(1:nbAnimals,fitTMB$trProbs$lower[i,j,],
                     1:nbAnimals,fitTMB$trProbs$upper[i,j,],length=0.025, angle=90, code=3, col=gray(.5))
  }
}

# small differences when it comes to Viterbi-decoded state assignment
sum(fitTMB$states==momentuHMM::viterbi(fitmix1))/nrow(pilotData) # 94% agreement between tmb and basic model used by Isojunno et al
sum(momentuHMM::viterbi(fitmix3)==momentuHMM::viterbi(fitmix1))/nrow(pilotData) # 91% agreement between K=3 mixtures and basic model used by Isojunno et al
sum(fitTMB$states==momentuHMM::viterbi(fitmix3))/nrow(pilotData) # 95% agreement between tmb and K=3 mixtures
```
