## Supplementary material for "Worth the effort? A practical examination of random effects in hidden Markov models for animal telemetry data": Code: pilotWhale_re.cpp

```
#include <TMB.hpp>
#include <cmath>

using namespace density;
using std::sqrt;

template <class Type>
int which_max(vector<Type> x){
  int first = 0;
  int last = x.size()-1;
  if (first==last) return last;
  int largest = first;
  
  while (++first<last+1)
    if (x(largest)<x(first))    
      largest=first;
    return largest;
}


/* Numerically stable forward algorithm based on http://bozeman.genome.washington.edu/compbio/mbt599_2006/hmm_scaling_revised.pdf */
template<class Type>
Type forward_alg(array<Type> delta, array<Type> trMat, matrix<Type> lnProbs, int nbSteps, vector<int> aInd) {
  int nbStates = trMat.cols();
  Type logalpha;
  matrix<Type> ltrMat(nbStates,nbStates);
  vector<Type> ldeltaG(nbStates);
  vector<Type> lalpha(nbStates);
  vector<Type> lnewalpha(nbStates);
  Type jnll = 0.0;
  Type sumalpha;
  int nbAnimals = aInd.size();
  int animal = 0;
  for(int t=0; t < nbSteps; t++){
    if(animal<nbAnimals && t==aInd(animal)){
      sumalpha = -INFINITY;
      for(int j=0; j < nbStates; j++){
        ldeltaG(j) = -INFINITY;
        for(int i=0; i < nbStates; i++){
          ltrMat(i,j) = log(trMat(animal,i,j));
          ldeltaG(j) = logspace_add(ldeltaG(j),log(delta(animal,0,i))+ltrMat(i,j));
        }
        lalpha(j) = ldeltaG(j)+lnProbs(t,j);
        sumalpha  = logspace_add(sumalpha,lalpha(j));
      }
      jnll -= sumalpha;
      lalpha -= sumalpha;
      animal++;
    } else {
      sumalpha = -INFINITY;
      for(int j=0; j < nbStates; j++){
        logalpha = -INFINITY;
        for(int i=0; i < nbStates; i++){
          logalpha = logspace_add(logalpha,lalpha(i)+ltrMat(i,j));
        }
        lnewalpha(j) = logalpha + lnProbs(t,j);
        sumalpha  = logspace_add(sumalpha,lnewalpha(j));
      }
      jnll -= sumalpha;
      lalpha = lnewalpha - sumalpha;
    }
  }
  return jnll;
}

/* Numerically stable viterbi based on Rabiner 1989 IEEE 77(2):257-286 */
template<class Type>
vector<int> viterbi(array<Type> delta, array<Type> trMat, matrix<Type> lnProbs, int nbSteps, vector<int> aInd) {
  int nbStates = trMat.cols();
  matrix<Type> phi(nbSteps,nbStates);
  matrix<int> psi(nbSteps,nbStates);
  vector<Type> tmpphi(nbStates);
  vector<int> states(nbSteps);
  states.setOnes();
  matrix<Type> ltrMat(nbStates,nbStates);
  vector<Type> ldeltaG(nbStates);
  int nbAnimals = aInd.size();
  int animal = 0;
  for(int t=0; t < nbSteps; t++){
    if(animal<nbAnimals && t==aInd(animal)){
      for(int j=0; j < nbStates; j++){
        ldeltaG(j) = -INFINITY;
        for(int i=0; i < nbStates; i++){
          ltrMat(i,j) = log(trMat(animal,i,j));
          ldeltaG(j) = logspace_add(ldeltaG(j),log(delta(animal,0,i))+ltrMat(i,j));
        }
        psi(t,j) = 0;
        phi(t,j) = ldeltaG(j)+lnProbs(t,j);
      }
      animal++;
    } else {
      for(int j=0; j < nbStates; j++){
        for(int i=0; i < nbStates; i++){
          tmpphi(i) = phi(t-1,i)+ltrMat(i,j);
        }
        psi(t,j) = which_max(tmpphi);
        phi(t,j) = tmpphi(psi(t,j))+lnProbs(t,j);
      }
    }
  }
  
  states(nbSteps-1) += which_max(vector<Type>(phi.row(nbSteps-1)));
  animal = nbAnimals-1;
  for(int t=(nbSteps-2); t >= 0 ; t--){
    if(animal>0 && t==(aInd(animal)-1)){
      states(t) += which_max(vector<Type>(phi.row(t)));
      animal--;
    } else {
      states(t) += psi(t+1,states(t+1)-1);
    }
  }
  return states;
}

/* Numerically stable forward log-probabilities based on http://bozeman.genome.washington.edu/compbio/mbt599_2006/hmm_scaling_revised.pdf */
template<class Type>
matrix<Type> logAlpha(array<Type> delta, array<Type> trMat, matrix<Type> lnProbs, int nbSteps, vector<int> aInd) {
  int nbStates = trMat.cols();
  Type logalpha;
  matrix<Type> elnalpha(nbSteps,nbStates);
  matrix<Type> ltrMat(nbStates,nbStates);
  vector<Type> ldeltaG(nbStates);
  int nbAnimals = aInd.size();
  int animal = 0;
  for(int t=0; t < nbSteps; t++){
    if(animal<nbAnimals && t==aInd(animal)){
      for(int j=0; j < nbStates; j++){
        ldeltaG(j) = -INFINITY;
        for(int i=0; i < nbStates; i++){
          ltrMat(i,j) = log(trMat(animal,i,j));
          ldeltaG(j) = logspace_add(ldeltaG(j),log(delta(animal,0,i))+ltrMat(i,j));
        }
        elnalpha(t,j) = ldeltaG(j)+lnProbs(t,j);
      }
      animal++;
    } else {
      for(int j=0; j < nbStates; j++){
        logalpha = -INFINITY;
        for(int i=0; i < nbStates; i++){
          logalpha = logspace_add(logalpha,elnalpha(t-1,i)+ltrMat(i,j));
        }
        elnalpha(t,j) = logalpha+lnProbs(t,j);
      }
    }
  }
  return elnalpha;
}

/* Numerically stable backward log-probabilities based on http://bozeman.genome.washington.edu/compbio/mbt599_2006/hmm_scaling_revised.pdf */
template<class Type>
matrix<Type> logBeta(array<Type> delta, array<Type> trMat, matrix<Type> lnProbs, int nbSteps, vector<int> aInd) {
  int nbStates = trMat.cols();
  Type logbeta;
  matrix<Type> elnbeta(nbSteps,nbStates);
  matrix<Type> ltrMat(nbStates,nbStates);
  int nbAnimals = aInd.size();
  
  int animal = nbAnimals-1;
  for(int j=0; j < nbStates; j++){
    elnbeta(nbSteps-1,j) = Type(0.0);
    for(int i=0; i < nbStates; i++){
      ltrMat(i,j) = log(trMat(animal,i,j));
    }
  }
  for(int t=(nbSteps-2); t >= 0 ; t--){
    if(animal>0 && t==(aInd(animal)-1)){
      animal--;
      for(int j=0; j < nbStates; j++){
        elnbeta(t,j) = Type(0.0);
        for(int i=0; i < nbStates; i++){
          ltrMat(i,j) = log(trMat(animal,i,j));
        }
      }
    } else {
      for(int i=0; i < nbStates; i++){
        logbeta = -INFINITY;
        for(int j=0; j < nbStates; j++){
          logbeta = logspace_add(logbeta,ltrMat(i,j)+lnProbs(t+1,j)+elnbeta(t+1,j));
        }
        elnbeta(t,i) = logbeta;
      }
    }
  }
  return elnbeta;
}

/* Numerically stable state probabilities based on http://bozeman.genome.washington.edu/compbio/mbt599_2006/hmm_scaling_revised.pdf */
template<class Type>
matrix<Type> stProbs(matrix<Type> elnalpha, matrix<Type> elnbeta, int nbSteps) {
  int nbStates = elnalpha.cols();
  Type normalizer;
  matrix<Type> gamma(nbSteps,nbStates);
  
  for(int t=0; t < nbSteps; t++){
    normalizer = -INFINITY;
    for(int i=0; i < nbStates; i++){
      gamma(t,i) = elnalpha(t,i)+elnbeta(t,i);
      normalizer = logspace_add(normalizer,gamma(t,i));
    }
    for(int i=0; i < nbStates; i++){
      gamma(t,i) = exp(gamma(t,i)-normalizer);
    }
  }
  return gamma;
}

template<class Type>
Type objective_function<Type>::operator() ()
{
  // DATA
  DATA_MATRIX(data);	            //  observations
  DATA_INTEGER(nbStates);        // number of states
  DATA_INTEGER(nbSteps);          // number of observations
  DATA_IVECTOR(aInd);          // indicator for first observation of each individual
  
  // OBSERVATION PARAMETERS
  PARAMETER_VECTOR(l_ddurshape); 				// dive duration weibull shape (log scale)
  PARAMETER_VECTOR(l_ddurscale);     	// dive duration weibull scale (log scale)
  PARAMETER_VECTOR(l_ddepmu); 				// dive depth gamma mean (log scale)
  PARAMETER_VECTOR(l_ddepsd);     	// dive depth gamma sd (log scale)
  PARAMETER_VECTOR(l_GRspeedmu); 				// GR.speed gamma mean (log scale)
  PARAMETER_VECTOR(l_GRspeedsd);     	// GR.speed gamma sd (log scale)
  PARAMETER_VECTOR(l_dpitchshape1); 				// dive pitch beta shape1 (log scale)
  PARAMETER_VECTOR(l_dpitchshape2);     	// dive pitch beta shape2 (log scale)
  PARAMETER_VECTOR(l_brkappa);            // breath vm kappa (log scale)
  PARAMETER_VECTOR(l_GRsizerate);         // GR.size poisson rate (log scale)
  PARAMETER_VECTOR(l_GRtightprob);        // GR.tight bern prob (logit scale)
  PARAMETER_VECTOR(l_diveCSprob);        // dive.CS bern prob (logit scale)
  PARAMETER_VECTOR(l_diveSSprob);        // dive.SS bern prob (logit scale)
  PARAMETER_VECTOR(l_presurfprob);        // presurf bern prob (logit scale)
  PARAMETER_VECTOR(l_postsurfprob);        // postsurf bern prob (logit scale)
  
  // t.p.m. random effect parameters
  PARAMETER_MATRIX(l_gamma); // t.p.m. (mlogit scale) - should be 1 x (nbStates-1) * nbStates
  PARAMETER_VECTOR(l_sigma); // sd for random effects (log scale)
  
  // random effects
  PARAMETER_MATRIX(epsilon);
  
  // Transform parameters
  vector<Type> ddurshape(nbStates);
  vector<Type> ddurscale(nbStates);
  vector<Type> ddepshape(nbStates);
  vector<Type> ddepscale(nbStates);
  vector<Type> GRspeedshape(nbStates);
  vector<Type> GRspeedscale(nbStates);
  vector<Type> dpitchshape1(nbStates);
  vector<Type> dpitchshape2(nbStates);
  vector<Type> brkappa(nbStates);
  vector<Type> brmu(nbStates);
  vector<Type> GRsizerate(nbStates);
  vector<Type> GRtightprob(nbStates);
  vector<Type> diveCSprob(nbStates);
  vector<Type> diveSSprob(nbStates);
  vector<Type> presurfprob(nbStates);
  vector<Type> postsurfprob(nbStates);
  vector<Type> mu(nbStates);
  vector<Type> sd(nbStates);
  for(int i=0; i < nbStates; i++){
    ddurshape(i) = exp(l_ddurshape(i));
    ddurscale(i) = exp(l_ddurscale(i));
    mu(i) = exp(l_ddepmu(i));
    sd(i) = exp(l_ddepsd(i));
    ddepshape(i) = mu(i) * mu(i) / (sd(i) * sd(i));
    ddepscale(i) = sd(i) * sd(i) / mu(i);
    mu(i) = exp(l_GRspeedmu(i));
    sd(i) = exp(l_GRspeedsd(i));
    GRspeedshape(i) = mu(i) * mu(i) / (sd(i) * sd(i));
    GRspeedscale(i) = sd(i) * sd(i) / mu(i);
    dpitchshape1(i) = exp(l_dpitchshape1(i));
    dpitchshape2(i) = exp(l_dpitchshape2(i));
    brkappa(i) = exp(l_brkappa(i));
    brmu(i) = Type(0.0);
    GRsizerate(i) = exp(l_GRsizerate(i));
    GRtightprob(i) = exp(l_GRtightprob(i))/(Type(1.0)+exp(l_GRtightprob(i)));
    diveCSprob(i) = exp(l_diveCSprob(i))/(Type(1.0)+exp(l_diveCSprob(i)));
    diveSSprob(i) = exp(l_diveSSprob(i))/(Type(1.0)+exp(l_diveSSprob(i)));
    presurfprob(i) = exp(l_presurfprob(i))/(Type(1.0)+exp(l_presurfprob(i)));
    postsurfprob(i) = exp(l_postsurfprob(i))/(Type(1.0)+exp(l_postsurfprob(i)));
  }
  
  int nbAnimals = aInd.size();
  
  /* Define likelihood */
  //parallel_accumulator<Type> nll(this);
  Type nll=0.0;
  
  vector<Type> sigma(nbStates*(nbStates-1));
  
  array<Type> delta(nbAnimals, 1, nbStates); 
  matrix<Type> idelta(1, nbStates);
  matrix<Type> I = matrix<Type>::Identity(nbStates, nbStates);
  matrix<Type> tpminv = I; 
  
  // tpm
  array<Type> trMat(nbAnimals,nbStates,nbStates);
  matrix<Type> iMat(nbStates,nbStates);
  Type denom;
  int cpt;
  for(int s=0;s<nbAnimals;s++){
    cpt = 0;
    for(int i=0;i<nbStates;i++){
      denom = Type(0.0);
      for(int j=0;j<nbStates;j++){
        if(i==j) {
          trMat(s,i,j) = Type(1.0);
          cpt++;
        } else {
          sigma(i*nbStates+j-cpt) = exp(l_sigma(i*nbStates+j-cpt));
          trMat(s,i,j) = exp(epsilon(s,i*nbStates+j-cpt));
          nll -= dnorm(epsilon(s,i*nbStates+j-cpt),l_gamma(0,i*nbStates+j-cpt),sigma(i*nbStates+j-cpt),true);
        }
        denom += trMat(s,i,j);
      }
      for(int j=0;j<nbStates;j++){
        trMat(s,i,j) /= denom;
        iMat(i,j) = trMat(s,i,j);
      }
    }
    // Compute stationary distribution
    tpminv = I; 
    tpminv -= iMat; 
    tpminv = (tpminv.array() + 1).matrix(); 
    matrix<Type> ivec(1, nbStates); for (int i = 0; i < nbStates; ++i) ivec(0, i) = 1;
    // if tpm is ill-conditioned then just use uniform initial distribution 
    try {
      tpminv = tpminv.inverse();
      idelta = ivec * tpminv;
    } catch(...) {
      for (int i = 0; i < nbStates; ++i) idelta(0, i) = 1.0 / nbStates; 
    }
    for(int i=0;i<nbStates; ++i){
      delta(s,0,i) = idelta(0,i);
    }
  }
  
  matrix<Type> allProbs(nbSteps,nbStates);
  allProbs.setZero();
  
  // observations
  
  for(int state = 0; state < nbStates; state++){
    for(int t=0; t < nbSteps; t++){
      if(!isnan(data(t,0))) allProbs(t,state) += dweibull(data(t,0),ddurshape(state),ddurscale(state),1);
      if(!isnan(data(t,1))) allProbs(t,state) += dgamma(data(t,1),ddepshape(state),ddepscale(state),1);
      if(!isnan(data(t,2))) allProbs(t,state) += dgamma(data(t,2),GRspeedshape(state),GRspeedscale(state),1);
      if(!isnan(data(t,3))) allProbs(t,state) += dbeta(data(t,3),dpitchshape1(state),dpitchshape2(state),1);
      if(!isnan(data(t,4))) allProbs(t,state) +=  brkappa(state) *cos (data(t,4)-brmu(state)) - log (2* M_PI ) - log ( besselI(brkappa(state),Type(0.0)) ) ;
      if(!isnan(data(t,5))) allProbs(t,state) += dpois(data(t,5),GRsizerate(state),1);
      if(!isnan(data(t,6))) allProbs(t,state) += dbinom(data(t,6),Type(1.0),GRtightprob(state),1);
      if(!isnan(data(t,7))) allProbs(t,state) += dbinom(data(t,7),Type(1.0),diveCSprob(state),1);
      if(!isnan(data(t,8))) allProbs(t,state) += dbinom(data(t,8),Type(1.0),diveSSprob(state),1);
      if(!isnan(data(t,9))) allProbs(t,state) += dbinom(data(t,9),Type(1.0),presurfprob(state),1);
      if(!isnan(data(t,10))) allProbs(t,state) += dbinom(data(t,10),Type(1.0),postsurfprob(state),1);
    }
  }
  
  REPORT(allProbs);
  
  // forward algorithm
  Type hmmll = forward_alg(delta, trMat, allProbs, nbSteps, aInd);
  REPORT(hmmll);
  nll += hmmll;
  
  ADREPORT(ddurshape);
  ADREPORT(ddurscale);
  ADREPORT(ddepshape);
  ADREPORT(ddepscale);
  ADREPORT(GRspeedshape);
  ADREPORT(GRspeedscale);
  ADREPORT(dpitchshape1);
  ADREPORT(dpitchshape2);
  ADREPORT(brkappa);
  ADREPORT(GRsizerate);
  ADREPORT(GRtightprob);
  ADREPORT(diveCSprob);
  ADREPORT(diveSSprob);
  ADREPORT(presurfprob);
  ADREPORT(postsurfprob);
  
  if(nbStates>1){
    vector<int> states = viterbi(delta, trMat, allProbs, nbSteps, aInd);
    REPORT(states);
    matrix<Type> lalpha = logAlpha(delta, trMat, allProbs, nbSteps, aInd);
    //REPORT(lalpha);
    matrix<Type> lbeta = logBeta(delta, trMat, allProbs, nbSteps, aInd);
    //REPORT(lbeta);
    matrix<Type> stateProbs = stProbs(lalpha, lbeta, nbSteps); 
    REPORT(stateProbs);
    ADREPORT(delta);
    ADREPORT(trMat);
    ADREPORT(sigma);
  }
  
  return nll;
}
```
