## Supplementary material for "Worth the effort? A practical examination of random effects in hidden Markov models for animal telemetry data": Code: simFunctions.R

```
fitTMB <- function(data, covs, nbStates, nbSteps, aInd, covInd, par, ran = c("epsilon")) {
  dat <- list(data = data, cov = covs, nbStates=nbStates, nbSteps=nbSteps, aInd=aInd)
  # create function for automatic differentiation 
  # dat must contain all DATA required by hmm_tmb.cpp 
  # par must contain all PARAMETERS required by hmm_tmb.cpp 
  
  if(!covInd) map <- list(mu_1=factor(rep(NA,nbStates*(nbStates-1))))
  else map <- list()
  startTime <- proc.time()
  obj <- MakeADFun(data = dat, 
                   parameters = par, 
                   map = map,
                   random = ran,
                   DLL = "HMM_recov", 
                   hessian = TRUE,
                   silent = TRUE)
  mod <- do.call("optim", obj)
  sdr <- sdreport(obj,hessian.fixed=mod$hessian)
  endTime <- proc.time()-startTime
  rep <- summary(sdr, "report")
  
  nbAnimals <- length(aInd)
  
  trMat <- parSum(rep,which(rownames(rep)=="trMat"),nbAnimals,nbStates^2,FALSE,dimnames=list(paste0("ID",1:nbAnimals),c("1 -> 1","2 -> 1","1 -> 2","2 -> 2"))) 
  trProbs <- lapply(trMat,function(x) array(t(x),dim=c(nbStates,nbStates,nbAnimals)))
  trProbs <- lapply(trProbs,function(x){rownames(x)<-colnames(x)<-c("state 1","state 2");x})
  trProbs$lower <- 1/(1+exp(-(log(trProbs$est/(1-trProbs$est))-qnorm(0.975)*(1/(trProbs$est-trProbs$est^2))*trProbs$se)))
  trProbs$upper <- 1/(1+exp(-(log(trProbs$est/(1-trProbs$est))+qnorm(0.975)*(1/(trProbs$est-trProbs$est^2))*trProbs$se)))
  
  fxd <- summary(sdr,"fixed")
  
  CIbeta <- list()
  if(!is.null(ran)){
    ran <- summary(sdr, "random")
    CIbeta$beta <- parSum(ran,1:nrow(ran),nbAnimals,nbStates*(nbStates-1))
    betamle <- matrix(ran[,1],nbAnimals,nbStates*(nbStates-1),dimnames=list(paste0("ID",1:nbAnimals),c("1 -> 2","2 -> 1")))
  } else {
    if(!any(rownames(fxd) %in% "epsilon")) {
      CIbeta$beta <- parSum(fxd,which(rownames(fxd) %in% "mu_0"),nbAnimals,nbStates*(nbStates-1),byrow=TRUE,dimnames=list(paste0("ID",1:nbAnimals),c("1 -> 2","2 -> 1")))
      betamle <- matrix(mod$par[2*nbStates+1:(nbStates*(nbStates-1))],nbAnimals,nbStates*(nbStates-1),byrow=TRUE)
    } else {
      CIbeta$beta <- parSum(fxd,which(rownames(fxd) %in% "epsilon"),nbAnimals,nbStates*(nbStates-1))
      betamle <- matrix(fxd[which(rownames(fxd) %in% "epsilon"),1],nbAnimals,nbStates*(nbStates-1),dimnames=list(paste0("ID",1:nbAnimals),c("1 -> 2","2 -> 1")))
    }
  }
  
  
  CIbeta$step <- parSum(fxd,1:(nbStates*2),1,(nbStates*2),TRUE,list(NULL,c("mean_1:(Intercept)","mean_2:(Intercept)","sd_1:(Intercept)","sd_2:(Intercept)")))
  CIbeta$delta <- parSum(fxd,which(rownames(fxd) %in% "l_delta"),1,nbStates-1,TRUE,list("(Intercept)",paste0("state ",2:nbStates)))
  CIbeta$mu_0 <- parSum(fxd,which(rownames(fxd) %in% "mu_0"),1,nbStates*(nbStates-1),TRUE,list("(Intercept)",c("1 -> 2","2 -> 1")))
  if(covInd) CIbeta$mu_1 <- parSum(fxd,which(rownames(fxd) %in% "mu_1"),1,nbStates*(nbStates-1),TRUE,list("cov",c("1 -> 2","2 -> 1")))
  
  CIreal <- list()
  CIreal$step <- parSum(rep,1:(nbStates*2),2,nbStates,TRUE,list(c("mean","sd"),c("state 1","state 2")))
  CIreal$gamma <- lapply(trProbs,function(x) x[,,1])
  CIreal$delta <- parSum(rep,which(rownames(rep)=="delta"),nbAnimals,2,TRUE,list(paste0("ID",1:nbAnimals),c("state 1","state 2")))
  
  varcomp <- vector('list',nbStates*(nbStates-1))
  for(i in 1:length(varcomp)){
    varcomp[[i]] <- list()
    varcomp[[i]]$sigma <- parSum(rep,which(rownames(rep)=="sigma")[i],1,1)
    cn<-exp(qnorm(0.975)*sqrt(log(1+(varcomp[[i]]$sigma$se/(varcomp[[i]]$sigma$est))^2)))
    varcomp[[i]]$sigma$lower <- varcomp[[i]]$sigma$est/cn
    varcomp[[i]]$sigma$upper <- varcomp[[i]]$sigma$est*cn
    varcomp[[i]]$mu_0 <- lapply(CIbeta$mu_0,function(x) x[i])
    if(covInd) varcomp[[i]]$mu_1 <- lapply(CIbeta$mu_1,function(x) x[i])
  }
  mle <- list(step=matrix(rep[1:4,1],2,2,byrow=TRUE,dimnames=list(c("mean","sd"),c("state 1","state 2"))),
              beta=betamle,
              delta=matrix(rep[5:6,1],nbAnimals,2,byrow=TRUE,dimnames = list(paste0("ID:",1:nbAnimals),c("state 1","state 2"))))
  report <- obj$report(obj$env$last.par.best)
  nparm <- length(mod[["par"]])
  return(list(mod=list(minimum = mod$value, estimate = mod$par, gradient = obj$gr(mod$par), hessian=mod$hessian, code=mod$convergence, iterations=mod$counts, elapsedTime=endTime[3], wpar=mod$par, Sigma=MASS::ginv(mod$hessian), message=mod$message),mle=mle,CIreal=CIreal,CIbeta=CIbeta,trProbs=trProbs,states=report$states,stateProbs=report$stateProbs,varcomp=varcomp,aic =  2 * nparm + 2 * mod[["value"]],aicc =  2 * nparm + 2 * mod[["value"]]+2*nparm*(nparm+1)/(length(data)-nparm-1),bic =  log(length(data)) * nparm + 2 * mod[["value"]]))
}

parSum <- function(par,index,nr,nc,byrow=FALSE,dimnames=NULL){
  out <- list(est=matrix(par[index,1],nr,nc,byrow=byrow,dimnames=dimnames),se=matrix(par[index,2],nr,nc,byrow=byrow,dimnames=dimnames))
  out$lower <- out$est - qnorm(0.975)*out$se
  out$upper <- out$est + qnorm(0.975)*out$se
  return(out)
}

get_indProbs <- function(estimate,mod,covs=matrix(1,1,1),nbStates,i,j,betaRef,betaCons,workBounds=NULL,mixtures=1){
  
  nbCovs <- ncol(covs)-1
  
  pibeta <- estimate[4+1:(nbStates*(nbStates-1)*mixtures*(nbCovs+1)+(mixtures-1))]
  if(mixtures>1) {
    pie <- pibeta[length(pibeta)-(mixtures-2):0]
    pie <- momentuHMM:::mlogit(matrix(pie,1,mixtures-1),matrix(1,1,1),0,1,mixtures)
  } else pie <- 1
  
  
  beta <- matrix(pibeta[1:(nbStates*(nbStates-1)*mixtures*(nbCovs+1))],mixtures*(nbCovs+1),nbStates*(nbStates-1))
  
  pInd <- which(mapply(function(x) isTRUE(all.equal(x,0)),pie))
  if(length(pInd)){
    pie[pInd] <- 1.e-100
    pie[-pInd] <- pie[-pInd] - (1.e-100*length(pInd))/(mixtures-length(pInd))
  }
  
  par <- list(pi=matrix(1,1,1))
  par$step <- matrix(exp(estimate[1:4]),4,nrow(mod$data))
  gamma <- mixProbs <- lnum <- la <- numeric(mixtures)
  for(mix in 1:mixtures){
    par$beta <- beta[(mix-1)*(nbCovs+1)+1:(nbCovs+1),,drop=FALSE]
    par$delta <- momentuHMM:::mlogit(matrix(estimate[length(estimate)-mixtures+mix],1),mod$covsDelta,0,1,nbStates,1)
    la[mix] <- momentuHMM:::nLogLike_rcpp(nbStates,as.matrix(covs),mod$data,names(mod$conditions$dist),mod$conditions$dist,
                                          par,
                                          1,mod$conditions$zeroInflation,mod$conditions$oneInflation,mod$conditions$stationary,mod$conditions$knownStates,mod$conditions$betaRef,1)
    gamma[mix] <- momentuHMM:::trMatrix_rcpp(nbStates,beta[(mix-1)*(nbCovs+1)+1:(nbCovs+1),,drop=FALSE],as.matrix(covs),betaRef)[i,j,1]
    c <- max(-la[mix]+log(pie[mix]))
    lnum[mix] <- c + log(sum(exp(-la[mix]+log(pie[mix])-c)))  
  }
  c <- max(lnum)
  mixProbs <- exp(lnum - c - log(sum(exp(lnum-c))))
  mixProbs[pInd] <- 0
  return(sum(mixProbs * gamma))
}

mixTrProbs <- function(m){
  nbStates <- length(m$stateNames)
  nbAnimals <- length(unique(m$data$ID))
  est <- se <- lower <- upper <- array(NA,dim=c(nbStates,nbStates,nbAnimals))
  for(k in 1:(nbAnimals)){
    for(i in 1:nbStates){
      for(l in 1:nbStates){
        if(l==1) {
          tmp <- m
          tmp$data <- m$data[which(m$data$ID==k),]
          covs <- model.matrix(m$conditions$formula,tmp$data)
          tmp$covsDelta <- tmp$covsDelta[k,,drop=FALSE]
          tmp$covsPi <- tmp$covsPi[k,,drop=FALSE]
          tmp$conditions$knownStates <- rep(NA,nrow(tmp$data))
          est[i,l,k] <- get_indProbs(tmp$mod$estimate,mod=tmp,covs=covs,nbStates=nbStates,i=i,j=l,tmp$conditions$betaRef,tmp$conditions$betaCons,tmp$conditions$workBounds$beta,mixtures=j)
          dN<-t(tryCatch(numDeriv::grad(get_indProbs,tmp$mod$estimate,mod=tmp,covs=covs,nbStates=nbStates,i=i,j=l,betaRef=tmp$conditions$betaRef,betaCons=tmp$conditions$betaCons,workBounds=tmp$conditions$workBounds$beta,mixtures=j),error=function(e) NA))
          se[i,l,k] <- sqrt(dN %*% tmp$mod$Sigma %*% t(dN))
        } else {
          est[i,l,k] <- 1-est[i,l-1,k]
          se[i,l,k] <- se[i,l-1,k]
        }
        lower[i,l,k] <- 1/(1+exp(-(log(est[i,l,k]/(1-est[i,l,k]))-qnorm(0.975)*(1/(est[i,l,k]-est[i,l,k]^2))*se[i,l,k])))
        upper[i,l,k] <- 1/(1+exp(-(log(est[i,l,k]/(1-est[i,l,k]))+qnorm(0.975)*(1/(est[i,l,k]-est[i,l,k]^2))*se[i,l,k])))
      }
    }
  }
  return(list(est=est,se=se,lower=lower,upper=upper))
}

get_mixCovs <- function(parms,nbStates,mixtures,k){
  piInd <- 4+2*mixtures*(nbStates*(nbStates-1))+1:(mixtures-1)
  bInd <- 4+(k-1)*mixtures*2+seq(2,2*mixtures,2)
  sum(exp(c(0,parms[piInd]))/sum(exp(c(0,parms[piInd])))*parms[bInd])
  betaPi <- c(0,parms[piInd])
  sum(exp(betaPi)/sum(exp(betaPi))*parms[bInd])
}

mixCovs <- function(m){
  est <- se <- lower <- upper <- matrix(NA,1,nbStates*(nbStates-1),dimnames=dimnames(mixOut$cov[[1]]$CIreal$beta$est))
  for(k in 1:(nbStates*(nbStates-1))){
    est[,k] <- get_mixCovs(m$mod$estimate,nbStates,m$conditions$mixtures,k)
    dN<-t(tryCatch(numDeriv::grad(get_mixCovs,m$mod$estimate,nbStates=nbStates,mixtures=m$conditions$mixtures,k=k),error=function(e) NA))
    se[,k] <- sqrt(dN %*% m$mod$Sigma %*% t(dN))
    lower[,k] <- est[,k]-qnorm(0.975)*se[,k]
    upper[,k] <- est[,k]+qnorm(0.975)*se[,k]
  }
  return(list(est=est,se=se,lower=lower,upper=upper))
}

lrt <- function(null,mod){
  nbStates <- length(null$stateNames)
  LR<- -2*(-null$mod$minimum- -mod$mod$minimum)
  X2 <- 0
  q <- 0
  qprime <- nbStates*(nbStates-1)
  for(r in 0:qprime){
    X2 <- X2 + 2^(-qprime)*choose(qprime,r)*dchisq(LR,r)
    q <- q + 2^(-qprime)*choose(qprime,r)*pchisq(LR,r)
  }
  p <- 1-q # if p<0.05, reject null of no individual-level variation
  cat(ifelse(p<0.05,"reject","failure to reject")," null at 0.05 significance level: p-value = ",p,", LRT statistic = ",LR,"\n",sep="")
}

Kappa <- function(modstates,states){
  X<-matrix(0,nbStates,nbStates)
  for(i in 1:nbStates){
    for(j in 1:nbStates){
      X[i,j]<-sum(states==i & modstates==j)
    }
  }
  p_0 <- sum(diag(X))/sum(X)
  p_e <- sum(rowSums(X)*colSums(X))/(sum(X)^2)
  (p_0-p_e)/(1-p_e)
}
```
