## Supplementary material for "Worth the effort? A practical examination of random effects in hidden Markov models for animal telemetry data": Code: simulation.R

```
# requires momentuHMM >=1.5.2
library(momentuHMM)
library(TMB)
compile("HMM_recov.cpp")
source("simFunctions.R")

nbStates <- 2                         # number of states
M <- 5                                # number of individuals
obsVect <- c(30,50,70,150,250)        # number of observations per animal (length of obsVect must be a factor of M)
T_m <- rep(obsVect,M/length(obsVect))
maxMix <- 6                           # maximum number of finite mixtures to fit
retryFits <- 0                        # number of times to re-fit finite mixture models using random perturbations as initial values

## observation distribution parameters
dist <- "gamma" # gamma observation distribution
Par0 <- c(1,    # \mu^y_1
          10,   # \mu^y_2
          1,    # \sigma^y_1
          2)    # \sigma^y_2

## logit-scale state transition probability parameters (z)
z <- matrix(NA,M,nbStates*(nbStates-1))

## covariate effects
x_m <- rnorm(M)           # measurable covariate
mu_1 <- c(0,0)            # covariate effects for 1 -> 2 and 2 -> 1

#### continuous-valued random effects ####
mu_0 <- Sigma <- numeric(nbStates*(nbStates-1))
for(i in 1:(nbStates*(nbStates-1))){
  mu_0[i] <- 0          # mean for random effect
  Sigma[i] <- 0.416     # standard deviation for random effect
  z[,i] <- rnorm(M, mu_0[i] + x_m * mu_1[i], Sigma[i])
}
######### discrete random effects ########
#pie <- c(0.6,0.4)          # mixture probabilities 
#zMix1 <-  c(1.099,1.099)   # 1 -> 2 and 2 -> 1 for mixture 1
#zMix2 <-  c(-1.099,-1.099) # 1 -> 2 and 2 -> 1 for mixture 2
#gr <- apply(rmultinom(M,1,pie),2,which.max)
#for(i in 1:(nbStates*(nbStates-1))){
#  z[gr==1,i] <- zMix1[i] + x_m[gr==1] * mu_1[i]
#  z[gr==2,i] <- zMix2[i] + x_m[gr==2] * mu_1[i]
#}
##########################################


## generate data
exData <- momentuHMM::simData(nbAnimals=M,obsPerAnimal=as.list(T_m+1),nbStates=nbStates,dist=list(step=dist),
                  Par=list(step=Par0),beta=z,formula=~0+ID,covs=data.frame(cov=rep(x_m,times=T_m+1)),states=TRUE)
exData <- exData[-which(is.na(exData$step)),]
                  
covIndex <- c(1,cumsum(T_m)[1:(M-1)]+1) # first observation for each individual

## fit TMB models              
dyn.load(dynlib("HMM_recov"))
tmbOut <- list()
# no covariate
tmbOut$nocov <- fitTMB(exData$step, x_m, nbStates, nrow(exData), covIndex-1, FALSE, list(l_mu=log(Par0[1:2]),
                                                                              l_sd=log(Par0[3:4]),
                                                                              l_delta=0,
                                                                              mu_0=ifelse(rep(exists("mu_0"),nbStates*(nbStates-1)),mu_0,rep(0,nbStates*(nbStates-1))),
                                                                              mu_1=rep(0,nbStates*(nbStates-1)),
                                                                              l_sigma=ifelse(rep(exists("Sigma"),nbStates*(nbStates-1)),log(Sigma+1.e-3),rep(-3,nbStates*(nbStates-1))),
                                                                              epsilon=z))
tmbOut$nocov$kappa <- Kappa(tmbOut$states,exData$states) # proportion matching true states after accounting for chance

# with covariate
tmbOut$cov <- fitTMB(exData$step, x_m, nbStates, nrow(exData), covIndex-1, TRUE, list(l_mu=log(Par0[1:2]),
                                                                                             l_sd=log(Par0[3:4]),
                                                                                             l_delta=0,
                                                                                             mu_0=ifelse(rep(exists("mu_0"),nbStates*(nbStates-1)),mu_0,rep(0,nbStates*(nbStates-1))),
                                                                                             mu_1=mu_1,
                                                                                             l_sigma=ifelse(rep(exists("Sigma"),nbStates*(nbStates-1)),log(Sigma+1.e-3),rep(-3,nbStates*(nbStates-1))),
                                                                                             epsilon=z))
tmbOut$cov$kappa <- Kappa(tmbOut$cov$states,exData$states) # proportion matching true states after accounting for chance
dyn.unload(dynlib("HMM_recov"))


## fixed effects model                    
fixedOut<- momentuHMM::fitHMM(exData,nbStates=nbStates,dist=list(step=dist),
                              Par0=list(step=Par0),beta0=z,formula=~0+ID,modelName="fixed")
fixedOut$trProbs <- momentuHMM::getTrProbs(fixedOut,getCI=TRUE,covIndex=covIndex)
fixedOut$states <- momentuHMM::viterbi(fixedOut)
fixedOut$stateProbs <- momentuHMM::stateProbs(fixedOut)
fixedOut$kappa <- Kappa(fixedOut$states,exData$states) # proportion matching true states after accounting for chance

## BW random effects model
bwOut <- list()
# no covariate
bwOut$nocov <- momentuHMM::randomEffects(fixedOut,modelName="BWnocov")
bwOut$nocov$trProbs <- momentuHMM::getTrProbs(bwOut$nocov,getCI=TRUE, covIndex = covIndex)
bwOut$nocov$states <- momentuHMM::viterbi(bwOut$nocov)
bwOut$nocov$stateProbs <- momentuHMM::stateProbs(bwOut$nocov)
bwOut$nocov$kappa <- Kappa(bwOut$nocov$states,exData$states) # proportion matching true states after accounting for chance
# with covariate
bwOut$cov <- momentuHMM::randomEffects(fixedOut,Xformula=~cov,modelName="BWcov")
bwOut$cov$trProbs <- momentuHMM::getTrProbs(bwOut$cov,getCI=TRUE, covIndex = covIndex)
bwOut$cov$states <- momentuHMM::viterbi(bwOut$cov)
bwOut$cov$stateProbs <- momentuHMM::stateProbs(bwOut$cov)
bwOut$cov$kappa <- Kappa(bwOut$cov$states,exData$states) # proportion matching true states after accounting for chance

mixOut <- list(nocov=list(),cov=list())

## null model
# no covariate
mixOut$nocov[[1]] <- momentuHMM::fitHMM(exData,nbStates=nbStates,dist=list(step=dist),
                              Par0=list(step=Par0),beta0=matrix(apply(z,2,mean),1,2),modelName="nullnocov")
mixOut$nocov[[1]]$trProbs <- momentuHMM::getTrProbs(mixOut$nocov[[1]],getCI=TRUE,covIndex=covIndex)
mixOut$nocov[[1]]$states <- momentuHMM::viterbi(mixOut$nocov[[1]])
mixOut$nocov[[1]]$stateProbs <- momentuHMM::stateProbs(mixOut$nocov[[1]])
mixOut$nocov[[1]]$kappa <- Kappa(mixOut$nocov[[1]]$states,exData$states) # proportion matching true states after accounting for chance
# no covariate
mixOut$cov[[1]] <- momentuHMM::fitHMM(exData,nbStates=nbStates,dist=list(step=dist),formula=~cov,
                                    Par0=list(step=Par0),beta0=rbind(matrix(apply(z,2,mean),1,2),mu_1),modelName="nullcov")
mixOut$cov[[1]]$trProbs <- momentuHMM::getTrProbs(mixOut$cov[[1]],getCI=TRUE,covIndex=covIndex)
mixOut$cov[[1]]$states <- momentuHMM::viterbi(mixOut$cov[[1]])
mixOut$cov[[1]]$stateProbs <- momentuHMM::stateProbs(mixOut$cov[[1]])
mixOut$cov[[1]]$kappa <- Kappa(mixOut$cov[[1]]$states,exData$states) # proportion matching true states after accounting for chance

## finite mixture models
mixSet <- min(M-1,maxMix)
formula <- list(nocov=~1, cov=~cov)
for(i in c("nocov","cov")){

  for(j in 2:mixSet){
    
    ind <- which.min(unlist(lapply(mixOut[[i]][1:(j-1)],function(x) x$mod$minimum)))
    mixPar0 <- momentuHMM::getPar0(mixOut[[i]][[ind]],mixtures=j)
      
    mixOut[[i]][[j]] <- momentuHMM::fitHMM(exData,nbStates=nbStates,dist=list(step=dist),formula=formula[[i]],
                                        Par0=mixPar0$Par,beta0=mixPar0$beta,mixtures=j,modelName=paste0("mix",j,i),retryFits=retryFits)
    mixOut[[i]][[j]]$trProbs <- mixTrProbs(mixOut[[i]][[j]]) # individual-level state transition probability estimates
    if(i=="cov") mixOut[[i]][[j]]$mixCovs <- mixCovs(mixOut[[i]][[j]]) # population-level covariate estimates
    mixOut[[i]][[j]]$states <- momentuHMM::viterbi(mixOut[[i]][[j]])
    mixOut[[i]][[j]]$stateProbs <- momentuHMM::stateProbs(mixOut[[i]][[j]])
    mixOut[[i]][[j]]$kappa <- Kappa(mixOut[[i]][[j]]$states,exData$states) # proportion matching true states after accounting for chance
    mixOut[[i]][[j]]$mixtureProbs <- momentuHMM::mixtureProbs(mixOut[[i]][[j]])
  }
}

# BW cAICc model selection
cAIC <- AIC(fixedOut,!!!bwOut,!!!mixOut$nocov,!!!mixOut$cov,n=nrow(exData))
cAIC

# BW cBIC model selection
cBIC <- AIC(fixedOut,!!!bwOut,!!!mixOut$nocov,!!!mixOut$cov,k=log(nrow(exData)))
cBIC

# TMB mAICc model selection
tmp <- rbind(AIC(mixOut$nocov[[1]],!!!mixOut$nocov[2:mixSet],!!!mixOut$cov,n=nrow(exData)),data.frame(Model=c("tmbnocov","tmbcov"),AIC=c(tmbOut$nocov$aicc,tmbOut$cov$aicc)))
mAIC <- tmp[order(tmp$AIC),]
mAIC

# TMB mBIC model selection
tmp <- rbind(AIC(mixOut$nocov[[1]],!!!mixOut$nocov[2:mixSet],!!!mixOut$cov,k=log(nrow(exData))),data.frame(Model=c("tmbnocov","tmbcov"),AIC=c(tmbOut$nocov$bic,tmbOut$cov$bic)))
mBIC <- tmp[order(tmp$AIC),]
mBIC

# null/TMB likelihood ratio test
lrt(mixOut$nocov[[1]],tmbOut$nocov)
lrt(mixOut$cov[[1]],tmbOut$cov)
```
