## Supplementary material for "Worth the effort? A practical examination of random effects in hidden Markov models for animal telemetry data": Code: whaleFunctions.R

```
whaleTMB <- function(data, nbStates, nbSteps, aInd, par, DLL, ran) {
  dat <- list(data = data, nbStates=nbStates, nbSteps=nbSteps, aInd=aInd)
  # create function for automatic differentiation 
  # dat must contain all DATA required by hmm_tmb.cpp 
  # par must contain all PARAMETERS required by hmm_tmb.cpp 
  startTime <- proc.time()
  obj <- MakeADFun(data = dat, 
                   parameters = par, 
                   random = ran,
                   DLL = DLL, 
                   hessian = TRUE)
  mod <- do.call("optim", obj)
  sdr <- sdreport(obj,hessian.fixed=mod$hessian)
  endTime <- proc.time()-startTime
  rep <- summary(sdr, "report")
  
  trMat <- parSum(rep,which(rownames(rep)=="trMat"),nbAnimals,nbStates^2,FALSE,dimnames=list(paste0("ID",1:nbAnimals),paste0(rep(1:nbStates,nbStates)," -> ",rep(1:nbStates,each=nbStates)))) 
  trProbs <- lapply(trMat,function(x) array(t(x),dim=c(nbStates,nbStates,nbAnimals)))
  trProbs <- lapply(trProbs,function(x){rownames(x)<-colnames(x)<-paste0("state ",1:nbStates);x})
  trProbs$lower <- 1/(1+exp(-(log(trProbs$est/(1-trProbs$est))-qnorm(0.975)*(1/(trProbs$est-trProbs$est^2))*trProbs$se)))
  trProbs$upper <- 1/(1+exp(-(log(trProbs$est/(1-trProbs$est))+qnorm(0.975)*(1/(trProbs$est-trProbs$est^2))*trProbs$se)))
  
  fxd <- summary(sdr,"fixed")
  
  pMat <- matrix(paste0(rep(seq(1,nbStates),each=nbStates)," -> ",rep(1:nbStates,nbStates)),nbStates,nbStates)
  dia <- row(pMat) - col(pMat)
  betaNames <- c(pMat[dia!=0])
  
  CIbeta <- list()
  if(!is.null(ran)){
    ran <- summary(sdr, "random")
    CIbeta$beta <- parSum(ran,1:nrow(ran),nbAnimals,nbStates*(nbStates-1))
    betamle <- matrix(ran[,1],nbAnimals,nbStates*(nbStates-1),dimnames=list(paste0("ID",1:nbAnimals),betaNames))
  } else {
    CIbeta$beta <- parSum(fxd,which(rownames(fxd) %in% "epsilon"),nbAnimals,nbStates*(nbStates-1))
    betamle <- matrix(fxd[which(rownames(fxd) %in% "epsilon"),1],nbAnimals,nbStates*(nbStates-1),dimnames=list(paste0("ID",1:nbAnimals),betaNames),byrow=sum(rownames(fxd) %in% "epsilon")==(nbStates*(nbStates-1)))
  }
  
  CIreal <- list()
  CIreal$gamma <- lapply(trProbs,function(x) x[,,1])
  CIreal$delta <- parSum(rep,which(rownames(rep)=="delta"),nbAnimals,nbStates,FALSE,list(paste0("ID",1:nbAnimals),paste0("state ",1:nbStates)))
  
  varcomp <- vector('list',nbStates*(nbStates-1))
  for(i in 1:length(varcomp)){
    varcomp[[i]]$beta <- list()
    varcomp[[i]]$beta$sigma <- parSum(rep,which(rownames(rep)=="sigma")[i],1,1)
    cn<-exp(qnorm(0.975)*sqrt(log(1+(varcomp[[i]]$beta$sigma$se/(varcomp[[i]]$beta$sigma$est))^2)))
    varcomp[[i]]$beta$sigma$lower <- varcomp[[i]]$beta$sigma$est/cn
    varcomp[[i]]$beta$sigma$upper <- varcomp[[i]]$beta$sigma$est*cn
  }
  mle <- list(beta=betamle,
              delta=matrix(CIreal$delta$est,nbAnimals,nbStates,byrow=TRUE,dimnames = list(paste0("ID:",1:nbAnimals),paste0("state ",1:nbStates))))
  report <- obj$report(obj$env$last.par.best)
  return(list(mod=list(minimum = mod$value, estimate = mod$par, gradient = obj$gr(mod$par), hessian=mod$hessian, code=mod$convergence, iterations=mod$counts, elapsedTime=endTime[3], wpar=mod$par, Sigma=MASS::ginv(mod$hessian), message=mod$message),mle=mle,CIreal=CIreal,CIbeta=CIbeta,trProbs=trProbs,states=report$states,stateProbs=report$stateProbs,varcomp=varcomp,aic =  2 * length(mod[["par"]]) + 2 * mod[["value"]]))
}

parSum <- function(par,index,nr,nc,byrow=FALSE,dimnames=NULL){
  out <- list(est=matrix(par[index,1],nr,nc,byrow=byrow,dimnames=dimnames),se=matrix(par[index,2],nr,nc,byrow=byrow,dimnames=dimnames))
  out$lower <- out$est - qnorm(0.975)*out$se
  out$upper <- out$est + qnorm(0.975)*out$se
  out
}

lrt <- function(null,mod){
  nbStates <- length(null$stateNames)
  LR<- -2*(-null$mod$minimum- -mod$mod$minimum)
  X2 <- 0
  q <- 0
  qprime <- nbStates*(nbStates-1)
  for(r in 0:qprime){
    X2 <- X2 + 2^(-qprime)*choose(qprime,r)*dchisq(LR,r)
    q <- q + 2^(-qprime)*choose(qprime,r)*pchisq(LR,r)
  }
  p <- 1-q # if p<0.05, reject null of no individual-level variation
  cat(ifelse(p<0.05,"reject","failure to reject")," null at 0.05 significance level: p-value = ",p,", LRT statistic = ",LR,sep="")
}
```
